# Automating the Human Connectome Project’s Temporal ICA Pipeline

**DOI:** 10.1101/2024.01.15.574667

**Authors:** Chunhui Yang, Timothy S. Coalson, Stephen M. Smith, Jennifer S. Elam, David C. Van Essen, Matthew F. Glasser

## Abstract

Functional magnetic resonance imaging (fMRI) data are dominated by noise and artifacts, with only a small fraction of the variance relating to neural activity. Temporal independent component analysis (tICA) is a recently developed method that enables selective denoising of fMRI artifacts related to physiology such as respiration. However, an automated and easy to use pipeline for tICA has not previously been available; instead, two manual steps have been necessary: 1) setting the group spatial ICA dimensionality after MELODIC’s Incremental Group-PCA (MIGP) and 2) labeling tICA components as artifacts versus signals. Moreover, guidance has been lacking as to how many subjects and timepoints are needed to adequately re-estimate the temporal ICA decomposition and what alternatives are available for smaller groups or even individual subjects. Here, we introduce a nine-step fully automated tICA pipeline which removes global artifacts from fMRI dense timeseries after sICA+FIX cleaning and MSMAll alignment driven by functionally relevant areal features. Additionally, we have developed an automated “reclean” Pipeline for improved spatial ICA (sICA) artifact removal. Two major automated components of the pipeline are 1) an automatic group spatial ICA (sICA) dimensionality selection for MIGP data enabled by fitting multiple Wishart distributions; 2) a hierarchical classifier to distinguish group tICA signal components from artifactual components, equipped with a combination of handcrafted features from domain expert knowledge and latent features obtained via self-supervised learning on spatial maps. We demonstrate that the dimensionality estimated for the MIGP data from HCP Young Adult 3T and 7T datasets is comparable to previous manual tICA estimates, and that the group sICA decomposition is highly reproducible. We also show that the tICA classifier achieved over 0.98 Precision-Recall Area Under Curve (PR-AUC) and that the correctly classified components account for over 95% of the tICA-represented variance on multiple held-out evaluation datasets including the HCP-Young Adult, HCP-Aging and HCP-Development datasets under various settings. Our automated tICA pipeline is now available as part of the HCP pipelines, providing a powerful and user-friendly tool for the neuroimaging community.

## 1. Introduction

Functional magnetic resonance imaging (fMRI) is a powerful tool for mapping the brain’s functional organization noninvasively; however, fluctuations in fMRI data related to neural activity constitute only a small fraction of the overall variance (Glasser et al., 2018). The majority of fMRI fluctuations are instead due to structured artifacts such as head motion (Power et al., 2012; Power et al., 2014; Power et al., 2017; Satterthwaite et al., 2013; Yan et al., 2013), subject physiology such as respiration and cardiac pulsation (Birn et al., 2006; Chang & Glover, 2009; Golestani et al., 2015; Power et al., 2019; Shmueli et al., 2007), scanner artifacts (Glasser et al., 2018; Glasser et al., 2022), and unstructured (random) noise. Efforts to remove structured and unstructured noise from fMRI data have been ongoing since fMRI was developed (Birn et al., 2008; Jo et al., 2010; Murphy et al., 2013; Shmueli et al., 2007); however, many of the initially developed spatiotemporal filtering and nuisance regression methods do not selectively discriminate between true nuisance effects and correlated neural signal.

The Human Connectome Project (HCP) has helped popularize a data-driven approach based on independent components analysis (ICA) to selectively remove artifacts and nuisance signals, while retaining neurally related signals, to ensure that the resulting cleaned fMRI data is as close to the hypothetical neural signal as possible (i.e., avoiding both false positives and biases due to artifacts or nuisance signals, and also false negatives and biases due to neural signal removal). ICA (Beckmann & Smith, 2004; Hyvarinen, 1999) makes this possible by first decomposing the fMRI data into distinct components that are uncorrelated in either space or time using an unsupervised machine learning algorithm. Components are then classified into either signal or artifact classes by either an expert or a supervised machine learning algorithm. When cleaning fMRI data, neural signals are preserved by including them when computing regression coefficients (i.e., when regressing the data against the full set of components); the artifactual components are then selectively subtracted from the data (a non-aggressive approach, as opposed to the traditional aggressive approach of including only the nuisance components in the regression model, without consideration of any correlations with the neural signal).

When used on high spatial and temporal resolution fMRI data and precisely classified, spatial ICA (sICA) successfully removes the vast majority of spatially specific structured artifacts, the largest source of which is typically head motion (Salimi-Khorshidi et al., 2014). However, sICA is unable to separate global temporal fluctuations into spatially orthogonal signal and artifactual components because global spatial maps are inherently spatially correlated with each other and, therefore, end up mixed in with all of the components. Traditionally, “Global Signal Regression” has been widely used to remove global artifacts, but it is problematic because it removes both respiratory artifact and global or semi-global neural signals from the data, thereby distorting functional connectivity in network-specific ways (Glasser et al., 2018).

Recent studies show that temporal ICA (tICA) can segregate global and semi-global neural and nuisance signals into temporally orthogonal components (Glasser et al., 2018, 2019) and selectively remove the artifactual components without removing global and semi-global neural signals. tICA takes advantage of recent improvements in temporal resolution achieved by “multi-band’’ fMRI data acquisition and precise multi-modally driven cross-subject functional alignment in a combined cortical surface vertex and subcortical volume voxel CIFTI grayordinates space to enable group-level ICA decomposition. Despite the promising capabilities of tICA in fMRI data denoising, the previous lack of an automated pipeline, with multiple steps requiring manual intervention, presented barriers to widespread use of this approach. In particular, a simple automated approach is needed for selecting the spatial and temporal ICA dimensionalities based on the data, rather than selecting an arbitrary number of components manually. Also, manual labeling of tICA components as signal vs artifact is time-consuming and requires expert knowledge that is impractical for many investigators to learn. Moreover, tICA has thus far only been used on large-scale datasets with hundreds to thousands of participants, and it has not been clear how many subjects and timepoints are needed to avoid undesirable temporal ICA behaviors such as splitting components between subjects instead of separating artifacts from neural signals. Finally, we have recently achieved substantial improvements in spatial ICA component classification relative to the sICA-FIX method (Salimi-Khorshidi et al., 2014) by adding additional features, using multiple simple yet effective classifiers, and training on very large datasets.

In this paper, we present an automated tICA standardized pipeline. The proposed pipeline builds upon previous work (Glasser et al., 2018, 2019) and aims to automate the processing within a single user-friendly pipeline that contains nine steps: 1) MELODIC’s Incremental Group-PCA (MIGP; (Smith et al., 2014)) on sICA cleaned or recleaned data; 2) group-level sICA on MIGP after automated dimensionality estimation by fitting multiple Wishart distributions to the MIGP PCA series; 3) weighted regression into the individual dense timeseries using group sICA; 4) concatenation of the individual sICA timeseries; 5) group-level tICA on concatenated timeseries; 6) single regression into the individual dense timeseries using group tICA; 7) computation of various handcrafted features and latent spatial features from tICA components; 8) automatic classification of tICA components using a hierarchical classifier; 9) cleaning of the fMRI data. To adapt this tICA cleanup method to different study group sizes, the standardized pipeline is introduced with three tICA decomposition modes: i) computing a tICA decomposition from scratch (appropriate for large groups), ii) computing a tICA decomposition based on an initialized mixing matrix from a pre-decomposition result (appropriate for groups of modest size), and iii) directly using an existing tICA decomposition from another study, e.g., the young adult Human Connectome Project (HCP-YA), appropriate for single participants.

Automated ICA dimensionality estimation previously was typically performed in the volume by estimating the smoothness of the input fMRI timeseries (Beckmann & Smith, 2004). The Wishart-based approach incorporated here works in either volume or CIFTI grayordinate space based on fitting multiple Wishart distributions to a MIGP PCA series (Glasser et al, OHBM2021). We estimate the optimal number of Wishart distributions (WDs) needed to fit the HCP-Young Adult 3T and 7T datasets according to the resulting sICA estimated dimensionality, the quality of the fit of the Wishart distributions to the data, and the component reproducibility, which is applied to HCP-Young adult, HCP Aging and HCP Development official 3.0 release datasets.

The automated classification of tICA components using a hierarchical classifier involved both handcrafted features and latent spatial features learned from spatial autoencoders, and it showed overall good performance across a variety of HCP-style datasets. We also show that the hierarchical classifier is generalizable across a wide range of individuals, timepoints per subject, acquisition resolutions and field strengths, fMRI sessions, and tICA decomposition modes, suggesting that our pipeline is broadly applicable for automatically cleaning various types of HCP-style fMRI data from globally structured artifacts.

The tICA pipeline is publicly available as a part of the HCP Pipelines https://github.com/Washington-University/HCPpipelines.

## 2. Methods

### 2.1 Dataset information

#### Participants

We used resting-state fMRI and task-fMRI data from the 3T and 7T HCP Young Adult (HCP-YA) project (Van Essen et al., 2013) and from the two HCP Lifespan projects: HCP Aging (HCP-A) (Bookheimer et al., 2019) and HCP Development (HCP-D) (Somerville et al., 2018). All datasets had successfully completed 3T or 7T preprocessing by the HCP Pipelines (Glasser et al., 2013) plus spatial denoising using sICA+FIX (Salimi-Khorshidi et al., 2014) and areal-feature-based surface registration using MSMAll (Robinson et al., 2018; Robinson et al., 2014). This included 1071 HCP-YA participants (ages 22 – 35) scanned at 3T with 45 retest sessions, 175 HCP-YA participants scanned at 7T, 1798 3T HCP-A sessions (ages 36-100+), 1695 3T HCP-D sessions (ages 5 – 21). These data were acquired in accordance with the Washington University Institutional Review Board (Harms et al., 2018; Van Essen et al., 2013).

#### HCP-YA images

T1w and T2w HCP-YA structural scans were described previously (Glasser et al., 2022; Glasser et al., 2013; Ugurbil et al., 2013). The 3T resting state fMRI data were acquired with 2.0 mm isotropic resolution, TR=720 ms and 1200 frames (14.4mins) for each of 4 runs. Two runs each were acquired with reversed phase encoding directions (RL or LR) with the order counterbalanced across each of two scan sessions (Smith et al., 2013) for a total of 4800 frames of resting state per subject along with matching phase reversed spin echo fMRI scans for distortion correction. The 3T task fMRI data were acquired using identical pulse sequence settings while participants performed 7 tasks (working memory, gambling, motor, language, social cognition, relational processing, and emotion processing; (Barch et al., 2013)) with runs lasting between 2 and 5 min (176-405 frames) and totaling 22.6min (1884 frames) in session 1 and 24min (1996 frames) in session 2 for a total of 3880 frames of task fMRI per subject. Each task scan comprised a pair of runs with reversed phase encoding directions, first RL then LR. Task runs were halted if a subject stopped performing the task. The 7T resting-state fMRI data were acquired with 1.6mm isotropic resolution, TR=1.0s and a maximum of 3600 frames for 4 runs. The 7T movie and retinotopy fMRI data were acquired using identical pulse sequence settings as the resting-state fMRI; they included a total of 3655 frames for the movies (4 runs) and 1800 frames for retinotopy (6 runs).

#### HCP-A and HCP-D images

The acquisition of the HCP-A and HCP-D structural scans has been described previously (Glasser et al., 2022; Harms et al., 2018). Resting-state fMRI scans were acquired with 2.0 mm isotropic resolution, TR=800ms and 488 frames (6.5min) for each of 4 runs. Two runs each with anterior-to-posterior (AP) and posterior-to-anterior (PA) phase encoding for a total of 1952 frames of resting state were acquired per subject with matching phase reversed spin echo fMRI data for distortion correction. Task fMRI data were acquired with identical pulse sequence settings while subjects performed 3 tasks per project (VisMotor, Go/NoGo, and FaceName for HCP-A subjects; (Bookheimer et al., 2019); Guessing, Go/NoGo, and Emotion for HCP-D subjects; (Somerville et al., 2018)) with runs lasting between 2 and 5 min (178-345 frames), and a total 2721 frames for HCP-A, 3200 frames for HCP-D (2246 frames for age 5-7). HCP-D included two runs of the first two tasks, acquired with opposite phase encoding polarity, and a single run for the last; HCP-A included only a single run for all three tasks. In addition, Arterial Spin Labeling (ASL) scans were acquired at 2.5mm isotropic resolution with multiple labeling delays to estimate Cerebral Blood Flow (CBF) and Arterial Transit Time (ATT) (Harms et al., 2018).

#### Image preprocessing

All datasets were preprocessed using the HCP Pipelines (Glasser et al., 2013) which includes three structural preprocessing pipelines, two functional preprocessing pipelines, multi-run spatial ICA+FIX (Glasser et al., 2018; Salimi-Khorshidi et al., 2014) fMRI data denoising pipelines and the multi-modal ‘MSMAll’ areal-feature-based surface registration (Robinson et al., 2018; Robinson et al., 2014). Additionally, we applied a secondary step of a spatial Independent Component Analysis (sICA) reclean process to all datasets (Refer to Supplementary Methods), which improves the sICA+FIX component classification by adding additional features and training data. Subsequently, we retrained Multi-layer Perceptron and Random Forest classifiers on the sICA components having high FIX confidence, which included both raw FIX features and the newly introduced features. This was a semi-automated approach in which we manually checked components where different classifiers generated conflicting classifications. (See Supplementary Methods for detailed information about the reclean process and the automated reclean pipeline). Importantly, while the reclean process is recommended to further reduce spatially specific artifacts following sICA+FIX, we intentionally included datasets without the reclean process to demonstrate the robustness of this tICA pipeline. Consequently, the preprocessing results yielded MSMAll-registered grayordinate-wise (dense) or voxel-wise sICA+FIX cleaned/recleaned timeseries data for each subject.

### 2.2 The tICA pipeline

#### tICA pipeline

The HCP-style tICA cleanup introduced by (Glasser et al., 2018) demonstrated tICA’s effectiveness in selectively separating temporally global or semi-global artifacts from neural signals. To standardize the tICA cleanup process and enable widespread access for the community, we implemented a 9-step pipeline that uses group PCA and spatial ICA to perform a consistent spatial dimensionality reduction across subjects, produce temporally concatenated sICA timeseries across all subjects, decompose these timeseries into temporally independent components including neural signals, physiology artifacts, and other artifacts, classify these components automatically, and then regress the artifactual components out of each subject’s data in a non-aggressive way (similar to sICA-FIX). This tICA pipeline provides three estimation modes for tICA decomposition to encompass a wide range of study group sizes:

- ESTIMATE: tICA decomposition is computed from scratch if a sufficient number of subjects and timepoints are collected for a group decomposition initialized from a random matrix.
- INITIALIZE: a previously computed group sICA decomposition and tICA mixing matrix are used to initialize a final tICA decomposition.
- USE: a precomputed group sICA and a tICA mixing matrix are directly used for decomposition without modification.

Each of the nine steps are sequentially illustrated in Fig 1 with detailed information in the following sections.

**Figure 1.**
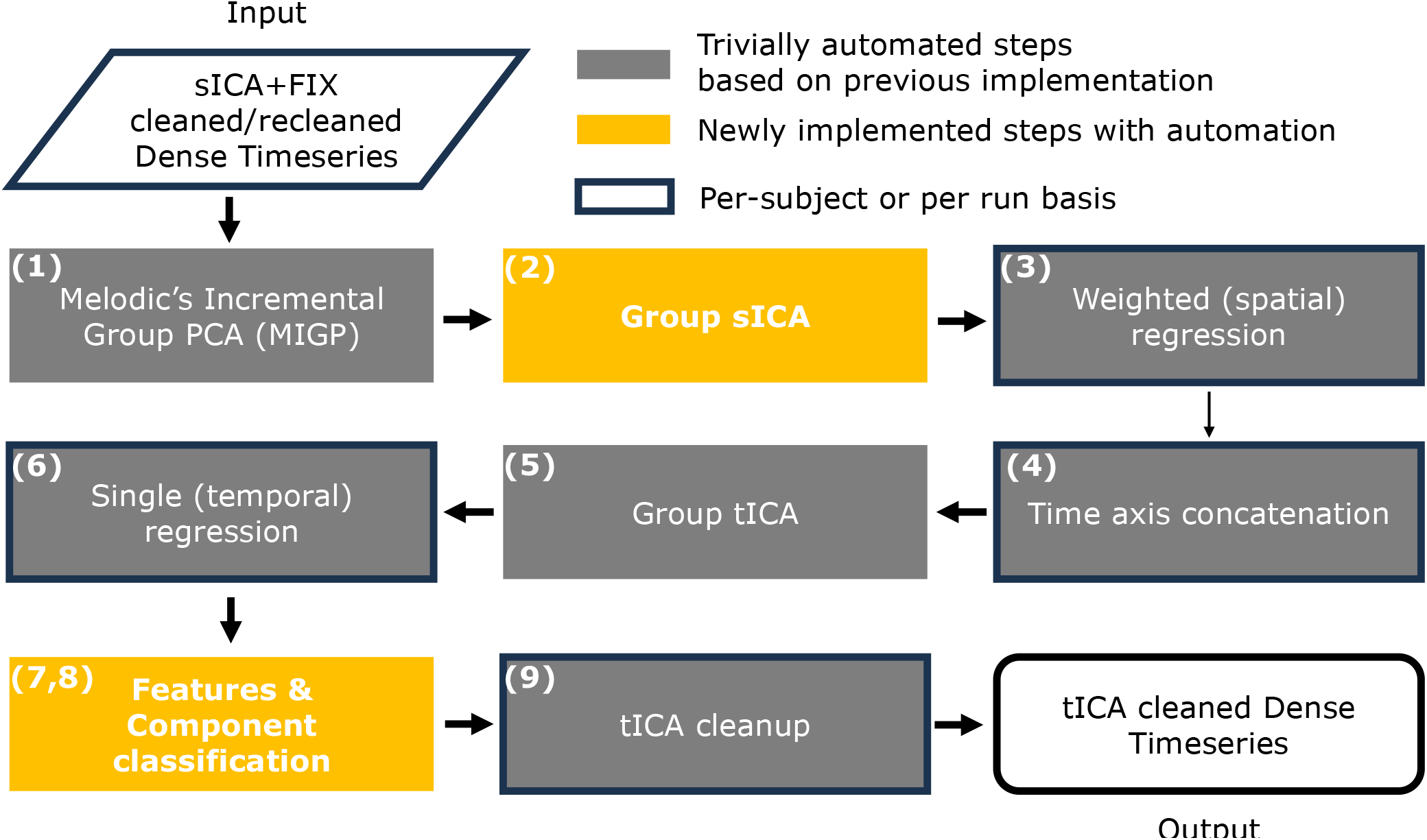
shows an overview of the automated temporal ICA (tICA) pipeline for the HCP projects from preprocessed sICA+FIX cleaned or recleaned dense timeseries to the final tICA cleaned dense timeseries. Yellow boxes indicate steps that are newly automated in this study.

#### 2.2.1 Step 1. Creation of a compact group fMRI dataset using MELODIC’s Incremental Group PCA (MIGP)

ICA requires PCA to be performed before dimensionality estimation, and MIGP enables generation of a compact group PCA even when the sum of all subjects’ number of timepoints is very large (Smith et al., 2014). MIGP incrementally estimates the PCA of the dense timeseries for a group of subjects in a sequential manner (reducing memory requirements without disrupting the component amplitudes). MIGP aims to capture the structured variance from all subjects (while not excessively modelling the unstructured noise), reducing the PCA dimensionality to a predetermined number. The inputs to MIGP consist of each subject’s sICA+FIX cleaned or recleaned MSMAll aligned dense timeseries and the maps needed to variance normalize these timeseries according to the distribution of unstructured noise. The outputs of MIGP are a dense PCA series representing the most structured signals common to the group, and a group-average spatial map of unstructured noise variance that is used to convert group spatial ICA components from intensity normalized maps to variance normalized maps.

#### 2.2.2 Step 2. Group spatial ICA decomposition with dimensionality estimation using multiple Wishart distributions (GroupSICA)

Group spatial Independent Component Analysis (ICA) serves a dual purpose: it ensures a consistently reduced spatial dimensionality across all subjects in a group and identifies shared spatial patterns among subjects, thereby enhancing the stability of subsequent group temporal ICA operations. Prior to running group spatial ICA on the PCA dense timeseries obtained through the MIGP approach, it is essential to determine the number of components for decomposition directly from the data. To estimate the structured dimensionality of the data, we first fit a Wishart Distribution (WD; (Wishart, 1928)) to the smallest component amplitudes, which represent unstructured noise. Because the MIGP PCA series has undergone extensive preprocessing (MRI image reconstruction, MRI preprocessing, resampling to MNI space, mapping from volume to surface, and the MIGP reconstruction, at least some of which cause different amounts of data mixing at different locations), it does not exactly fit a single Wishart distribution. Accordingly, multiple Wishart distributions are first fit to the data, and then a single Wishart distribution is fit to the sum of the multiple Wishart distributions. This single Wishart distribution is then subtracted from all of the component amplitudes, and the Bayesian model selection using Laplace approximation (Beckmann & Smith, 2004; Minka, 2000) is used on the modified amplitudes that are greater than zero to estimate the data dimensionality (i.e., the number of components that are substantially above the modeled noise amplitude, and thus represent structured variance of interest).

For MIGP in CIFTI space for most HCP-style datasets (2721 to 4800 PCA components), we found that six Wishart distributions produce ICA dimensionalities that were comparable to those previously produced by an alternative, more manual method of dimensionality estimation that focused on producing the largest number of reproducible temporal ICA components ((Glasser et al., 2018); d=84 for 3T resting state and d=70 for 3T task fMRI), neither oversplitting nor undersplitting the data. The shorter HCP Young Adult 7T retinotopy dataset (1800 PCA components) uses a value of five to produce a comparable dimensionality to the other datasets. A value of 4 clearly includes substantial unmodelled unstructured noise autocorrelation from the MIGP components and as a result produces dimensionality estimates that are too large. A value of 7 clearly produces too few components relative to other more manual methods of dimensionality estimation.

Once the ICA dimensionality is determined, ICA is performed on the MIGP data after Wishart Filtering (reconstructing the MIGP PCA series after subtracting the single Wishart Distribution from the amplitudes as in (Glasser et al., 2016) so as to further improve the ICA decomposition reproducibility and focus the ICA decomposition on structured real signals rather than the autocorrelations in the otherwise unstructured noise). The ICASSO approach (Himberg & Hyvarinen, 2003) is used to estimate the optimal mixing matrix in a reproducible way in 100 resampling cycles. The ICA mixing matrix describes the relationship between the input channels (in this case the timeseries associated with sICA components) and the estimated ICA sources (the tICA component timeseries). This mixing matrix is used to initialize a final FastICA (the algorithm used for spatial and temporal ICA decomposition; (Hyvärinen & Oja, 2000)) run, which generates a reproducible ICA estimation. In addition to the group spatial ICA components obtained under the estimated dimensionality, fixed low-dimensional group ICA decompositions (ranging from seven to twenty-one) are generated for the purpose of weighting the spatial regression towards the parts of the brain that are well aligned functionally across subjects, as described in (Glasser et al., 2016).

#### 2.2.3 Steps 3 & 4. Weighted spatial multiple regression (indProjSICA) and concatenation of individual’s sICA dense timeseries (ConcatGroupSICA)

Weighted spatial multiple regression (WR), a variant of dual regression proposed in (Glasser et al., 2016), addresses misalignments that remain after MSMAll areal-feature-based registration and produces individualized sICA components for each subject’s fMRI data while ensuring that each component reflects functionally corresponding regions across individuals. This technique employs spatial regression, with weighting based on vertex area and alignment quality as derived from low-dimensional independent component (IC) maps, which are less vulnerable to the effects of misalignments than high dimensional maps. The vertex area map represents the vertex areas of the individual subject’s midthickness surface in native volume space and avoids the distortion of the MNI space midthickness surface. The low-dimensional independent component maps serve as an alignment check, generated from weighted dual regressions of low ICA dimensionalities ranging from 7 to 21, weighted solely by the vertex area map, and where the individualized low-dimensional map does not match the group map, we assume the functional alignment of that subject is locally not as good.

In the first round, group spatial ICA components are regressed into the dense timeseries of individual subjects to obtain individual sICA timecourses. These are weighted based on the element-wise product of the vertex area map and the alignment map. Subsequently, these timecourses are temporally regressed into the grayordinate individual subject timeseries, resulting in individual subject sICA component spatial maps. In the second round, the first-round individual spatial maps undergo spatial multiple regression into the individual subject dense timeseries, with weighting solely from the vertex area map to produce refined individual subject timeseries, which are then temporally multiple regressed into the individual subject CIFTI and volume timeseries to generate refined individual subject component spatial maps in both CIFTI and volume spaces.

To facilitate the temporal ICA decomposition, the cross-subject individual sICA timecourses are concatenated along the time axis. In cases where a subject’s sICA timecourse contains fewer timepoints than the expected number across the group, zero values are padded to the end of its time axis. This ensures that the group concatenated time dimension is the product of the total number of subjects in the group and the expected timepoint number for each subject, making recordkeeping easier.

The outputs of these two steps encompass the following: (1) for each subject, a timecourse corresponding to each group sICA component, (2) a subject-specific CIFTI grayordinates spatial map for each sICA component, (3) a subject-specific volume spatial map for each sICA component, (4) the group average of the subject-specific grayordinate-based spatial maps and the subject-specific volume-based spatial maps (2 and 3 above), and (5) the temporal concatenation of the individual sICA timecourses (1) across all subjects, which serves as the input for group tICA.

#### 2.2.4 Single-subject effects in tICA decomposition using FastICA

Temporal ICA is capable of identifying temporally independent components (e.g., respiration) on a large-scale group dataset as shown previously (Glasser et al., 2018, 2019). However, when the number of subjects or the total number of concatenated timepoints is relatively small, subject-specific dominance within some components can emerge. The affected components exhibit a disproportionately high variance within a single subject or a small subset of subjects while demonstrating minimal variance across the remainder of the group. We define these tICA components dominated by individual subjects as single-subject components, and we incorporate two alternative decomposition modes to mitigate their impact (as they may not accurately separate somewhat temporally correlated signal and artifacts).

This phenomenon is largely attributed to the intrinsic characteristics of the FastICA algorithm, which seeks to identify statistically independent sources of signal based on their non-Gaussianity. In essence, FastICA favors super-Gaussian distributions (characterized by greater kurtosis, i.e., heavier tails, and sharper peaks compared to a Gaussian distribution). With a large pool of subjects, the individual differences, or unique sources of signal variance, tend to average out. This allows the algorithm to extract independent components that are consistent across the group, thereby revealing shared temporal patterns. However, if the subject pool is more modest in size, these unique sources of signal variance (e.g., the unique T2* weighting across each subject’s brain determined by their head orientation in the scanner and unique folding pattern) can have exaggerated effects. These effects are stronger at higher field strength (e.g., 7T). The presence of subject-specific spatio-temporal patterns results in a super-Gaussian distribution, which FastICA may identify as an independent component. As a result, directly estimating tICA decomposition on a smaller subject group can lead to components that are specific to particular subjects but represent combinations of different signals or between signal and artifact. This can reduce denoising efficiency and the generalizability of the group decomposition, thereby negatively impacting the interpretability of the results.

While conventional FastICA typically starts with a random initial mixing matrix, we incorporated an initialization mode that allows users to launch the algorithm with a precomputed mixing matrix that is closer to the ideal mixing matrix. This can be particularly advantageous in scenarios where prior knowledge on a large-scale dataset and a good estimate of the mixing matrix is available, significantly accelerating the convergence of the algorithm and also reducing the single subject effect in small group sizes. (See evidence in Table S.1)

To make the tICA pipeline applicable to HCP-style datasets spanning a wide range of subject counts and fMRI frames acquired, we do not require that a tICA decomposition be estimated from scratch (using the ESTIMATE mode). Instead, we added two modes based on a pre-estimated large scale tICA mixing matrix (e.g., from HCP Young Adult or HCP Lifespan): either initializing the tICA decomposition from the pre-estimated mixing matrix (INITIALIZE mode), enabling further optimization for each dataset, or directly using the mixing matrix without any re-estimation (USE mode) (see below for details).

#### 2.2.5 Step 5. Group temporal ICA decomposition (ComputeGroupTICA)

Each run of the concatenated individual subject sICA timecourses is first variance normalized to ensure equal contribution from all runs in the subsequent decomposition. Group tICA is then performed on the run-wise variance-normalized concatenated sICA timecourses using the FastICA algorithm. The dimensionality of the tICA is chosen to match the previously estimated sICA dimension in Step 2. For each tICA decomposition, the “tanh” contrast function and a symmetrical approach are employed, with a maximum of 1000 iterations used for ICA estimation.

In the ESTIMATE mode, where the group tICA mixing matrix is estimated from scratch, each of the first four rounds of decompositions uses ICASSO with 100 estimations. Because temporal ICA is less robust than spatial ICA (in this context, with weaker non-Gaussianity in time than in space), we use multiple rounds to make convergence of a reproducible decomposition more likely. The first round of decomposition uses both data bootstrapping and randomization of initial conditions. The subsequent three decompositions use only data bootstrapping, with each one initializing the estimation process using the previously determined best estimated mixing matrix. Finally, a single-pass ICA decomposition is performed on the data matrix, with the last best estimate of the mixing matrix serving as the initialization. This generates the final group tICA components and the tICA mixing matrix. For clarity, we define the group tICA components as the resulting unmixed timecourses with the dimension in the time axis equal to the product of the number of subjects and the expected number of timepoints for each subject, representing the underlying temporally independent latent sources.

In the INITIALIZE mode, where the group tICA has an assigned pre-computed mixing matrix as the initialization, the process begins from the second round of decomposition in the ESTIMATE mode, using the given initialization, while the subsequent steps remain the same as in the ESTIMATE mode.

In the USE mode, no new ICA decomposition is performed, and the mixing matrix provided is directly applied to unmix the data matrix to generate group tICA components. This mode can directly use the component classification results from the precomputed datasets and be applied to single subjects.

Subsequently, the group-level temporal ICA spatial maps in grayordinate and volume space are generated by matrix multiplying the corresponding group-averaged spatial ICA maps with the resulting tICA mixing matrix (Glasser et al., 2018). The group tICA components are then ordered based on the descending order of the standard deviation of the unnormalized group tICA components, with components that explain more variance appearing earlier. Importantly, the sign of ICA components is arbitrary, because flipping the sign of the mixing matrix and component matrix at the same time results in the same observation. By convention, the final group tICA components’ maximum spatial map value is set to be positive.

To aid visual inspection and component classification, the group-averaged tICA components (timecourses) and power spectra are generated.

#### 2.2.6 Step 6. Single regression (indProjTICA)

The group tICA temporal components are de-concatenated to produce each individual subject’s tICA temporal components, which are then regressed into each individual subject’s dense timeseries in either volume or grayordinate space to produce individual tICA spatial maps. For each group tICA component, the individual for each tICA component that has the largest variance explained by that component is defined as the strongest subject for that group tICA component.

#### 2.2.7 Step 7 & 8. Feature generation (ComputeTICAFeatures) and component classification (ClassifyTICA)

After tICA decomposition, it is crucial to determine which components are artifacts to remove and which are signals to retain. The automated classifier described here and the process used to train the classifier are based on diverse types of information, including handcrafted features from volume, surface, timeseries, power spectrum, and greyplots - a 2D data representation of fMRI data with time along the x axis and space (grayordinates or parcels; parcels are used in this study) along the y axis, and fMRI image intensity rendered in greyscale (Glasser et al., 2018, 2019; Power et al., 2017) at both group and individual levels. In addition, spatial latent features from volume and surface autoencoders are used in a self-training schema. Details about the handcrafted features, the spatial latent features and the classifier are described below (section 2.3). The output of this step is a list of artifactual components for cleanup.

#### 2.2.8 Step 9. tICA cleanup

The individual subjects’ spatial beta maps are produced from the single regression in Step 6. Subsequently, the group tICA components classified as artifacts for removal and their associated beta maps are multiplied together to determine the portion of the dense timeseries that can be best attributed to the artifactual components in a non-aggressive fashion. Finally, this artifactual dense timeseries is subtracted from the sICA+FIX cleaned or recleaned timeseries data, resulting in the generation of the tICA cleaned dense timeseries data.

### 2.3 Automated tICA component classification

The tICA classifier is based on handcrafted features and latent spatial features. These features together serve as discriminative cues to distinguish signal from artifactual components. A hierarchical classifier is then trained on the features and manual labels to predict temporal ICA components at the group level. To validate its effectiveness, we evaluated its performance on eleven held-out datasets, demonstrating its potential as a solution for automated tICA component classification. We start this section by introducing the datasets involved in the training and evaluation process, each of which underwent the full tICA pipeline described above in Steps 1-6.

#### 2.3.1 Splitting of HCP datasets into study datasets

##### Train-valid-eval datasets

The HCP-YA dataset was split into one training dataset and one evaluation dataset, each with 535 subjects for 3T scans and 87 subjects for 7T scans, and with no family relationships across datasets (i.e., no subject in one dataset was related to any subject in the other dataset; all related subjects were in the same dataset to prevent data leakage). Two subgroups were generated from HCP-A: a training dataset and a testing dataset (Aging-s in Table 1), each with 617 subjects who shared no family relationship. HCP-D was separated into seven datasets: a main training dataset and a main evaluation dataset, each with 314 subjects; three independent training datasets with 60, 100, 160 subjects and two independent evaluation datasets with 60 subjects and 100 subjects, to represent studies having relatively few subjects. No family relationship is shared across any of the seven datasets. Two of the four independent evaluation datasets from HCP-D went through both the ESTIMATE mode and the INTIALIZE mode with the HCP-YA 3T resting state evaluation dataset as the precomputed decomposition. All remaining datasets were computed with ESTIMATE mode only. Details of the training and evaluation datasets are shown in Table 1, including the percentage of variance explained by signal components, artifactual components and weak components. The majority of the variance is contributed by signal components (67%∼93%); the components that show a weak intensity profile and are borderline to classify are defined as weak components and explain only a small portion of the total variance (0%∼28%).

**Table 1.**
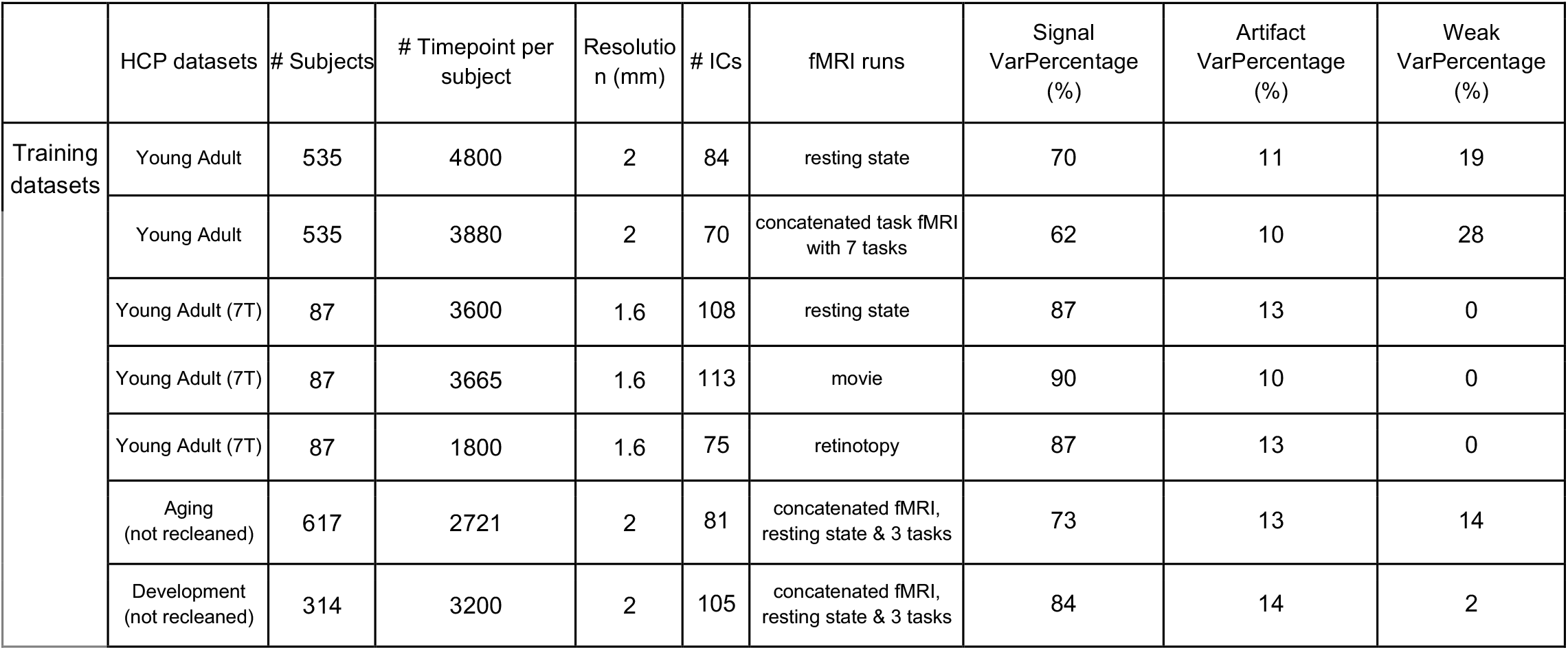
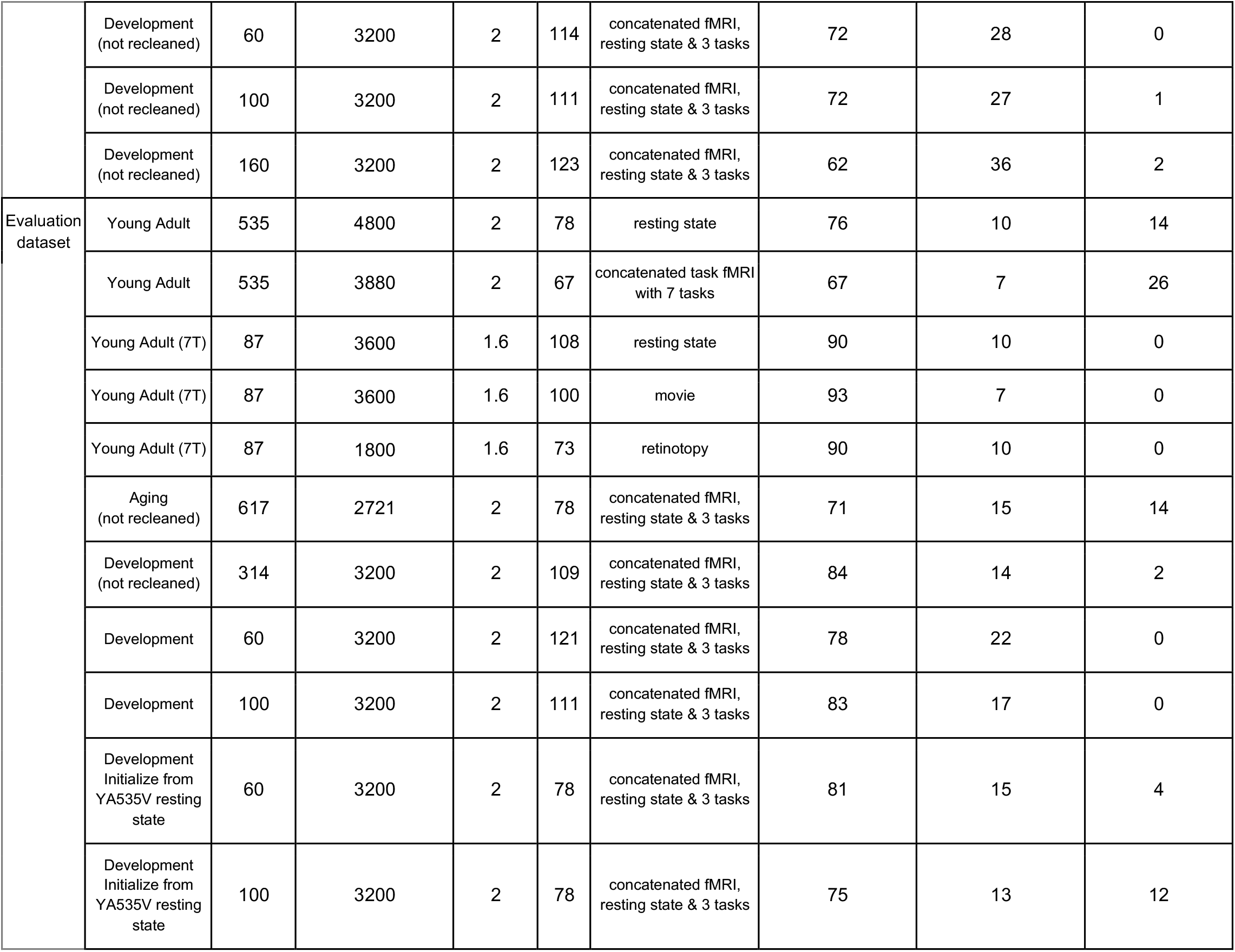
A summary of the datasets analyzed in this study. Each dataset was processed using the tICA pipeline up to the feature generation step. Including a variety of dataset properties helps to generalize the trained model across a range of subject counts (single-subject effect changes), acquisition resolutions, timepoints per subject, fMRI sessions and tICA modes. If a dataset did not undergo the sICA reclean process, “not recleaned” is included in the dataset column, otherwise, it has been recleaned. YA535V: The Young Adult 3T evaluation tICA dataset with 535 subjects. VarPercentage: the percentage of variance explained by certain type of components.

#### 2.3.2 tICA component categories and manual classification

The classification of group tICA components into binary labels, namely signal or artifact, is a crucial step for enabling tICA cleanup, similar to spatial ICA. However, during our manual classification process, we identified a need for more detailed categorization, as we encountered uncertain or disputed categories that could not be convincingly classified as signal or artifact. Here, we summarize these categories for users who opt to manually classify their group tICA components.

##### Clear Signal

This category comprises the majority of group tICA components representing neural signal. Determination of clear signal components can be based on 1) similarity to known networks or task activation patterns, 2) alignment with known areal boundaries such as those in HCP-MMP1.0 (Glasser et al., 2016), and 3) correspondence with established somatotopic or retinotopic topographic organization.

##### Special Signal

Special signal components exhibit spatial maps that respect brain areal and functional network boundaries but display lagged correlation with the respiratory belt envelope signal (Power et al., 2020) (but with shorter response functions that are typical of neural activity rather than the longer respiratory response function). Some investigators might wish to remove such components whereas others would not (Glasser et al., 2018, 2019; Power, 2019). Recently, such components have been implicated in travelling waves across the cortex (Raut et al., 2021). Additional analysis and discussion of these components is warranted in the future.

##### Respiration Artifact

Components falling into this category typically exhibit a globally positive spatial map (with the exception of regions with very short T2*) across the gray matter in both resting state and task-based fMRI or display a spatial pattern that aligns with known vascular delay maps (e.g., ATT as measured with ASL), suggesting respiratory-related nuisance signal. The vascular delay maps represent the differential arrival times of a bolus of blood with differential pCO2 content than the prior blood leading to lags in the global respiratory effect that respect vascular territories (particularly core versus watershed and anterior circulation faster than posterior circulation).

If a group tICA component is dominated by one subject or a subset of subjects, there tend to be outlier variances among the group of the individual subject’s tICA components, resulting in a distribution of variances with a heavy tail compared to a normal distribution. The variance distribution that includes outlier variances can be characterized by high kurtosis values, which is useful to find single subject-related patterns.

##### Single Subject Signal

If a group tICA component is single subject dominated and the dominating subject’s spatial map represents neural signals similar to the Clear Signal category, we categorize it as Single Subject Signal.

##### Single Subject Respiration

If a group tICA component is single subject dominated, with its dominating individual component(s) displaying relatively high covariance with the global signal (defined as the average intensity across the grayordinate space) compared to the rest of the subjects’ tICA components, it is defined as single subject respiratory-related artifact. Such components can be characterized by high kurtosis values on the covariances distribution between the individual components and the global signal.

##### Spatially Specific Artifact

This category encompasses remaining spatially specific artifacts after sICA+FIX and recleaning. It includes 1) weak but consistent artifact patterns, 2) volume distortion components resulting from severe head motion in HCP 7T datasets leading to incomplete distortion correction and thereby misalignment of greymatter throughout an fMRI run (see supplementary Fig S11), 3) clear head motion artifacts (e.g., alternating positive and negative intensity around the brain boundary) in the dominating subject’s tICA spatial map, and 4) unexpectedly high intensity in the distal brainstem.

##### Weak Signal

Weak components are identified by a relatively low maximum intensity value in the group-averaged power spectra. If a weak component respects known areal boundaries or regions of high signal intensity in the cortical surface maps, it is classified as a weak signal component.

##### Weak Subcortical

The weak subcortical category includes weak components that display patches largely restricted to well-defined subcortical areas.

##### Weak Other

The weak other category encompasses the remaining weak components not identified as weak signal or weak subcortical.

Determining the signal or artifact category for weak components poses challenges. However, since weak components (weak signal, weak subcortical, and weak other) typically account for only a small percentage of the variance (see Table 1), they can be either categorized as signal to prevent potential signal loss or classified as artifact to ensure the removal of ambiguous data variance.

During the development of the automated classifier, we followed the same category-to-label mapping as in the official release for the HCP-YA and HCP-Lifespan tICA cleaned dataset, which assigned Clear Signal, Single Subject Signal, Weak Signal, Weak Subcortical, Weak Other into the signal category, and Respiration Artifact, Single Subject Respiration, Spatially Specific Artifact into the artifact category, to minimize potential signal loss.

##### Manual classification

Each of the 1,985 tICA components referred to in Table 1 was first manually classified by CY based on domain knowledge and component similarities compared with the tICA group components from the official release of HCP-YA, HCP-A, HCP-D. Any component that was uncertain by CY was further evaluated by MFG to determine the final category.

#### 2.3.3 Handcrafted features

Handcrafted features were designed based on domain knowledge derived from multiple data types, including volume and cortical surface maps, timeseries, power spectra, and spatio-temporal ‘greyplot’ representations (Glasser et al., 2018, 2019; Power et al., 2017). Fig 2 shows a schematization of handcrafted features.

**Figure 2.**
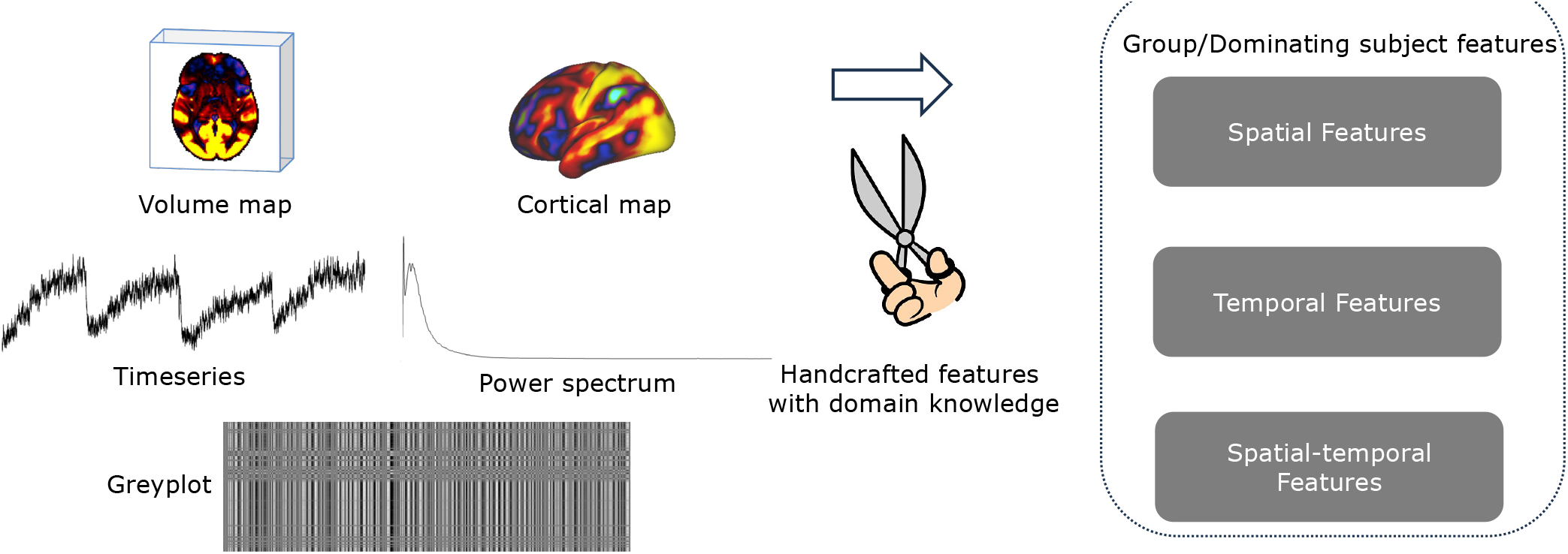
shows a high-level framework of handcrafted features, which fall into three main categories: spatial features, temporal features and spatio-temporal features at both the group level and individual level.

##### Outlier identification

Similar to the manual classification process, the first step in classifying a group tICA component is to identify whether it is dominated by one or a few individual subjects. The appearance of variance outliers across the individual subject tICA components corresponding to each group tICA component indicates that a group tICA component is single subject dominated (typically with one or two outlier subjects). To quantify the degree of “outlierness” in a set of variances ***a***(usually a vector for each individual tICA component), we used an outlier effect indicator ***O***:

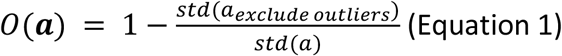

Each outlier is identified if the value of ***O*** meets two criteria: (i) the value is more than three scaled median absolute deviations (MAD) away from the median (MATLAB command: *isoutlier*); (ii) the value is outside of the 5 to 95 percentile range. This combination helps distinguish true outliers from extreme values or random fluctuations, providing a more robust approach for outlier detection. If there is a strong outlier effect (e.g., several individuals’ tICA components exhibit extremely high variances compared with the rest), the outlier effect indicator tends to have a value close to 1 because the standard deviation of the remaining variances is much smaller than the standard deviation of all the variances.

Similarly, such a function can be used to quantify the “outlierness” in covariances between a set of individuals’ tICA components and their corresponding global signal. This outlier indicator is also used to quantify outlier effects in spatially masked regions.

##### Spatial features

The tICA spatial maps, both on the cortical surface and in the volume, provide crucial information for the classification of components. Components belonging to the signal and artifact categories exhibit distinct characteristics across various tissue types, including cortical gray matter, white matter, subcortical gray matter, and cerebrospinal fluid (CSF). BOLD signals are primarily expected to be detected in gray matter regions. Volume maps of artifactual tICA components may exhibit changes in subcortical regions, white matter, and CSF. Spatially restricted masks are utilized to identify regional patterns, such as the left and right cerebellum, as well as left and right gray matter regions excluding the cerebellar areas. Nine vascular territories are masked to extract information related to vascular delay components. To map the mean transit time of blood in the brain, high-resolution arterial spin labeling data was carefully preprocessed to remove distortions, intensity biases, and motion artifacts (see Fig S13A; Kirk et al., In prep). The resulting data was then precisely mapped onto individual cortical surfaces, aligned using MSMAll across subjects, and averaged. Spatially specific artifact resulting from head motion can be identified using brain boundary masks eroded from 2 to 5 times the fMRI resolution (e.g., 2mm, 1.6mm). Mean, variance, and outlier effect indicators derived from these masks in the volume or CIFTI spaces are computed as spatial features.

To identify artifacts associated with large vessels, a vessel probability atlas (Mouches & Forkert, 2019) is incorporated with the tICA volume maps. Features based on the vessel probability atlas include the sum of signal intensity multiplied by the per-voxel probability, the sum of absolute signal intensity multiplied by the per-voxel probability, and the outlier effect indicator within vessel regions (non-zero regions in the vessel probability map).

In sagittal views of tICA volume maps, components affected by multiband-related autocorrelation exhibit alternating positive and negative stripes along the slice direction—a consistent spatial pattern primarily observed in 3T HCP data, which conveniently has a consistent “location” when transformed back to k-space, for a given multiband factor and number of slices. A template was created using the training dataset by averaging the magnitude of the k-space data from hand-selected “stripey” components and subsequently thresholding the averaged k-space data to generate a binary mask. The correlation coefficient between the template and the k-space data is computed for each group tICA volume map to detect artifacts associated with multiple bands.

All predefined template masks in the volume space (e.g., vessel probability map, multiband k-space template, vascular territories) are based on a voxel size of 1.6mm isotropic. For datasets represented at a different spatial resolution, features based on such masks are computed by first resampling the volume maps to 1.6mm isotropic voxels.

The updated globality index serves as a component-wise measure on tICA cortical surface maps, aiding in the detection of global respiratory artifacts. It is computed based on the tICA surface maps in grayordinate space using the following formulation:

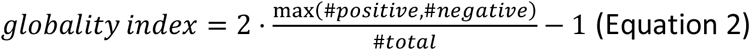

The above measure differs from an analogous measure used in (Glasser et al., 2018),as it has been modified to ensure that its bounded range lies between zero and one. This ensures compatibility with machine learning classifiers that are sensitive to feature ranges.

The last set of surface map features extracted includes the mean of the absolute values of the derived tICA surface map within each of the 360 parcels from the HCP-MMP1.0 parcellation (210 validation; (Glasser et al., 2016)) for both hemispheres, which is intended to explicitly aid in learning functional network-level information, thus aiding in the preservation of functional network activations that are seen in neural signal components. Features based on surface maps are computed on the 32k standard mesh.

Each spatial feature is computed for the raw group tICA spatial maps and the group-wise normalized tICA spatial maps, so that it can adapt to cross-dataset variability and also have a non-amplified version of a single-subject component. The absolute values of any spatial features that can be affected by the sign-flip of a spatial component, e.g., the mean value of a ROI-masked region, are added to the feature vectors to enable sign invariance.

##### Temporal features

Temporal features, including timeseries and power spectra, complement the spatial information provided by spatial maps and enable a more extensive characterization of the temporal dynamics of each component category. Outlier values in the tICA component of one or a few subjects usually suggest the presence of artifacts due to reconstruction, volume distortion, or dropout effect, whereas a pure neuronal signal should be smooth and devoid of outliers. Thus, the first handcrafted temporal feature is intended to convey the extent of this property, which is the square root of the timepoint-wise sum of the differences between the original tICA timeseries from the dominating subject and an updated timeseries with outlier values identified by the criteria in *Outlier Identification* and then filled by the nearest nonoutlier value (see Fig S13B).

Even though sICA is normally performed prior to tICA, thereby removing the great majority of spatial artifacts and nuisance signals, a small proportion of head motion artifacts may remain due to imperfect FIX predictions of sICA components or uncorrected distortions in the case of the 7T data in instances of large head movements that substantially change the magnetic field. An abrupt change in the amplitudes of successive timepoints suggests the presence of such an artifact, which can be quantified for a given timeseries using the complexity estimate (CE; (Batista et al., 2014)), defined as the L2 norm of the differences between consecutive data points in the time series.

The parameters of an autoregressive (AR) model at various orders of autoregression can be used to estimate temporal smoothness. Timeseries from signal components should be smooth and easy to fit using an AR model, helping to distinguish them from many types of artifacts. Here, we use five AR model parameters: the parameters from the 1st and 2nd order AR models, as well as the first two variances of the residual from the AR(1) and AR(2) models. The slope and intercept are two additional temporal features obtained by fitting a straight line to the variance of the residual in AR(p) models up to order p=8. The final set of temporal characteristics defined on timeseries includes the distributional properties of the dominating subject’s tICA timeseries (maximum, mean, standard deviation, kurtosis), along with the outlier effect indicator.

If there are no large neuronal signals or artifacts involved, a group tICA component is considered weak and thus has a low averaged frequency profile across the group. Therefore, the maximum value of the group-averaged tICA power spectrum is included as the first temporal feature defined on the power spectra. Additionally, three distributional properties (maximum, mean, standard deviation) are used to derive features for the dominating subject’s power spectra.

##### Greyplots

We create greyplots from timeseries data for each subject, using an improved method that uses cortical surface representations and parcellation with the HCP MMP 1.0, with display of rows scaled based on parcel size (Glasser et al., 2018, 2019; Power et al., 2017). Here we concentrated on two types of greyplots for the dominating subject of each group tICA component, 1) the <u>G</u>reyplot for the corresponding Individual tICA component (GIT), 2) the <u>G</u>reyplot for the sum of <u>A</u>ll individual <u>t</u>ICA components (GAT), which reflects the fluctuation of the individual tICA components on the full tICA modelled space.

##### Greyplot features

An averaged timeseries or the spatial map of a single subject is not always a good way to identify artifacts that have a spatiotemporal pattern, such as vascular delay or volumetric spatial distortion. Instead, subject-level features derived from the two types of greyplots were used to model subtle cortical area and time dimension characteristics. The first category of greyplot-related properties is the distribution property. A higher standard deviation of GIT indicates greater variability and potentially more pronounced spatial or temporal patterns, whereas a lower standard deviation suggests more uniform or stable values across the greyplot. Additionally, a higher mean value of GIT indicates a higher magnitude of the greyplot.

A single-subject global effect of an individual tICA component can be revealed when its GIT has an entire (or nearly complete) positive/negative stripe at a particular time interval that also appears at a nearly identical time range in the GAT. The rationale is that the individual tICA component of interest, from which the dominating subject’s cortical signal over time is modeled, exhibits a strong spatiotemporal pattern of fluctuations across many or all parcels and simultaneously dominates all other tICA components in this subject. Accordingly, we created a function to reflect this property. Given GIT and GAT, the timepoint-wise proportion of variance in GAT explained by GIT is measured by the eta^2^ (Cohen et al., 2008) which is then multiplied by two weight terms (both range from 0 to 1), using a rescaled sigmoid function on the average intensity of GIT and GAT at each timepoint (see Fig S12 for an example illustration). This stripe pattern is then quantified by locating the peaks of the resulting vector that has the same timepoint dimension as the original greyplot.

Greyplots can also reveal a pattern of cross-correlation between global artifact components associated with vascular delays, where the correlation between the GAT and lagged GIT is stronger for some lags than others. We then compute the cross-correlation between the original timeseries in each parcel and their lagged counterparts, yielding a cross-correlation matrix of size of parcel dimensions by number of lag values. The selected lag values are zero-centered. Since a high value is expected on only one side of the zero-lag axis (the time lag shouldn’t appear more than one time), this cross-correlation matrix should be asymmetric in the presence of a vascular delay pattern. Such asymmetries are evaluated by an eta-squared function on the original matrix and flipped matrix. We choose nonuniform ranges of lag values for the asymmetry comparison (from minus to plus 5, 50, 100, 200, and 400 frames), to reduce the computation compared to computing all possible integer frame lags, and also using multiple window sizes in case the delay is longer in a subset of subjects. We then use linear regression to explain the eta-squared values in terms of the window size. The resulting slope and intercept are included in the feature set to summarize the asymmetry pattern with varying ranges.

#### 2.3.4 Spatial latent features

The importance of obtaining spatial latent features is based on two primary considerations. First, spatial handcrafted features (see above section; (Salimi-Khorshidi et al., 2014)) largely rely on ROI masks that may not consistently identify intricate and nuanced spatial patterns within the data, whereas deep learning methods have demonstrated to be robust in this regard for volume and surface representations of fMRI data (Dahan, Fawaz, et al., 2022; Dahan, Williams, et al., 2022; Duc et al., 2020; Vu et al., 2020). Second, spatial handcrafted features are formulated at the group level, specifically for group maps and predominant subject maps. This does not harness the full potential of subject-level tICA spatial maps and thereby increases vulnerability to dataset variability and may reduce the ability to generalize. To address these issues, we have incorporated spatial latent features derived from both volume and surface autoencoders. These are self-supervised and are trained exclusively on subject-level tICA spatial maps. The aggregated group spatial latent features are subsequently derived by averaging the latent features of individual subjects within the group.

##### Volume map VQ-VAE

The Vector-Quantized Variational AutoEncoder (VQ-VAE) specializes in learning latent discrete representations of both 2D images and 3D volumes (Oord et al., 2017). In this model, the conventional VAE’s posterior is substituted with an embedding space *e* ∈ *R*^*K* × *D*^, where *K* represents the number of embedding vectors in the space and *D* signifies the dimension of each respective embedding vector. For each spatial code *z*_*e*_ (in our context, the volume latent feature), an element-wise quantization aligns it to its nearest vector *e*_*K*_ from a codebook using the L2 distance metric. This codebook is jointly learnt with all other model parameters through exponential moving average updates. The decoder then attempts to recreate the observations from this quantized latent space. Within our framework, the volume VQ-VAE loss function is characterized as follows:

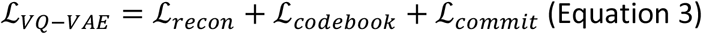

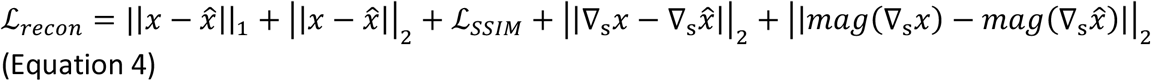

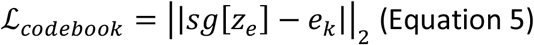

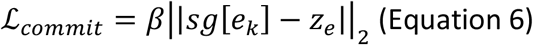

The Structural Similarity Index (SSIM) loss, denoted as *ℒ*_*SSIM*_, evaluates the disparity between the original input and its reconstructed version (Bergmann et al., 2018; Zhou et al., 2004). *∇*_*s*_ serves as the gradient operator using the Sobel filter, ensuring edge preservation. The *mag* function computes magnitude, while the *sg* operator denotes the stop-gradient operation, which passes zero gradients during the backpropagation process. The *ℒ*_*recon*_ loss addresses inconsistencies between the input and its reconstructed output by considering a multifaceted approach: evaluating L1 and L2 distances, assessing image quality metrics like luminance, contrast, and structure (*ℒ*_*SSIM*_), and ensuring preservation specific to edges. *ℒ*_*codebooK*_ and *ℒ*_*commit*_ mirror the variables described in the original VQ-VAE 2 paper (Razavi et al., 2019). To expedite the training process, we adopted the strategy of exponential moving average updates for the codebook (Oord et al., 2017) as a surrogate for the traditional codebook loss. We derived the volume latent features of size 256 by averaging the spatial code *z*_*e*_ across its feature channel dimension (see Fig 3 upper section).

**Figure 3.**
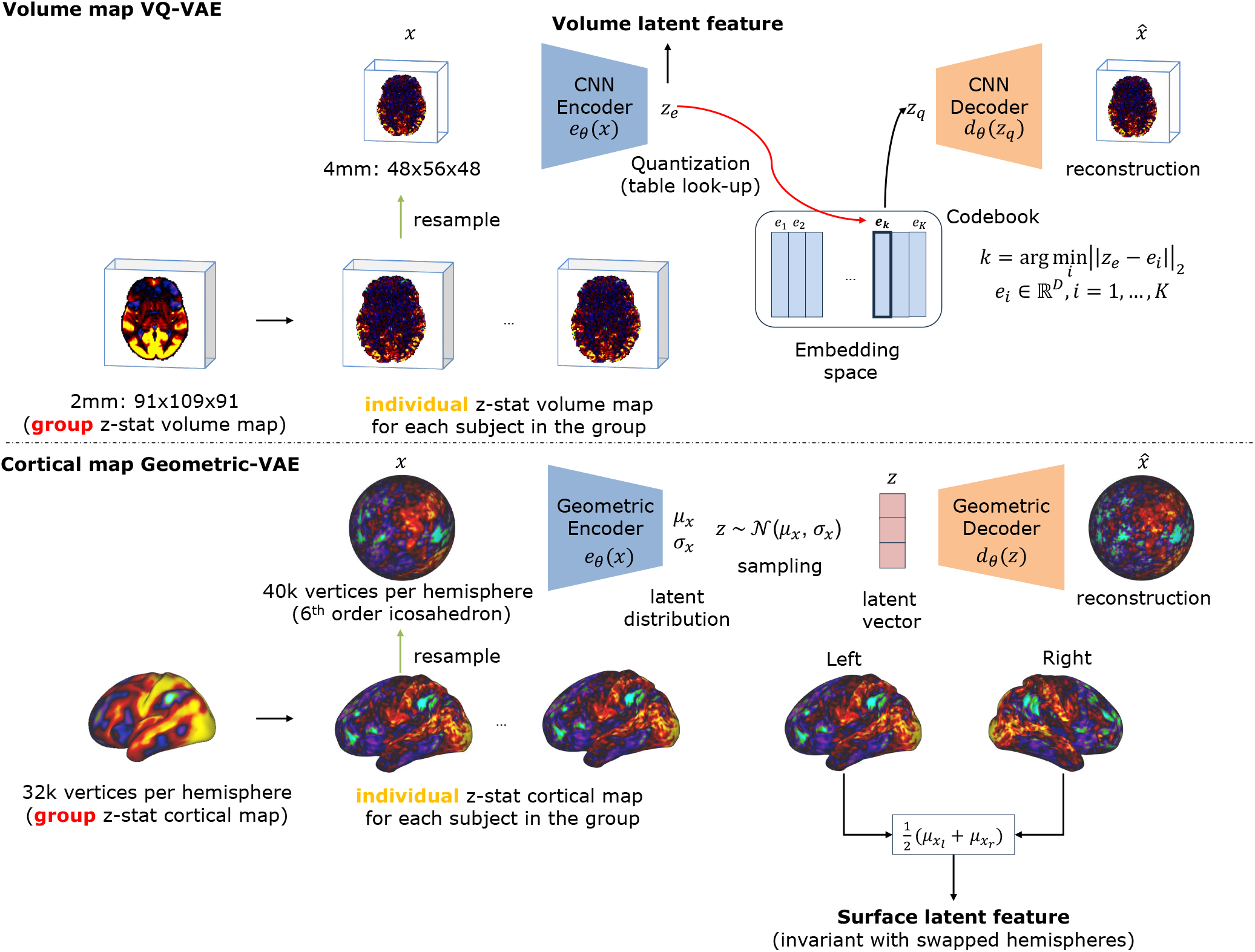
illustrates the workflows employed to extract individual(subject)-level volume and surface latent features. This is achieved using 1) the volume Vector-Quantized VAE (upper row) and 2) the cortical map geometric VAE (bottom row). Each encoder generates spatial latent features. Both the encoder and decoder are honed using a self-training schema, where the type of input data serves dual roles: as both the input and target. This training approach ensures that the acquired latent features remain unbiased by any labels. Importantly, to achieve hemispheric invariance, the surface latent features for individual tICA components are derived by averaging the feature vectors from both hemispheres.

##### Cortical map Geometric-VAE

The cortical geometric VAE was inspired by the cortical segmentation benchmark and the geometric deep learning model developed for UK Biobank parcellation (Fawaz et al., 2021; Williams et al., 2023). Diverging from their established geometric deep learning frameworks, which are typically U-Net based, our approach harnesses a VAE architecture. This architecture incorporates 5 encoding and 5 decoding blocks. Key components include the MoNet (mixture model networks; (Monti et al., 2017)) convolutional layer, parameterized using K=35 Gaussian kernels and polar pseudo-coordinates, subsequently followed by a SiLU (Sigmoid Linear Unit; (Ramachandran et al., 2017)) nonlinearity. The essence of a VAE is captured by its encoder, which models a posterior *p*(*z*|*x*) for a random variable *z*, considering the input *x*. Typically, a prior distribution *p*(*z*) is assumed to be *𝒩*(0,1). Meanwhile, the distribution *p*(*x*|*z*) regarding the input data is mapped via a decoder network. For our model, the cortical Geometric-VAE loss function is articulated as follows:

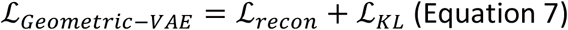

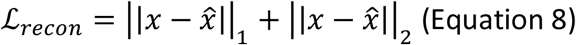

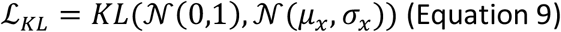

The *ℒ*_*recon*_ loss penalizes discrepancies between the input and its reconstructed output, as quantified by the L1 and L2 distances. Meanwhile, *ℒ*_*KL*_ aims to minimize the KL divergence between the approximated posterior and a standard normal distribution. The processing framework yields surface latent features of size 162 for each cortical map, encompassing both hemispheres. Given that a surface component and its mirror-symmetric counterpart (with left and right hemisphere swapped) share the same category, it is imperative to achieve hemispheric invariance. Without this invariance, a hemispheric bias in the training data might affect the final classification performance. Initially, for each hemisphere, the spatial code *μ*_*x*_ is averaged over the feature channel dimension. Subsequent to this, the mean of the latent features from both hemispheres is computed. This step ensures hemispheric invariance in the final representation, aligning with the need to account for the flexible nature of tICA surface maps (i.e. swapped hemispheres represent the same component category, see Fig 3 bottom section).

##### Training

The individual tICA z-stat spatial maps from the training datasets, as listed in Table 1, served as both inputs and targets for each autoencoder. We excluded the 7T datasets, given that the other large datasets already provided an ample number of training samples, each comprising hundreds of subjects. Specifically, the HCP-D training datasets with 100 and 160 subjects were used as validation sets, assisting in the prevention of overfitting and the fine-tuning of hyperparameters. As a result, our training data consisted of 172,107 individual maps, with an additional 4,680 maps allocated for validation.

For the volume VQ-VAE, with the workflow shown in the upper section from Fig 3, the 3T 2mm individual z-stat tICA volume map as input data was downsampled to 4mm using 3rd order spline interpolation. This adjustment facilitated compatibility with GPU memory limitations and expedited the training process. This model underwent training for five epochs to learn the volume latent features by minimizing the VQ-VAE loss, utilizing batch sizes of 80. The Adam optimizer (Kingma & Ba, 2014), paired with a learning rate of 1e-4, guided the training process. Of these five epochs, the model exhibiting the lowest reconstruction loss on the validation datasets was chosen as the best model to use.

For the cortical Geometric-VAE, with the workflow shown in the bottom section of Fig 3, the input individual z-stat tICA surface map (grayordinate) was initially divided between the two hemispheres. Following this, each 32k sphere underwent resampling to achieve a 40k sphere configuration, based on a 6th order icosahedron. (The icosahedron is a convex polyhedron made from triangles having 6 triangles at a vertex, except 12 vertices which have 5 triangles. The order represents the subdivision frequency *ν*, where each edge of the icosahedron is split into *ν* segments. The relation between the order and the number of vertices can be expressed by the following formula *NumVertices* = 12 + 10 × (4^*v*^ − 1). Hence, the 6^th^ order icosahedron has 40,962 vertices.) Given this bifurcation, the total count of training and validation samples for the cortical autoencoder was double that of the surface maps. Training was executed over four epochs, using a batch size of 1, and was facilitated by the AdamW optimizer (Loshchilov & Hutter, 2017) with a learning rate of 1e-3. Due to the substantial number of training samples, evaluations occur every 102,400 steps on the validation datasets. Ultimately, the model achieving the minimal reconstruction loss on the validation datasets was selected as the best model to use.

##### Feature generation

The spatial latent features were calculated at the individual level. To derive a group-level spatial latent feature, an averaging process was employed across all spatial latent features within a group, based on their specific types: either volume or cortical surface.

##### Implementation

The volume networks utilized the Medical Open Network for AI (MONAI) library (Cardoso et al., 2022) for their implementation, while the surface networks were developed using the PyTorch Geometric library (Fey & Lenssen, 2019). Each experiment was conducted on V100-SXM2 graphics cards, equipped with 32GB of memory and based on the Volta architecture by NVIDIA (Santa Clara, CA).

#### 2.4.4 Data augmentation

Machine learning typically benefits from incorporating as much data as possible into the training phase. Given the modest count of group tICA components (typically ranging from 68 to 123 per dataset as seen in Table 1), we initiated a pseudo-group data augmentation strategy to expand the training dataset’s sample size. This approach capitalizes on the idea that individual variability inherently serves as a form of data augmentation. By randomly selecting subsets of subjects to create a pseudo-group, we can generate an averaged spatial map and power spectrum for each selected group.

The spatial maps are particularly valuable for classification, and our findings showed that pseudo-group averaged spatial maps maintain a cosine similarity of over 0.85 with their original group counterparts, provided that more than 40 subjects are randomly selected from a pool of 535 subjects from the HCP-YA resting state dataset (as illustrated in Fig S4). This ensures a consistent spatial profile at the group level. We further augmented the group power spectrum, given that only one handcrafted feature is designated for this data type. With a large pool of subjects available for a group, this augmentation approach can significantly amplify the training dataset’s size. Additionally, augmenting with smaller group sizes replicates scenarios where group tICA datasets have limited subjects.

Since this augmentation method diversifies data at the group level for both spatial and power spectrum facets, each newly created feature vector boasts distinct group spatial and power spectrum attributes, while the temporal, individual, and outlier features are copied from the original full group, rather than being recomputed. Pseudo-group spatial map labels are defined to be consistent with the raw full group tICA component, though in practice this labeling may not always be precise, because of the existence of single subject-dominated components or a limited group size that can result in a dissimilar pseudo-group spatial map compared with the raw group ones (the low cosine similarity shown in Fig S4). However, noise introduced from the labeling of the pseudo group serves as a regularization, compelling the classifier to prioritize non-group-level-spatial features, such as the dominant subject features or temporal patterns.

Theoretically, given a group of *N* subjects, the upper limit of augmented datapoints is as 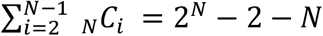 where *i* represents the pseudo-group size, ranging from 2 to *N* − 1. (*i* = 1 means to treat each individual subject’s tICA component as an additional pseudo group; *i* = *N* stands for the raw full group). During training, we initially randomly sampled a group size between 1 and *N* followed by uniform subject sampling from the entire training group. Subsequently, we averaged the spatial maps and power spectra to establish the augmented dataset. Instead of using this augmentation method on the fly, we generated 100 augmented datapoints locally for every group tICA component in the training datasets, expanding the training samples from 750 (the raw group tICA components) to 75,750 datapoints (the raw group tICA components plus the pseudo-group tICA components). The pseudo group sizes are uniformly sampled between 1 and each tICA dataset’s original size.

#### 2.4.5 Classification metrics

To evaluate the discriminative power of our model candidates in distinguishing signal components from artifactual ones, we compared predicted labels to manual labels using a range of criteria given in Table 2 along with their definitions and specialties. It is worth noting that tICA component classification is a label-imbalanced binary classification task, where signal components consistently exceed artifactual ones, as evidenced by Table 1.

**Table 2.** An overview of measures used to evaluate the performance of proposed models on a broad scale, including balanced accuracy and F1 score that rely on a decision threshold of 0.5, as well as multiple AUC-based measures for robust and comprehensive evaluation without considering the decision thresholds. (TP: true positive; TN: true negative; FN: false negative; FP: false positive; TPR: true positive rate; TNR: true negative rate)

| metric | definition | specialty |
| --- | --- | --- |
| Balanced accuracy | $\frac{1}{2}(\frac{TP}{TP + FN} + \frac{TN}{FP + TN})$ | Average between sensitivity (TPR) and specificity (TNR), which takes the label imbalance into account, and uses decision threshold 0.5. |
| ROC-AUC | Area under ROC curve | A robust and threshold-free metric describing the model's prediction ranking performance. The value can be biased by the imbalance of data. |
| PR-AUC | Area under the precision-recall curve | A robust and threshold-free metric describing the model's overall performance for predicting the positive instances. The value is not biased by the imbalance of data. |
| NPV-Specificity AUC | An analogous metric to PR-AUC, where it measures the area under the negative predictive value-specificity curve | A robust and threshold-free metric describing the model's overall performance for predicting the negative instances. The value is not biased by the imbalance of data. |
| Correct prediction variance percentage | The total variance correctly classified by the classifier | This measure is per-dataset based, ranging from 0% to 100%, calculated by summing up the variances of the components that belong to the True Positive and True Negative predictions. |
| FN prediction variance percentage | The total variance incorrectly predicted as negative which are positive (signals) | This measure is per-dataset based, ranging from 0% to 100%, calculated by summing up the variances of the components that belong to the False Negative predictions. |
| FP prediction variance percentage | The total variance incorrectly predicted as positives which are negative (artifacts) | This measure is per-dataset based, ranging from 0% to 100%, calculated by summing up the variances of the components that belong to the False Positive predictions. |

Our evaluation encompasses both threshold-free and threshold-based metrics. Among the threshold-free metrics, ROC-AUC (Receiver Operating Characteristic - Area Under the Curve), PR-AUC (Precision Recall - Area Under the Curve), and NPV-Specificity AUC (Negative Predictive Value Specificity - Area Under the Curve) assess each model’s comprehensive performance across both classes. However, ROC-AUC can be overly optimistic when there is a significant class imbalance. PR-AUC focuses on the positive class while omitting true negatives. NPV-Specificity AUC concentrates on the negative class while omitting true positives. On the other hand, balanced accuracy, though highly beneficial for imbalanced datasets, operates on a predetermined threshold — in this context, 0.5 (0 for negative class: artifacts; 1 for positive class: signals). The other threshold-based metrics are based on the variance percentages for True Positive (TP), True Negative (TN), False Positive (FP), and False Negative (FN) predictions. The correct prediction variance percentage covers the TP and TN predictions together, describing the amount of variance correctly identified by the model which focuses on the ability of the model to handle the most critical components. The FP prediction variance percentage only focuses on the FP predictions, describing the variance of artifactual components that are wrongly predicted as signals, while the FN prediction variance percentage focuses on the FN predictions, describing the variance of signal components that are wrongly predicted as artifacts.

The rationale for choosing any particular metric is strongly related to the application scenario: For instance, if the goal leans towards minimizing missed signal components, PR-AUC emerges as the optimal metric. Conversely, if the objective is to identify the maximum number of artifactual components, NPV-Specificity AUC should be prioritized. A pragmatic approach often involves first discerning the use case, followed by fine-tuning the decision threshold based on the curves used from the threshold-free metrics assessed on a validation dataset. Throughout our approach, we have accorded greater emphasis to threshold-free metrics to ensure a comprehensive evaluation of the classifier’s performance.

#### 2.4.6 The hierarchical classifier

Inspired by FIX (Salimi-Khorshidi et al., 2014), we formulated a hierarchical classifier based on ensemble learning. This classifier leverages both handcrafted features from multiple sources as domain-knowledge-related information and group-averaged spatial latent features as visual and perceptual knowledge related information, concatenating them into a single feature vector. Contrary to FIX’s approach, we opted not to feed features into traditional classifiers based on their feature types. This decision primarily rests on the rationale that the outlier indicator feature — employed to discern if a component’s property is determined by its predominant subject — should be evaluated in concert with other features. Our approach allows the model to learn the interaction between group or individual-level features and the outlier characteristic. Notably, our hierarchical architecture sidesteps the need to meticulously fine-tune a single optimal classifier, as the ensemble of base learners combine to make a robust classifier.

Our selection process started with an initial pool of ten potential base learners. This pool included five tree models: decision tree, random forest (Breiman, 2001), Extra tree (Geurts et al., 2006), XGBoost (Chen & Guestrin, 2016), and bagging tree (Ho, 1998); two k-NN (nearest neighbor) models: uniformly weighted k-NN and distance-weighted k-NN (Fix & Hodges, 1989); two multi-layer perceptrons (MLPs) (Glorot & Bengio, 2010; He et al., 2015; Hinton, 1989; Kingma & Ba, 2014): single-layer and dual-layer; and logistic regression (Zhu et al., 1997). Every base learner was trained using five distinct random seeds on the augmented dataset, each original training sample being expanded with 100 augmented data points, culminating in a total of 75,750 training data points. Performance evaluations were then conducted on the validation set (HCP-D datasets with 100 and 160 subjects). The top five base learners were selected based on their excellent performance across ROC-AUC, PR-AUC, NPV-Spec AUC, and balanced accuracy (See Fig S8 for details). These comprise the extra tree, a single-layer MLP, a dual-layer MLP, XGBoost, and logistic regression (Fig 4B). Each selected base learner has unique strengths and focal patterns that fortify the ensemble’s collective learning powers, making our final hierarchical classifier both robust and generalizable. For instance, tree-based models adeptly capture discrete, non-linear, and interaction-driven relationships, whereas MLPs excel in discerning complex, continuous, and deeply non-linear patterns.

**Figure 4.**
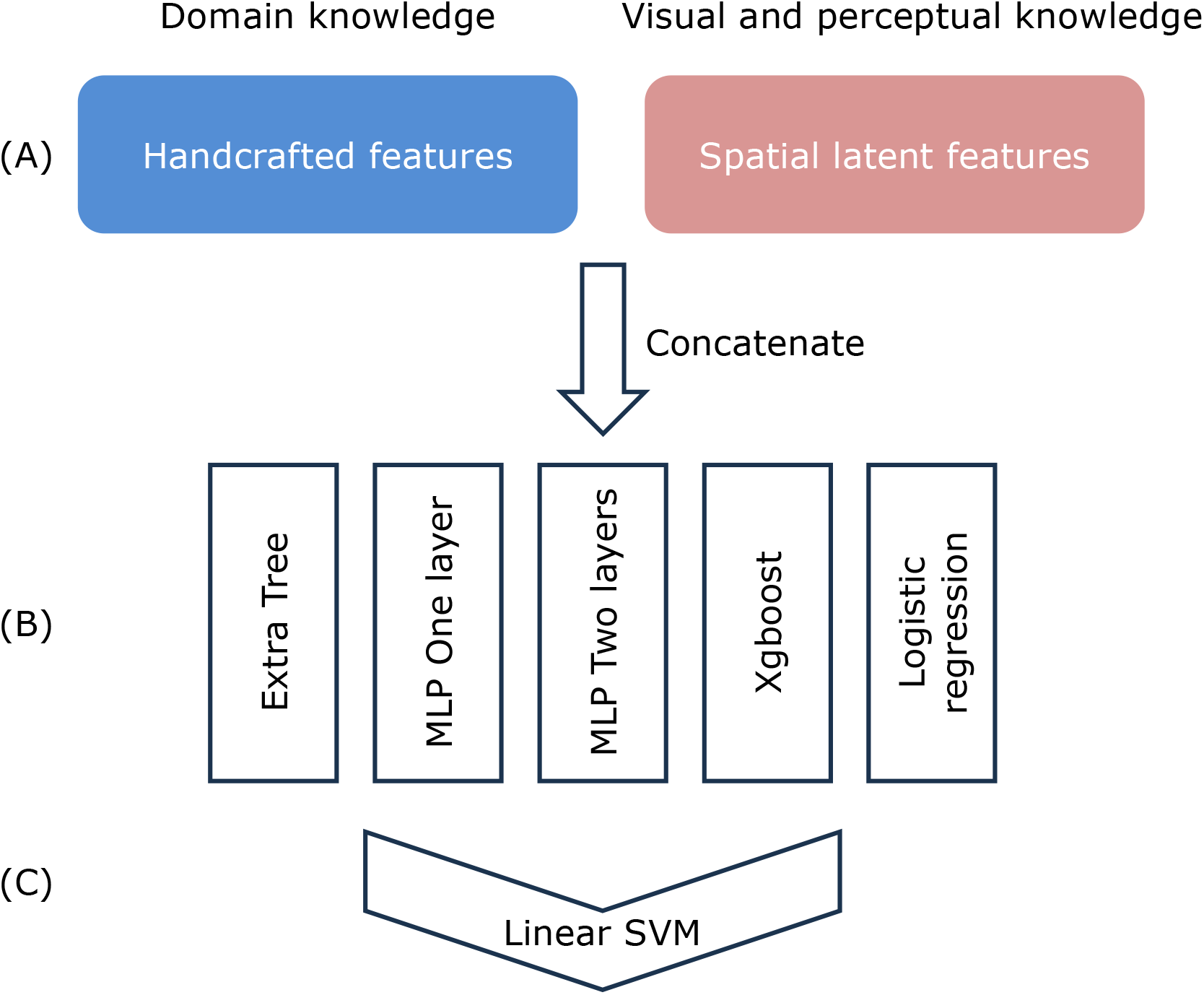
The hierarchical classifier for tICA classification. The handcrafted features and spatial latent features (level A) are fed into five base learners (level B)., The stacking layer (level C) is a linear SVM that fuses the probability arrays generated by the base learners and then outputs the final probability/class.

To combine the decisions and provide the final prediction, a high-level stacking layer, a Support Vector Machine (SVM; (Cortes & Vapnik, 1995)) with a linear kernel, was trained on the probabilities predicted by the five base learners (Fig 4C).

##### Training and Evaluation

Following the learner selection process, the hierarchical classifier was trained using the entire training dataset, encompassing the validation datasets, and subsequently tested on the evaluation set. Importantly, both the base learners and the stacking layer are considered with class imbalance, where more weight is assigned to the minority class based on its frequencies in the training data. We undertook evaluations of the datasets in two distinct manners: (i) retaining all datapoints, and (ii) excluding datapoints that exhibit volume distortion (in 7T dataset exclusively) or are ambiguously labeled as weak components with the subcategory (weak signal, weak subcortical, or weak other). The latter evaluation approach ensures that our proposed model effectively addresses the most critical categories.

##### Implementation

The XGBoost learner was implemented using the XGBoost module with scikit-learn wrapper. All the other base learners were implemented using the scikit-learn library (Pedregosa et al., 2011). Every model was trained on CPU only, and no GPU acceleration was involved during training.

## 3. Results

The Results are grouped into two major sections, each with multiple subsections. Section 3.1 and Section 3.2 deal with early stages of tICA pipeline automation. This begins with automatic dimensionality selection (#3.1). Here, we unveil the method’s capability for achieving robust dimensionality estimations and high reproducibility. Subsequently, we investigate single-subject effects in HCP-Style large-scale datasets (#3.2), showing the relationship between total subject count/timepoints and the fraction of potential single-subject dominated components among the group components. Section 3.2 also provides recommendations for selecting the appropriate tICA mode given a study’s total number of timepoints, which should prove valuable to users.

The second segment of our automation strategy encompasses the tICA classifier and includes five key aspects: 1) visualization of latent features, demonstrating how these features can effectively represent similar group components (#3.3), 2) a comparative analysis across different feature settings, highlighting our rationale for combining handcrafted features and spatial latent features as the final configuration (#3.4), 3) an assessment of the effectiveness of pseudo-group data augmentation (#3.5), 4) benchmarking of the tICA classifier, evaluating its performance on the most critical components and all eleven evaluation datasets (#3.6), and lastly 5) an exploration of the classifier’s interpretability (#3.7).

### 3.1 Effect of multiple Wishart distributions on group sICA

Overall, our goal with dimensionality estimation is to find the greatest number of highly reproducible ICA components. We tested multiple numbers of Wishart distributions (WD1 through WD8) on the PCA dense timeseries produced after MIGP using the full HCP Young Adult released dataset (1071 subjects) before estimating the sICA dimensionality. The cluster quality index Iq from ICASSO (Himberg & Hyvarinen, 2003) was used to measure the reproducibility of each sICA component, with values ranging from 0 to 1, with numbers closer to 1 representing more similar components from ICA runs with different random initializations. Given the relatively high sICA dimensionality estimated for WD1 – WD3, which also have low Iq, only WD4 – WD8 were used to derive the Iq values to reduce the computational burden. The averaged Iq value for a given dimensionality was used to describe the overall reproducibility under each WD. Three random experiments were launched to consider the randomness of the estimations. This schema was applied to HCP-YA 3T resting state, 3T task fMRI, 7T resting state, 7T movie and 7T retinotopy. Fig 5 shows results for HCP-YA 3T resting state (Fig 5A, B) and 3T task fMRI (Fig 5C, D); results for the other datasets are in Fig S1. As the number of Wishart Distributions (WD) increases, sICA dimensionality monotonically declines because an increasing portion of unstructured random noise is removed from the MIGP data, and the quality index Iq tends to increase, but not monotonically. However, as the number of Wishart Distributions increases further, the structured component of the signal is also reduced, resulting in even lower sICA dimensionality (Fig 5A, C) and variable effects on Iq (Fig 5B, D). A single WD is insufficient, as it produces nearly 1000 components, resulting in a massive oversplitting (whereby known network patterns are split across multiple components, and the most structured (i.e., spatially autocorrelated) of the unstructured noise is modeled into ICA components), extremely high computational burden, and difficulty with ICA convergence. A tradeoff point is evident in the Iq plots (Fig 5B, D) where the Iq declines from WD6 to WD7. Even though WD8 has a higher Iq value (0.93 for 3T resting state, 0.97 for 3T task fMRI), the low sICA dimensionality (56 for 3T resting state, 39 for 3T task fMRI) signifies that the combination conveys less information. Thus, determining the optimal WD number involves two considerations: 1) a reasonable sICA dimensionality (d=84 for 3T resting state and d=70 for 3T task fMRI from (Glasser et al., 2018)), 2) a high Iq value. We chose WD6 (labeled bold red) for HCP-YA 3T resting state, 3T task fMRI, 7T resting state, 7T movie, HCP-Aging and HCP-Development and WD5 for HCP-YA 7T retinotopy to include more sICA components while ensuring the sICA dimensionality was in a range close to the other 7T decompositions. The medians of each estimated sICA dimensionality and each averaged Iq value among the three random experiments are indicated in the legends.

**Figure 5.**
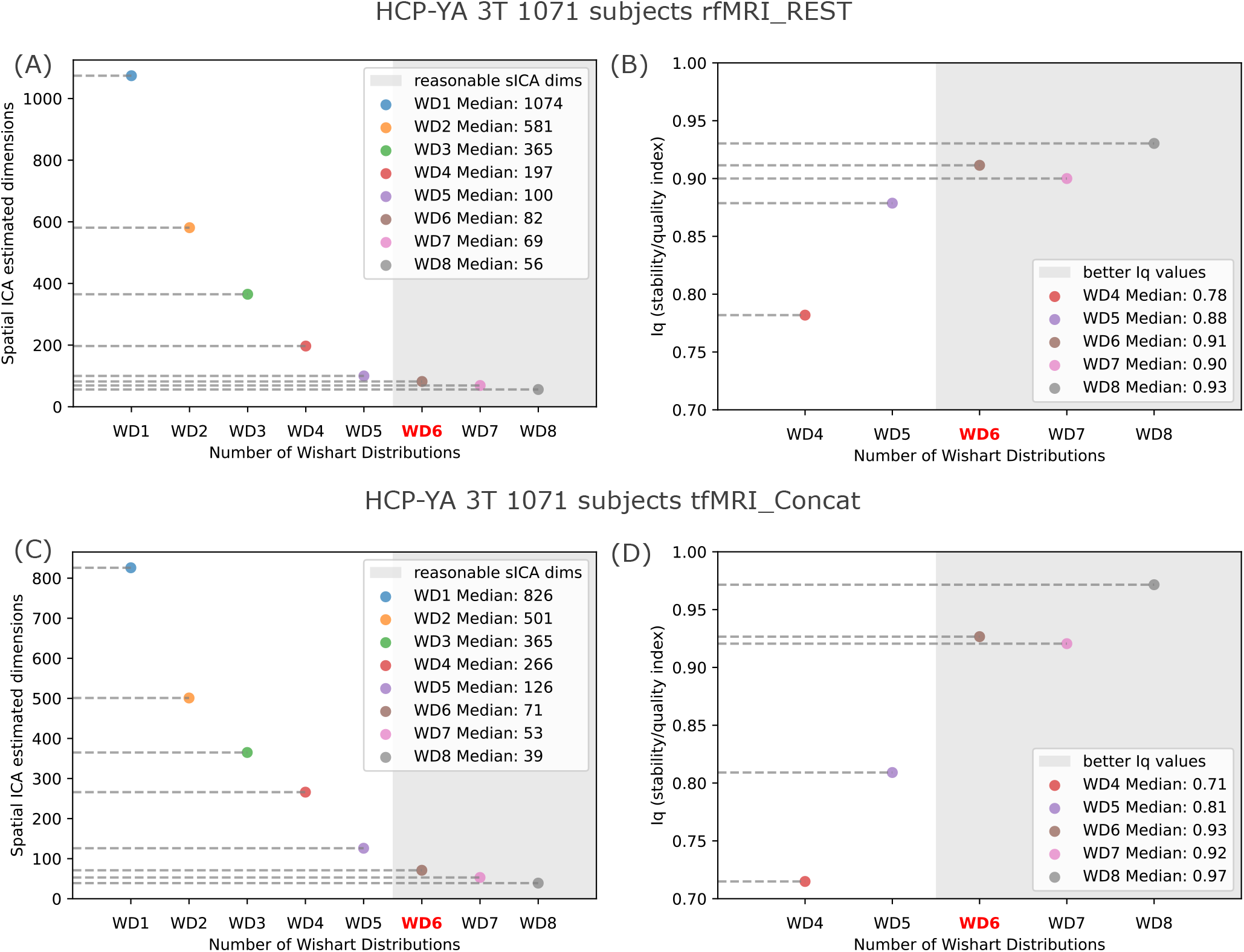
The estimated sICA dimensionality and the averaged Iq value across sICA components for HCP-YA 3T resting state and 3T task fMRI datasets. A single WD produces close to one thousand sICA components while multiple WD removes the unstructured noise from MIGP and reduces the sICA dimensionality to a feasible range for classification. The reproducibility of sICA components is measured by Iq. WD6 is chosen based on the number of sICA dimensionality containing enough information and high reproducibility. A: WD6 results in 82 sICA components for the MIGP of HCP-YA 3T resting state; B: WD6, WD7 and WD8 have high reproducibility measured by Iq, while WD6 is chosen because of sufficient sICA dimensionality; C: WD6 results in 71 sICA components for the MIGP of HCP-YA 3T task fMRI; D. figure has similar trend to B, and thus WD6 is chosen.

### 3.2 Effect of the number of subjects and total timepoints on the temporal ICA single-subject effect

The temporal ICA single-subject effect involves the splitting of strong variance from individual subjects into separate components, rather than creating components that reflect variance consistent within a group. This behavior is undesirable because it may fail to split signal and artifact within individual subjects and tends to reduce the number of reproducible group-level components. This effect is progressively dampened as subject numbers/total timepoints increase, as each individual contributes less to the overall variance. The HCP Young Adult 3T datasets and the HCP Lifespan (Aging and Development) datasets are used to illustrate this trend. The group tICA decompositions for each of the released datasets are shown in Table 3. 10 to 800 subjects at intervals of 5 were randomly sampled from the full list of subjects for each dataset to form the group concatenated spatial ICA dense timeseries as the input for group tICA (Step 5). The number of tICA components was set to the same as the released version which respects the group averaged results and avoids extensive computation of group sICA on the subset group. After the group tICA, individual tICA components (timeseries) were de-concatenated and the variance for each individual tICA component was computed. To illustrate the single subject effect without manual labeling of the random groups, if certain individual tICA components show high variance as outliers, then the group component is defined to be dominated by individual subjects. To measure such variability across varying subjects, for each group component, we computed the coefficient of variation of the individual variances of a pseudo group. An outlier coefficient of variation within the group indicates a single subject dominated group component. To better illustrate the trend under different outlier detection thresholds, we used 3, 6, and 9 scaled Mean Absolute Deviations (MAD). We performed random sampling 5 times for a given subject number. We fit a linear regression to the mean of the 5 random experiments and show the goodness of fit R^2^, the p value and the fitting parameters. In Fig 6, the fitted curve shows a decreasing trend of the number of potentially single subject dominated group components when more subjects are involved in the group decomposition, indicating the single subject effect is dampened. To recommend a cut-off of the single-subject effect, a 7% threshold was chosen from the HCP-D dataset with the most severe single-subject effect under the lowest criteria (3 MAD) among the three choices (see Fig 6 bottom left) to provide a strict, but reasonable threshold in the range of 5% to 10%. The intersection between the cut-off line and the fitted curve roughly lies under the second half of the fitted curve. When we used more extreme criteria to decide outliers (9 scaled MADs), the trend holds. In the HCP Young Adult 7T dataset in Fig S3, the single subject effect shows an increasing trend, which might reflect the limited number of 7T subjects.

**Table 3.** The tICA decomposition info for the official released HCP-YA, HCP-A and HCP-D datasets.

| HCP datasets | # Subjects | # Timepoints per subject | Resolution (mm) | # ICs | fMRI runs |
| --- | --- | --- | --- | --- | --- |
| Young Adult | 1071 | 4800 | 2 | 82 | resting state |
| Young Adult | 1071 | 3880 | 2 | 72 | concatenated task fMRI with 7 tasks |
| HCP-Aging | 1798 | 2721 | 2 | 86 | concatenated fMRI, resting state & 3 tasks |
| HCP-Development | 1695 | 3200 | 2 | 122 | concatenated fMRI, resting state & 3 tasks |

**Table 4.** The performance of the hierarchical classifier with combined feature settings on each evaluation dataset. The mean metric value from five random experiments is reported along with the standard deviation. By default, each of the evaluation datasets has been sICA recleaned (See supplementary methods for detail). YA resting state: HCP-Young Adult 3T 535 subjects with resting state fMRI data; YA task: HCP-Young Adult 3T 535 subjects with concatenated task fMRI data; YA 7T resting state: HCP-Young Adult 7T 87 subjects with resting state fMRI data; YA 7T resting state: HCP-Young Adult 7T 87 subjects with resting state fMRI data; YA 7T movie: HCP-Young Adult 7T 87 subjects with movie task fMRI data; YA 7T retinotopy: HCP-Young Adult 7T 87 subjects with retinotopy task fMRI data; Aging concat w/o reclean: HCP-Aging 3T 617 subjects with non-sICA-recleaned concatenated fMRI data; Development concat w/o reclean: HCP-Development 3T 314 subjects with non-sICA-recleaned concatenated fMRI data; Development 60 subjs: HCP-Development 3T 60 subjects with concatenated fMRI data; Development 100 subjs: HCP-Development 3T 100 subjects with concatenated fMRI data; Development 60 subjs (initialize): the same as Development 60 subjs but the decomposition is initialized by the mixing matrix from YA resting state; Development 100 subjs (initialize): the same as Development 100 subjs but the decomposition is initialized by the mixing matrix from YA resting state.

| Evaluation datasets | ROC AUC | PR AUC | NPV Spec AUC | Balanced Accuracy | Correct prediction variance percentage % | FN prediction variance percentage % | FP prediction variance percentage % |
| --- | --- | --- | --- | --- | --- | --- | --- |
| YA resting state | 0.96 ± 0.01 | 1.00 ± 0.00 | 0.87 ± 0.02 | 0.83 ± 0.04 | 97.9 ± 0.4 | 0.0 ± 0.0 | 2.1 ± 0.4 |
| YA task | 1.00 ± 0.00 | 1.00 ± 0.00 | 1.00 ± 0.00 | 0.93 ± 0.09 | 99.5 ± 0.6 | 0.0 ± 0.0 | 0.5 ± 0.6 |
| YA 7T resting state | 0.99 ± 0.00 | 1.00 ± 0.00 | 0.97 ± 0.01 | 0.90 ± 0.02 | 96.4 ± 0.5 | 0.7 ± 0.3 | 2.9 ± 0.3 |
| YA 7T movie | 0.99 ± 0.00 | 1.00 ± 0.00 | 0.93 ± 0.01 | 0.81 ± 0.03 | 96.9 ± 0.6 | 0.5 ± 0.0 | 2.5 ± 0.6 |
| YA 7T retinotopy | 0.99 ± 0.00 | 1.00 ± 0.00 | 0.98 ± 0.01 | 0.92 ± 0.02 | 98.6 ± 0.5 | 0.0 ± 0.0 | 1.4 ± 0.5 |
| Aging concat w/o reclean | 0.98 ± 0.00 | 1.00 ± 0.00 | 0.94 ± 0.01 | 0.73 ± 0.00 | 94.2 ± 0.0 | 0.0 ± 0.0 | 5.9 ± 0.0 |
| Development concat w/o reclean | 0.96 ± 0.00 | 0.99 ± 0.00 | 0.90 ± 0.01 | 0.86 ± 0.01 | 93.0 ± 0.3 | 4.1 ± 0.4 | 3.0 ± 0.4 |
| Development 60 subjs | 1.00 ± 0.00 | 1.00 ± 0.00 | 0.99 ± 0.00 | 0.97 ± 0.01 | 96.8 ± 1.0 | 2.9 ± 1.1 | 0.3 ± 0.5 |
| Development 100 subjs | 0.95 ± 0.00 | 0.98 ± 0.00 | 0.89 ± 0.01 | 0.87 ± 0.02 | 88.6 ± 0.8 | 8.7 ± 0.4 | 2.8 ± 0.6 |
| Development 60 subjs (initialize) | 0.90 ± 0.01 | 0.98 ± 0.00 | 0.69 ± 0.01 | 0.73 ± 0.00 | 92.2 ± 0.6 | 4.2 ± 0.6 | 3.6 ± 0.0 |
| Development 100 subjs (initialize) | 0.90 ± 0.01 | 0.98 ± 0.00 | 0.67 ± 0.06 | 0.68 ± 0.06 | 93.5 ± 1.0 | 1.4 ± 0.6 | 5.1 ± 0.9 |

**Figure 6.**
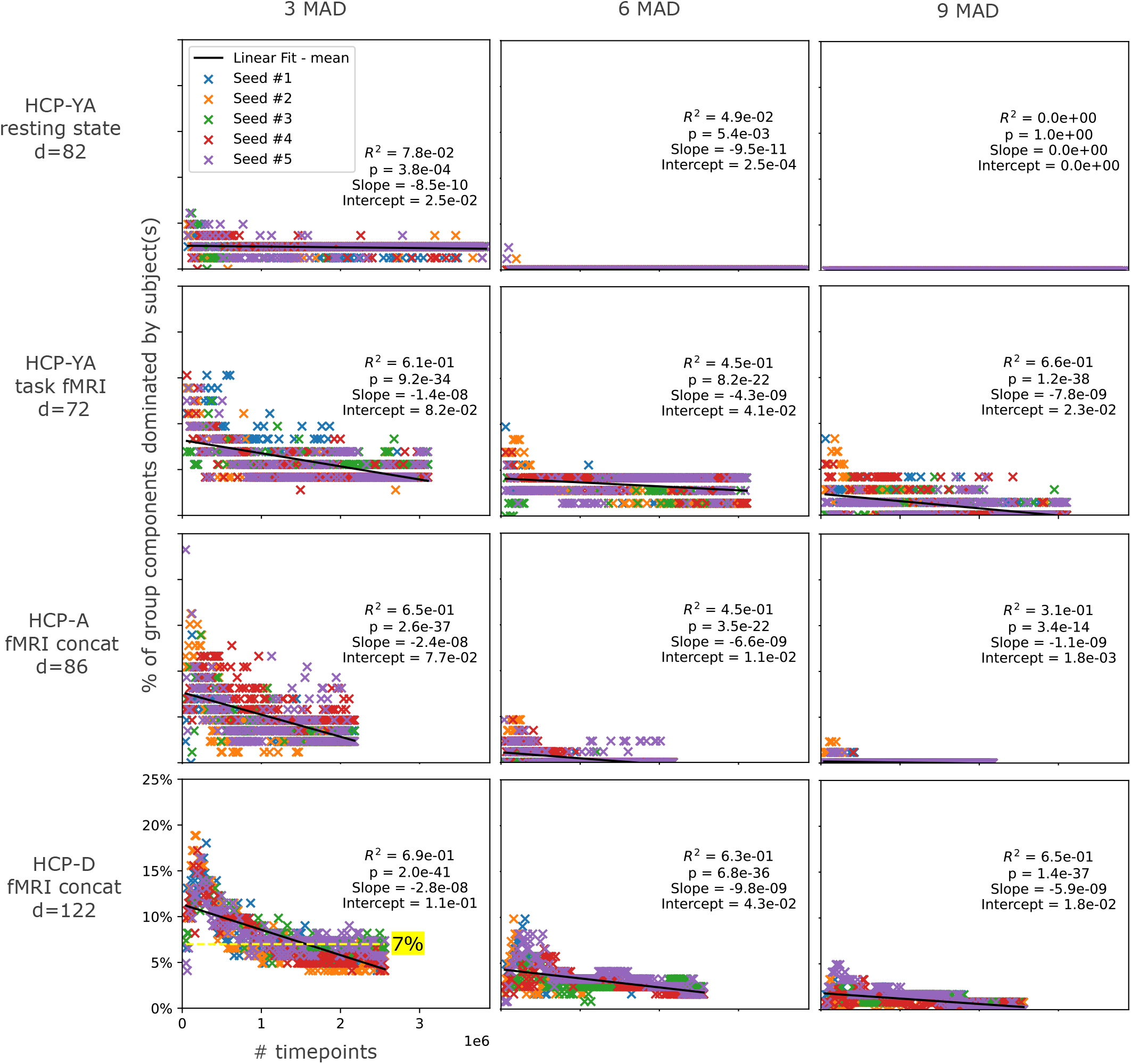
shows the single-subject effect (percentage of group components dominated by individual subjects) plotted as a function of the number of timepoints included in the group tICA for different datasets (rows) and for different outlier detection thresholds (columns). Row 1: HCP-Young Adult (YA) resting-state; Row 2: HCP-Young Adult (YA) task-fMRI; Row 3: HCP-Aging (A) all fMRI; Row 4: HCP-Development (D): all fMRI. Outlier detection threshold = 3 scaled Mean Absolute Deviations (MADs) (left column), 6 scaled MADs (center column), and 9 scaled MADs (right column). Due to computation limitations, a maximum of 800 subjects were evaluated for each plot. The maximum total timepoints come from HCP-YA rfMRI_REST, which has 4800 × 800 = 3.84e6. Five random experiments were run to consider the randomness in ICA decomposition. A linear fit was performed for the mean value in each subplot to illustrate the dampened single subject effect when more subjects/timepoints are fed into the group tICA. The dampening effect is consistent under outlier detection thresholds from loose to strict.

Based on the above analysis, we have endeavored to provide guidelines for the recommended number of total timepoints under the three tICA modes we have developed. This guideline aims to facilitate the adjustment of single subject effects within both the HCP officially released datasets and newly acquired HCP-style datasets from the community. We initially used the 3 scaled MAD metrics, denoted as the first column in Fig 6, as a foundational reference point, incorporating a broader consideration of outliers. Using the mean value derived from five random experiments for each subject number, we fitted a Histogram-based Gradient Boosting Tree regressor (HGBT; Ke et al., 2017), constraining it to exhibit a monotonic decrease in accordance with our observation that increased subject count leads to a reduction in single-subject effects. Subsequently, the resulting stepwise fitting curve in Fig 7 is divided into two distinct regions by a red dashed line using a thresholding methodology. The recommended number of timepoints is identified as the first plateau which consistently demonstrates less than 7% dominance by single-subject components across large-scale HCP datasets. Notably, this 7% threshold (yellow dashed line in Fig 6) corresponds to a range of 5 to 9 components within the group tICA decomposition, with a total component count spanning from 82 to 122. In the case of HCP-YA resting state data (Fig 7A), the influence of single-subject effects is relatively modest, thus permitting the reasonable utilization of all tICA modes. As for HCP-YA task-based fMRI data (Fig 7B), we have opted for a threshold of 620k timepoints, corresponding to ∼170 subjects, below which USE and INITIALIZE are the only recommended tICA modes. In the case of HCP-A with concatenated fMRI data (Fig 7C), a threshold of 480k timepoints was selected, corresponding to ∼176 subjects. Lastly, for HCP-D with concatenated fMRI data (Fig 7D), a more substantial threshold of 1 million timepoints has been chosen, encompassing ∼312 subjects. As a result, we recommend that users follow these guidelines and use the recommended number of timepoints to identify the most suitable tICA mode for their specific datasets. For users having sufficient computational resources and time, a more comprehensive strategy is encouraged, involving the application of all three modes to facilitate outcome comparisons.

**Figure 7.**
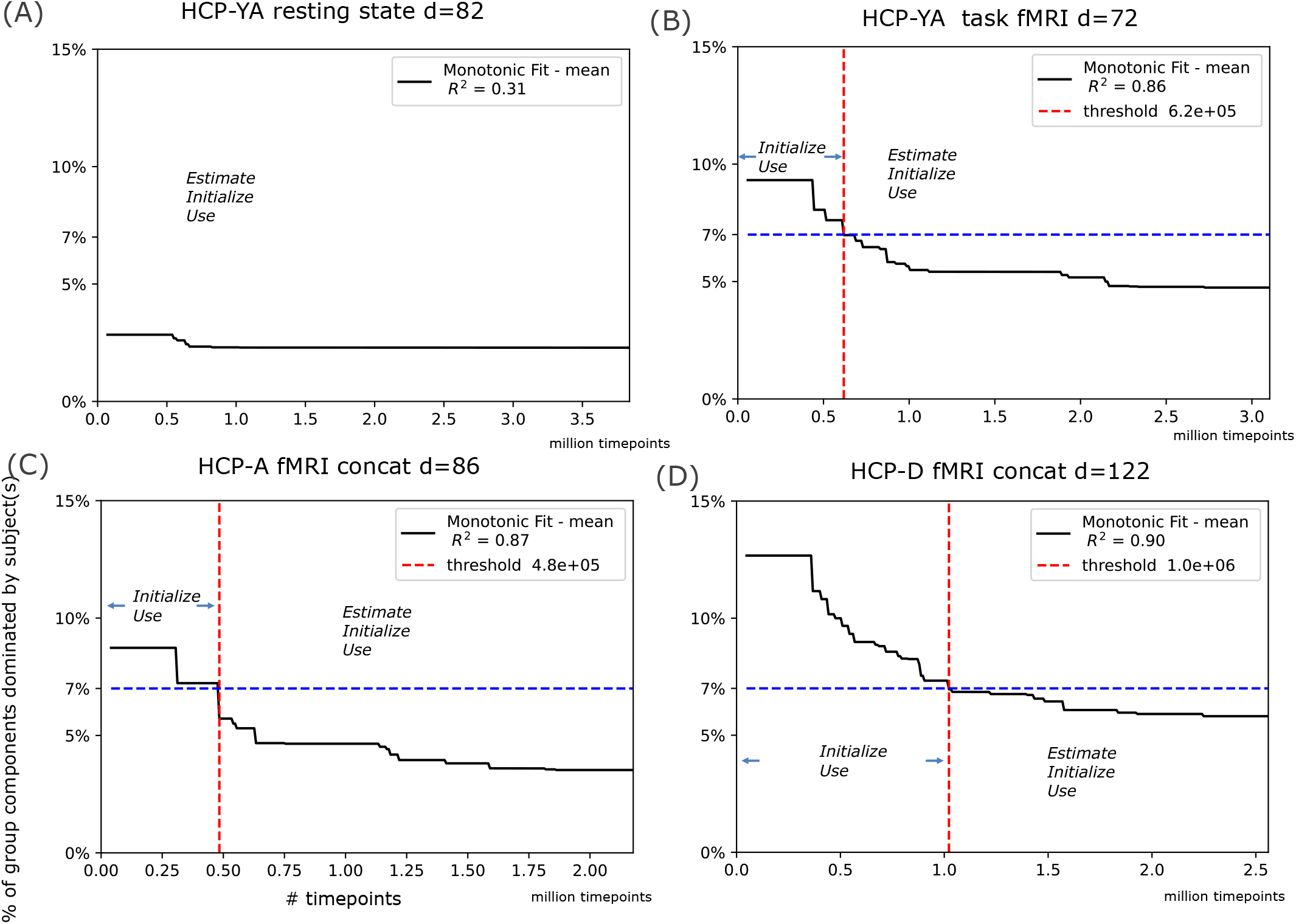
Recommended choice of tICA mode for different numbers of timepoints. 3 MAD is used to detect outlier variances and to estimate the group components dominated by individuals. A histogram-based gradient boosting tree regressor is fit to the mean of the percentages across five runs shown as different seeds in Fig 6 with the x axis as the number of timepoints, constrained to be monotonically decreasing. The recommended threshold for timepoints is determined by identifying the first plateau that has lower than 7% single subject dominated components (roughly 6, 5, 6 and 9 components for HCP-YA resting state, HCP-YA task fMRI, HCP-A and HCP-D with concatenated fMRI, respectively). A vertical red dashed line is drawn as the boundary between the region where the precomputed modes are preferred and where all the modes can be applied. Note that HCP-YA resting state doesn’t clearly suffer from the single subject effect, thus all the three modes can be applied. The recommended number of timepoints are shown as the threshold in the legend, which is the equivalent of around 170, 176, 312 subjects for HCP-YA task fMRI, HCP-A and HCP-D with concatenated fMRI, respectively.

We provide two examples here to assist users in determining which mode to use: 1) 270 subjects with 440k timepoints, and 2) 40 subjects with 80k timepoints.

First, it is important to take into account the age distribution of the overall subject group to identify the HCP dataset that best matches the customized dataset. Assuming the closest match is the Young Adult subjects, here are the recommended modes for each case:

1. If case #1 only involves resting state fMRI runs, users are encouraged to apply the ESTIMATE or INITIALIZE modes to better fit the data while USE mode can be selected as a tradeoff to reduce the computational burden (refer to Fig 7A). The ESTIMATE and INITIALIZE modes will both estimate the tICA mixing matrix that can separate the given dataset into independent temporal components, but INITIALIZE will use a pre-existing sICA decomposition and start tICA decomposition with a previous result rather than random initialization.
2. If case #1 includes task fMRI runs, it is advisable to use the USE or INITIALIZE modes instead, as indicated in Fig 7B (440k < 620k).
3. Given that case #2 has a significantly smaller number of timepoints, the USE mode is the more recommended choice over the INITIALIZE mode. The ESTIMATE mode is not recommended at all because more single subjected dominated components will show up.

### 3.3 Visualization of the latent features and interpretation

What did the autoencoders learn? Did the network acquire domain knowledge, and how might it function with unseen datasets? Can it produce similar features for visually similar spatial maps? To delve into these learned latent features, we visualized low-dimensional projections of individual tICA spatial maps for each subject, and visualized group cortical maps from selected clusters. After training the volume and surface autoencoders and affirming the best models using validation datasets according to the reconstruction loss, latent features were inferred for each subject’s individual tICA spatial maps across all tICA datasets. These latent features, when concatenated along the component axis (sample axis), were then reduced from their high-dimensional space (162 for surface and 252 for volume) to a two-dimensional plane using 2D t-distributed stochastic neighbor embedding (t-SNE; (Van der Maaten & Hinton, 2008)).

We highlighted three evaluation datasets to illustrate their low-dimensional projections: 1) the HCP-YA 3T resting state fMRI tICA dataset with 535 subjects and 78 group tICA components (Fig 8A) the HCP-D 3T concatenated fMRI tICA datasets with 100 subjects and 78 group tICA components initialized from the first dataset (Fig 8B); 3) the HCP-D 3T concatenated fMRI tICA datasets with 100 subjects and 111 group tICA components (Fig 8C). Additional t-SNE plots for other evaluation datasets can be found in supplementary materials Fig S5 and Fig S6. Each point on the t-SNE plot signifies an individual tICA component per subject, color-coded by its corresponding group component index. A distinct cluster mirrors its respective group tICA component. Surface and volume latent features are shown on the left and right columns of the t-SNE plots, which maintain consistent scales. Notably, surface latent features exhibited clearer dividing boundaries than volume latent features. The former’s clusters are more defined. This phenomenon is emphasized when the group tICA decomposition is applied on a smaller dataset, where the single subject effect is increased leading to more individualized patterns that are not commonly shared across the group. Comparisons between the t-SNE plots in INITIALIZE (row#2) and ESTIMATE modes (row #3) on the same small dataset with 100 subjects reveal superior separation in the former, especially when looking at the volume latent features, indicating the effectiveness of the initialization approach. The surface latent features are spread widely in both modes, suggesting the generalized ability of surface autoencoders.

**Figure 8.**
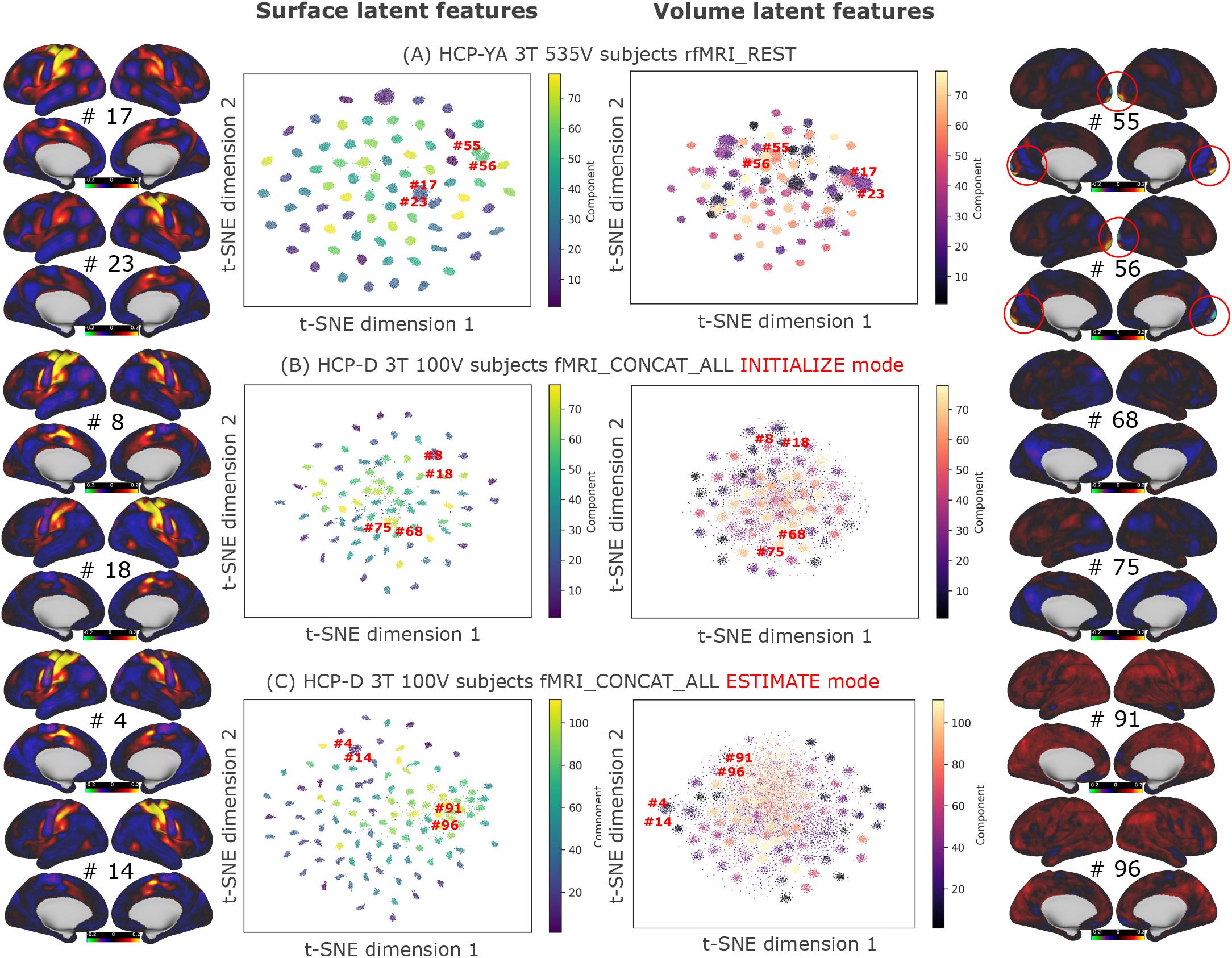
plots show the two-dimensional projection of the learned surface latent features and volume latent features using t-SNE (Van der Maaten & Hinton, 2008). Each point corresponds to one individual tICA component from one subject. Coloring is based on the group tICA component index. Two pairs of the example group components and their corresponding low dimensional location are shown for each t-SNE plot with the same scale. The left column shows somatotopic sensori-motor networks (signal component; Upper: Right hand, Bottom: Left hand), indicating the hemispheric invariance of learned surface latent features. The right column shows 1) Upper: visual signal components (see the red circled areas), 2) Middle: weak components, 3) Bottom: single subject respiration components.

Fig 8 shows example pairs of closely positioned clusters within the t-SNE plots. For instance, the exemplar surface latent features (left column), signal group components with somatotopic sensori-motor networks like #17 and #23 (row #1), #8 and #18 (row #2), and #4 and #14 (row #3) are adjacent, indicating hemispheric invariance – achieved by averaging features from both hemispheres. Volume latent features (right column) display similar hemisphere invariance, a result of the translation invariance characteristic of 3D convolutional kernels and the symmetric pattern in a large number of spatial maps. Other visually similar paired examples include visual signal components #55 and #56 (row #1), weak components #68 and #75 (row #2), and single-subject respiration components #91 and #96 (row #3).

Overall, the two-dimensional t-SNE visualizations of the latent features indicate that the autoencoders effectively extracted visual and perceptual knowledge of tICA components at both group and individual levels. Features that appear visually congruent are projected in close proximity, allowing distinct group components to be clearly differentiated, thus contributing to an interpretable classification.

### 3.4 Effect of feature types

Given the multiple types of features, several combinations of the feature settings should be validated to determine which setting should be considered to be the final one. Visual inspection of the projected latent features revealed that surface latent features generally were more distinct than their volume counterparts, raising the question of whether volume latent features contribute significantly to performance. The handcrafted features cover a wide range of data types focusing on both group and individual level characteristics. Hence, we sought to assess how the feature settings influence classifier performance, namely, 1) volume latent features, 2) surface latent features, 3) combined latent features, 4) handcrafted features and 5) combined latent and handcrafted features. The single XGBoost, one of the tree-based models in the candidate pool, and a logistic regression (the simplest linear model for binary classification) were trained with multiple feature settings using the eight tICA training datasets. Each was exposed to the full set of 100 locally augmented datapoints for each group component, resulting in 75,750 training samples. These two learners were subsequently validated on two tICA validation datasets, excluding the weak component categories to concentrate on performance of the most critical categories. This training process was replicated across five random seeds to account for random variability.

Performance was assessed using three threshold-free metrics: ROC-AUC, PR-AUC, NPV-Spec AUC, resulting in the three columns of subplots shown in Fig 9. The error bars indicate 95% confidence level.

**Figure 9.**
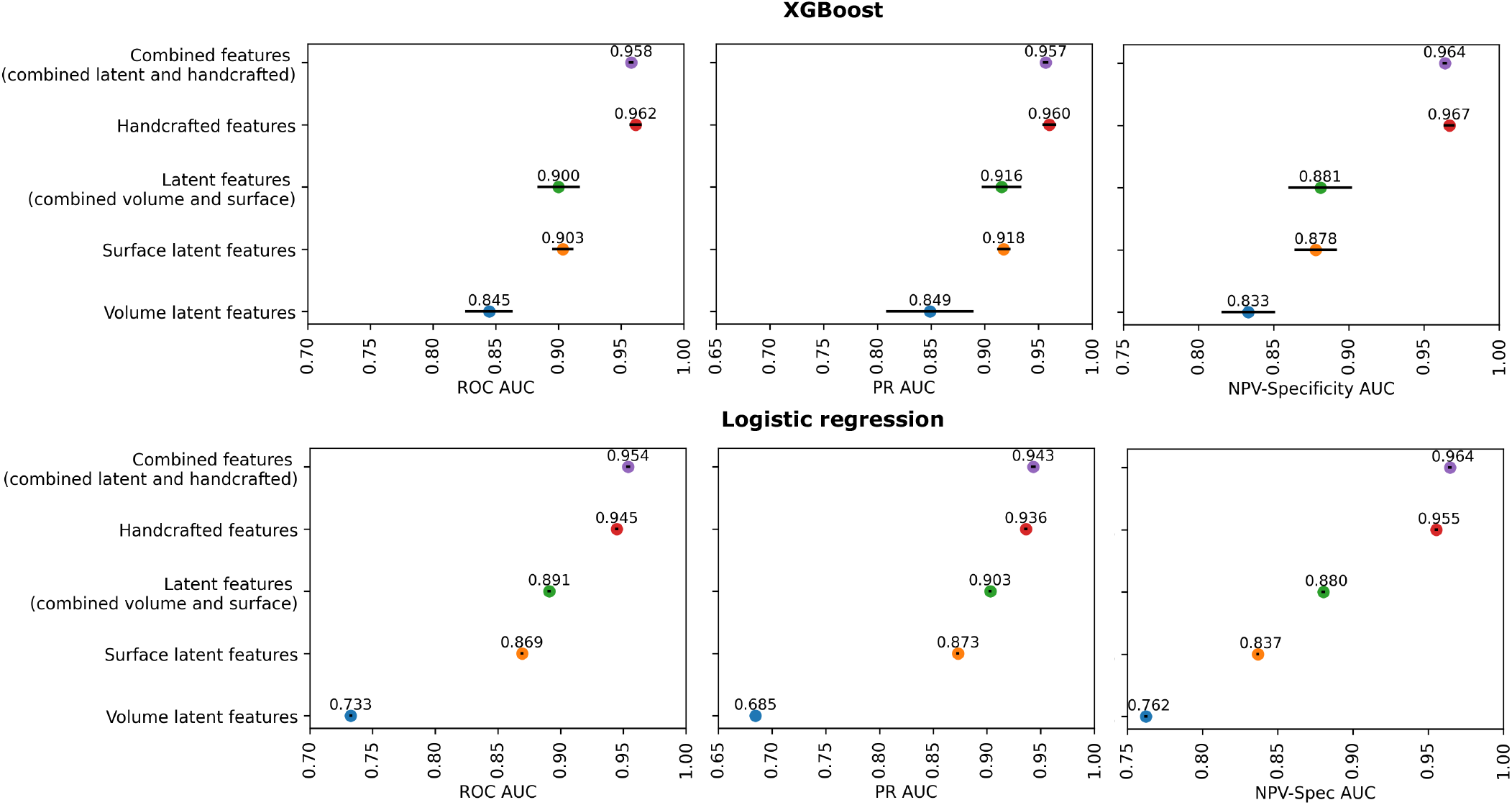
Effects of feature types. Ablation results on the validation sets of HCP-D with 100 and 160 subjects. The plots report the three threshold-free metrics: ROC AUC, PR AUC and NPV-Specificity AUC, for a single XGBoost model (upper) and a logistic regression (lower) trained on feature settings based on volume latent features (blue), surface latent features (orange), the combined latent features (green), handcrafted features (red), and the combined handcrafted and latent features (purple). Error bars indicate 95 % confidence interval.

The volume latent feature setting consistently resulted in the worst performance across metrics and the two learners. Meanwhile, performances of the surface-only and combined latent feature settings were quite similar for the XGBoost classifier, and the combined latent feature setting was slightly better for the logistic regression. The handcrafted feature setting consistently outperformed the latent features under all the three latent feature settings. The performance of combined handcrafted and latent features was comparable to the handcrafted feature setting under XGBoost classifier, but for the logistic regression, the combined feature setting outperformed the handcrafted feature settings.

The gains evident from the combined latent feature setup for logistic regression suggest that volume and surface features can cooperatively illuminate comprehensive patterns in the spatial domain of tICA group components. Notably, the k-NN method emerged as a top performer in balanced accuracy comparisons for surface latent features, reinforcing the interpretability and generalizability of the acquired surface latent features (See Fig S7). In practical terms, this implies that, even without the use of parametric models, solely comparing similarity on surface latent features across datasets can yield excellent performance for tICA component classification. This level of consistency across the training and validation datasets provides a robust basis for subsequent analyses.

Handcrafted features based on domain knowledge performed better than the latent features, suggesting the spatial and temporal domain should be considered together to determine the component category, which agrees with what we found in the manual classification process. The combination of these two feature settings shows limited performance gains in the validation datasets, and sometimes the combination setting performs slightly worse (e.g., in XGBoost) which might hint at potential over-complexity or redundancy when merging the features. Nevertheless, we opted for the combined setting in the final model. This was to ensure all derived features were incorporated, preventing any potentially useful features from being omitted. We then relied on the ensemble model, composed of base learners with diverse capabilities, to discern feature importance and optimize for the best performance.

### 3.5 Effects of data augmentation

The proposed pseudo averaged group data augmentation can dramatically expand the training datasets by randomly sampling a given number of subjects in the group and then averaging their spatial maps to form pseudo-average group spatial maps. To validate its effectiveness and how different levels of augmentation affect the classifier performance, we trained the two selected base learners, XGBoost and logistic regression on the 1) handcrafted features, 2) combined latent features and 3) the combined features with a list of increasing augmentation numbers (0, 20, 40, 60, 80, 100), which indicates the number of randomly sampled subjects to form the pseudo group. The same validation datasets from section 3.4 were used to evaluate the performance. The ROC-AUC was chosen to comprehensively compare the performances across five random experiments. Moreover, an experiment to validate this data augmentation method was performed to get a sense of how similar the pseudo-group spatial maps are when compared to the raw group spatial maps with an increasing number of subjects as shown in Fig S4.

Fig 10 shows the effect of data augmentation on XGBoost (upper) and logistic regression (lower), with the two base learners in the candidate pool and part of the final hierarchical classifier. A substantial performance gain on ROC-AUC can be observed from utilizing the augmentation factor of 20 (20 pseudo-groups per original group) compared with no augmentation for both learners across the feature settings, indicating the effectiveness of using the data augmentation. For the handcrafted features (column #1), XGBoost has a clear performance gain when the augmentation factor is increased from 20 to 100, however, logistic regression declines in performance as augmentation factor increases, perhaps because its simple linear fitting capability fails to capture complex patterns and interactions in the handcrafted features with more datapoints. For latent features, performance gain is unobvious between any nonzero augmentation factor for XGBoost or for logistic regression, which suggests that a substantial amount of augmented training vectors do not help the latent features. For the combined features, no clear performance gain is observed from increasing augmentation factor from 20 to 100 for XGBoost but a clear performance gain is observed for the logistic regression. Note that the combined feature setting for logistic regression has the best performance, better than the handcrafted features, matching the finding discussed in 3.4. Furthermore, a distinct trend emerges for logistic regression with combined features, deviating from both handcrafted and latent features. While handcrafted features exhibit a decrease in performance with increasing augmented samples, and latent features show no clear improvement from an augmentation factor of 20 to 100, the combined feature setting reveals a cooperative effect of features from both sources, resulting in enhanced performance for an augmentation factor of 100 vs 20.

**Figure 10.**
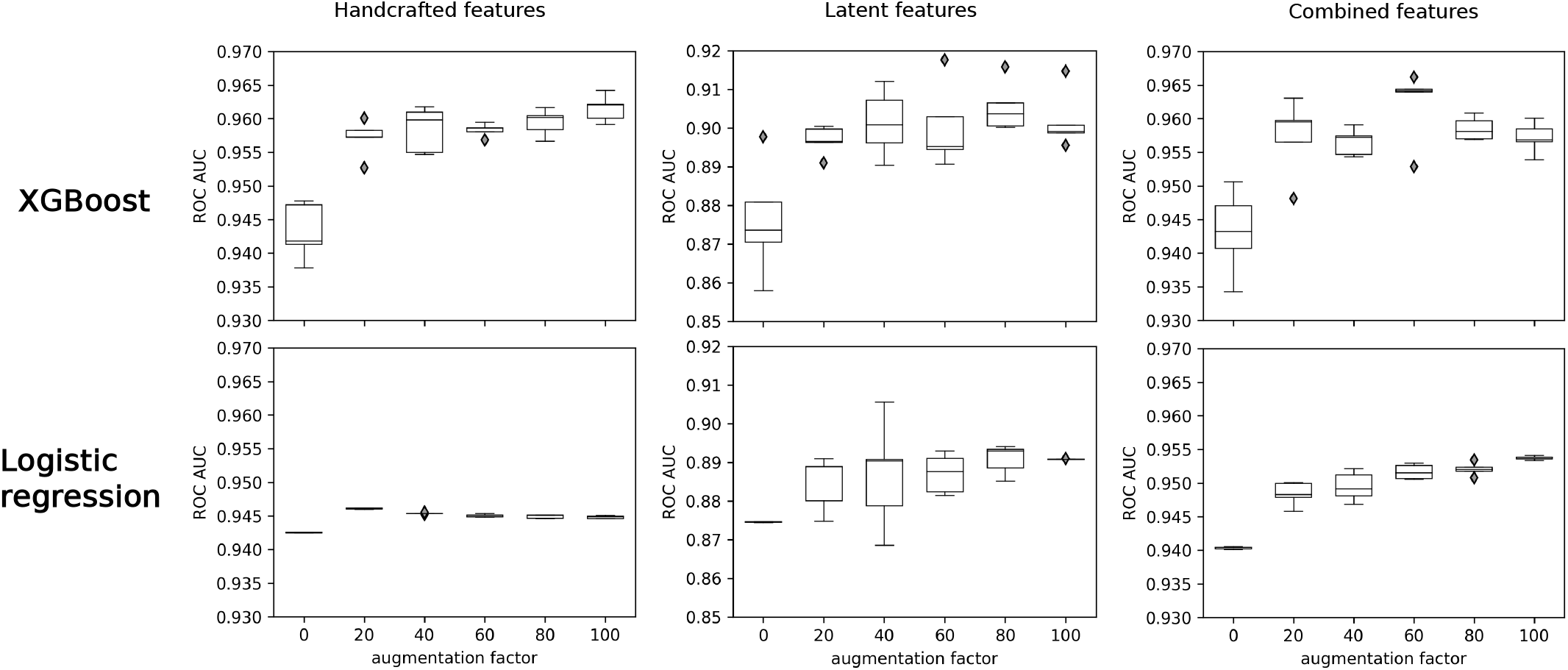
Effects of data augmentation. Ablation results on the validation sets of HCP-D with 100 and 160 subjects. The plots report the ROC AUC for a single XGBoost model (upper) and a logistic regression (lower) trained on feature settings based on handcrafted features (left), combined latent features (middle), and combined handcrafted and latent features (right).

### 3.7 Benchmark comparison using the hierarchical classifier

To test the handcrafted features, latent features and the combined features using the proposed hierarchical classifier, we trained the hierarchical classifier on the full ten training datasets and then evaluated it on the full eleven evaluation datasets, using ROC-AUC, PR-AUC, NPV-Spec AUC and balanced accuracy. The classifier was trained five times using different random seeds, and the median metric from the five experiments is reported in Fig 11. Two evaluation strategies were applied: 1) the most critical components including clear signals or artifacts but excluding the weak components and the 7T volume distortion related components; (Fig 11 upper) 2) no exclusion on the evaluation datasets (Fig 11 lower). For both evaluation strategies, the hierarchical classifier trained on the combined feature setting outperformed the rest in two metrics: ROC-AUC, PR-AUC, suggesting it as the overall best performance setting, while for NPV-Spec AUC, the latent feature setting from the evaluation strategy #1 outperformed the rest, indicating that the latent features are more able to correctly identify the clearly artifactual components. For balanced accuracy, the hierarchical classifier trained on the combined feature settings outperformed the rest when evaluated on the most critical components, whereas it performed slightly worse than the handcrafted feature setting for all the components without any exclusion. The performance of the latent feature setting dropped significantly when evaluated on the full evaluation datasets without exclusion. This setting only uses spatial information which, for example, is not sufficient to distinguish the artifactual components with volume distortion in the 7T tICA datasets, which are usually visually similar to signal components. The base learners both performed worse than the hierarchical classifier across the evaluation strategies and metrics, indicating that the ensemble learning method, even with extra complexity, improves performance for tICA component classification.

**Figure 11.**
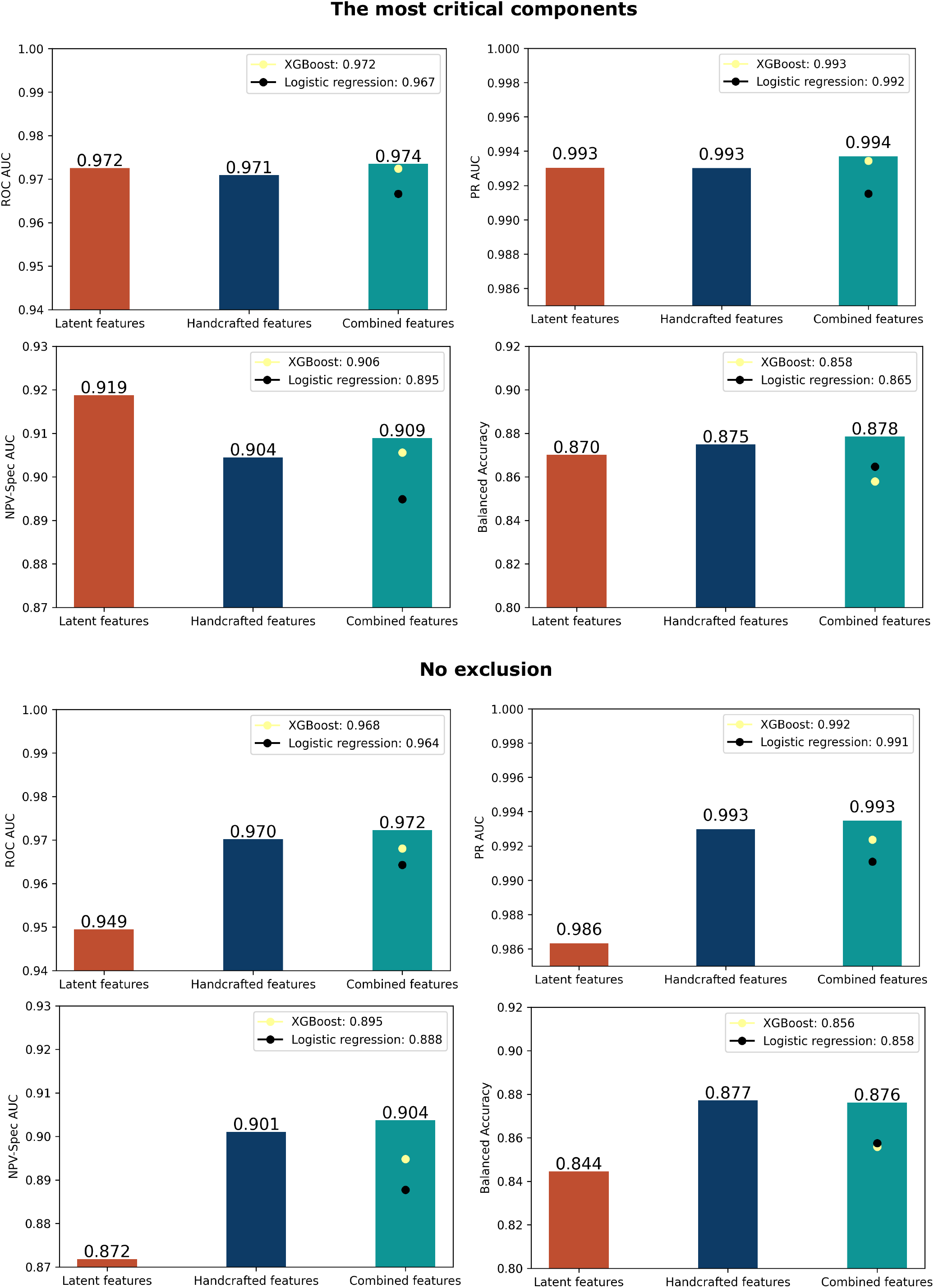
compares the evaluation performance on the full eleven evaluation datasets 1) with the most critical components (Upper), 2) without any exclusion (Lower), under the three feature settings, 1) latent features (dark orange), 2) handcrafted features (dark blue) and 3) the combination of the two (blue-green). The performances of the base learners XGBoost and logistic regression using the combined feature setting are shown as light-yellow circles and black circles, respectively. Each classifier is trained for five random seeds and the median of each evaluation metric is shown. The combined feature setting outperforms the other two feature settings in ROC-AUC and PR-AUC. The handcrafted feature setting has a comparable performance with the latent feature setting when evaluated on the most critical component. However, the latent feature setting performs worse when evaluated on the no-exclusion case, indicating that while spatial features alone are effective in identifying key components, they fall short in accurately classifying weak components or those whose distinguishing characteristics are primarily temporal (e.g., 7T volume distortion artifactual components).

We summarized the performance metrics of the hierarchical classifier with the combined feature setting for each of the eleven evaluation datasets (Table. 4). All datasets have an ROC-AUC larger than 0.9, a PR-AUC over 0.98, and a correct prediction variance percentage over 88%. The sICA-recleaned datasets with sufficient subject size: HCP-Young Adult 3T resting state with 535 subjects (2,568,000 concatenated timepoints), HCP-Young Adult 3T task fMRI with 535 subjects are classified with high performance (2,075,800 concatenated timepoints), and only 2.12% of the variance from the artifactual components is misclassified, and no signal components were misidentified. The sICA-cleaned datasets without further recleaning processing but with sufficient subject size and a wide range of subject ages: HCP-Aging 3T concatenated fMRI with 617 subjects (1,678,857 concatenated timepoints), HCP-Development 3T concatenated fMRI with 314 subjects (1,004,800 concatenated timepoints) are classified with comparable performances with close to 1 ROC-AUC, PR-AUC and NPV-Spec AUC, 93%∼94% of the variances are correctly classified. Performance on the sICA-recleaned 7T datasets is also high despite having fewer subjects. Specifically, for HCP-Young Adult 7T resting state with 87 subjects (317,985 concatenated timepoints), HCP-Young Adult 7T movie task fMRI with 87 subjects (313,200 concatenated timepoints) and HCP-Young Adult 7T retinotopy with 87 subjects (156,600 concatenated timepoints), ROC-AUC were all 0.99, PR-AUC were all 1, NPV-Spec AUC ranges from 0.93 to 0.98, and more than 96% of the total variances of the components was correctly classified. To further evaluate the model’s robustness for datasets with small sample size and different tICA decomposition modes (ESTIMATE: estimate from scratch; INITIALIZE: use a precomputed mixing matrix to initialize the estimation), we analyzed two pairs of tICA datasets from the HCP-Development dataset after sICA recleaning: 60 subjects (192,000 concatenated timepoints) and 100 subjects (320,000 concatenated timepoints) each of which was run through the two tICA modes. Under the ESTIMATE and INITIALIZE modes, the correct prediction variance percentage ranged from 88.6% to 96.8%, suggesting that the majority of the components were correctly classified no matter what mode the dataset is run in. However, for the ESTIMATE mode with 100 subjects, the model has 8.7% of variance from FN predictions, indicating that the model tends to assign a small number of signal components as artifact for this particular dataset. Since no training dataset was processed using a tICA mode other than ESTIMATE, the current high performance on the datasets under INITIALIZE mode suggests good generalizability of the generated features and the hierarchical classifier.

### 3.8 Interpretability of the model

The hierarchical classifier does not have a feature selection step because we wanted each base learner to be exposed to the full range of possible features and to make selections that diversify the model learning capacity. This section focuses on the interpretability of the model and endeavors to answer three questions: 1) does each base learner focus on different features? 2) which features are more important? 3) what does the learned model teach us about the decision-making process? The interpretability was analyzed for XGBoost and for logistic regression (the only linear base learner) on the eleven evaluation datasets, using the SHAP (SHapley Additive exPlanations; (Lundberg & Lee, 2017)) that is based on the concept of Shapley values from cooperative game theory. Both learners were directly taken from the hierarchical classifier trained on the full ten training datasets.

SHAP values distribute the difference between a model’s prediction for a specific instance (a datapoint in the feature set) and the average prediction over all instances among its features, based on their individual contributions derived from Shapley values in cooperative game theory. In Fig 12, we sorted the top 10 features for XGBoost (panel A) and logistic regression (panel B) in descending order according to their instance-averaged magnitudes of the SHAP value. The higher feature value is denoted in red, and the lower feature value is denoted in blue. A SHAP value of zero means that a feature does not contribute to either the signal or the artifactual component prediction. The distribution per feature for the SHAP values are plotted row-wise. A positive SHAP value for a given instance indicates a ‘vote’ that the feature in that instance contributes to signal prediction, whereas a negative SHAP value indicates a ‘vote’ that the feature in that instance contributes to artifact prediction. The farther the SHAP value is from zero, the greater the weight it contributes to the ‘vote’. For XGBoost, the top 10 features consist of one temporal feature and nine spatial features, suggesting that this model captures patterns using both data types. The top 3 features all reflect situations in which an artifact in a single ‘dominating’ subject (the subject that contributes the most of the total variance for a given group component) contributes to an artifact prediction with high standard deviation or mean value of the z-stat map for a particular ROI (left cerebellar, subcortical, or 1st vascular territory). The higher values for these features in the negative direction of the SHAP value axis suggest a stronger negative effect on predicting signal components, hence, they are helpful for predicting artifactual components. A larger standard deviation indicates a wide spread of the ROI intensities, as is common in artifactual components, and a high mean value in a vascular territory suggests an abnormal pattern that is common in artifactual components. The only temporal feature in the top 10 list describes the L2 norm between the dominating subject’s tICA component and itself after the outliers are replaced using nearest neighbor nonoutlier value. The several high values of such a feature fall on the left side of the y axis, indicating that the large L2 norm is an artifactual component. Among the nine spatial features, only one came from the latent features, which is the 59th surface latent feature that tends to have a positive influence on predicting signal components when it increases. The outlierness in the left grey matter region and the vessel region of the group normalized tICA volume map helps to identify signal outliers compared with the rest of the low values in such regions which has a positive impact on predicting signal components. The mean absolute intensity within parcel L_25_ROI is an indicator that can be helpful for identifying artifacts arising from head motion. As illustrated in Fig 12A, higher values of the mean of absolute intensities in this region tend to predict artifact.

**Figure 12.**
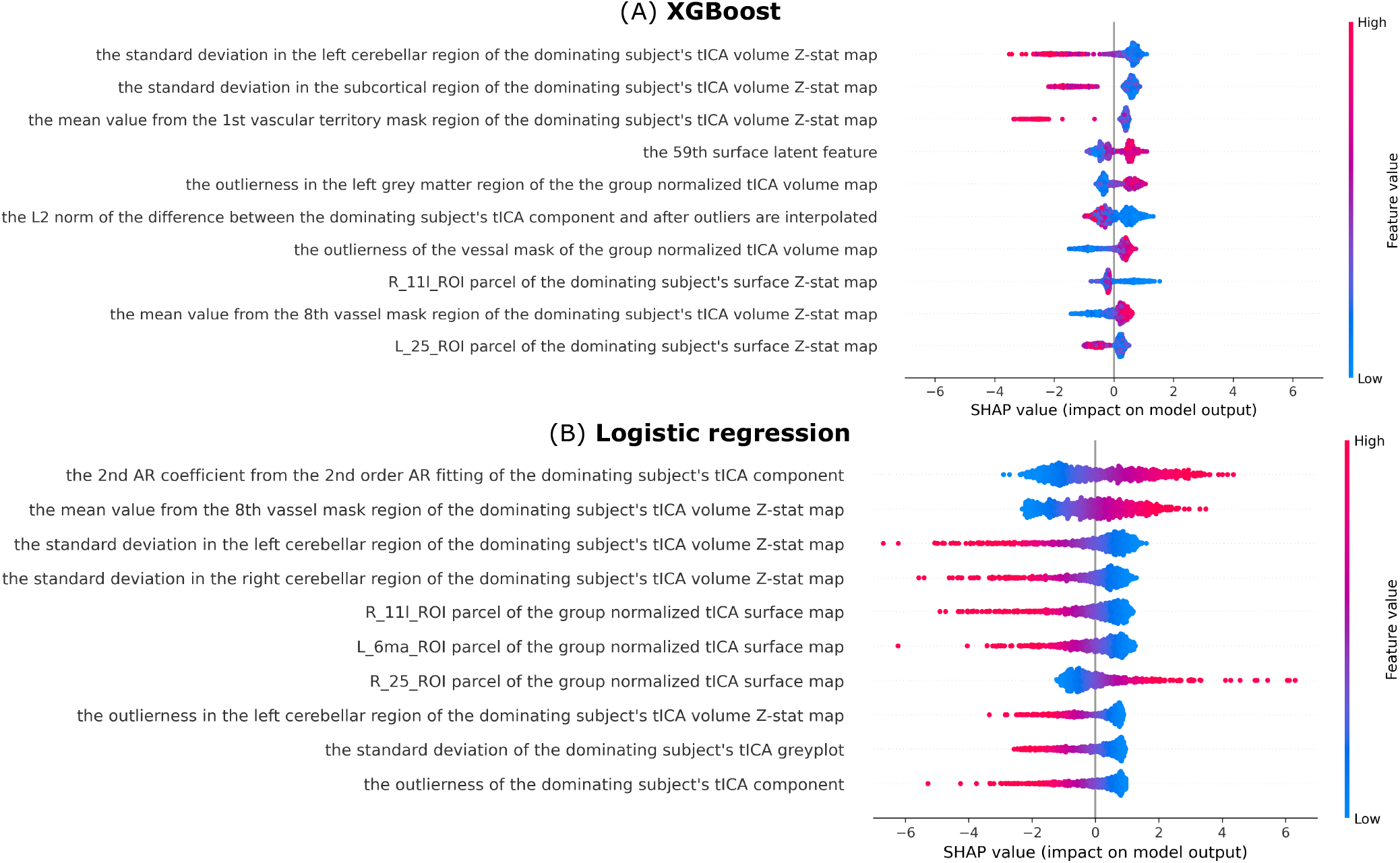
shows the SHAP feature importance values of two base learners from the hierarchical classifier XGBoost (upper) and logistic regression (bottom). The features are ordered by the global feature importance score, computed as the mean absolute SHAP values. Each point corresponds to a certain feature for one instance in the 11 evaluation datasets. A higher feature value is indicated in red, and a lower feature value is indicated in blue. A positive SHAP value for a feature indicates that the feature is pushing the model’s prediction towards a signal prediction. A negative SHAP value for a feature indicates that the feature is pushing the model’s prediction towards an artifactual prediction. The majority of the top importance features are spatial features, matching what was found in the manual classification process.

In the Fig 12B plot of logistic regression, the top 10 features include two temporal features, one spatial-temporal feature from the greyplot and seven spatial features. The top feature is one of the AR coefficients for the dominating subject’s tICA component (timeseries), indicating an autocorrelation effect on the dominating subject’s tICA timeseries tends to make a group component more signal-like. The third and fourth features are the standard deviation in the left and right cerebellar region of the dominating subject’s tICA volume Z-stat map, which suggests that the increase of the standard deviation in those regions tends to have a positive impact on identifying artifactual components. Fig S9 shows similar results for the odds ratio for each of the top 4 features. The other temporal feature is the outlierness of the dominating subject’s tICA component; as the SHAP value demonstrated, the higher value of outlierness the stronger positive effect on predicting artifactual components. The only spatial-temporal feature in the top 10 is the standard deviation of the subject’s tICA greyplot, and for this also the higher value leads to a stronger effect on predicting artifactual components. The parcel R_25_ROI in logistic regression have a contrary distribution compared with its symmetric counterpart L_25_ROI in XGBoost, which could indicate that this parcel-based feature has a non-symmetric effect on making the decision. The R_11l_ROI has the matched SHAP value distribution in both XGBoost and logistic regression, where the larger value suggests an artifactual component. Note that, the standard deviation in the left cerebellar region of the dominating subject’s tICA volume Z-stat map shows up under both base learners, suggesting the effectiveness of the handcrafted features that is demonstrated by both classifiers. The largely non-overlapping features indicate that different base learners can focus on different aspects of the features and thus can cooperatively lead to a robust final decision.

## 4. Discussion

This study introduces a fully automated temporal ICA pipeline for HCP-style datasets to clean globally or semi-globally structured artifacts associated with respiration or other physiological phenomena. Compared to the initial tICA pipeline, this entailed estimation of an appropriate number of Wishart distributions to MIGP data for automated group sICA dimension estimation in CIFTI space from a PCA series, and adding a hierarchical classifier to distinguish signal vs artifactual components using handcrafted features from domain knowledge plus automatically captured latent features via self-supervised learning. Given the limited number of group tICA components, two strategies were designed to leverage the large amount of subject-level information: 1) self-learning on subject-level spatial maps to obtain individual level features; 2) pseudo group averaging to increase the number of non-identical group components. Using both methods, we demonstrated a good overall performance of the classifier. We defined three variance percentage-based metrics to evaluate classifier performance based on the importance of each group component quantified by its percentage of the total variance: 1) correct prediction variance percentage covering TP and TN predictions, 2) FN prediction percentage and 3) FP prediction percentage. We demonstrated the effectiveness and generalizability of the hierarchical classifier on datasets in several contexts 1) completeness of sICA denoising: sICA cleaned and sICA recleaned with an improved classification; 2) data acquisition specifics: 3T vs 7T scanner; 3) fMRI characteristics: resting state vs movie vs retinotopy vs task-fMRI (working memory, language processing, relational processing, emotion processing, guessing, Go/NoGo task, vismotor, and facename tasks); 4) tICA mode: ESTIMATE, INITIALIZE; 5) group size: 60, 87 (7T data), 100, 314, 535, 617. Previous work (Glasser et al., 2018, 2019) has demonstrated the advantages of tICA cleanup when compared to global signal regression (GSR) following sICA+FIX and to sICA+FIX alone. Specifically, tICA cleanup effectively eliminates the residual uniform global artifact that is unable to be cleaned by sICA+FIX. Global respiratory artifact results in a global positive bias in functional connectivity, which can vary between groups in a study. In contrast, GSR not only shifts functional connectivity gradients but also introduces a network-specific negative bias in both task fMRI and resting-state fMRI connectivity and activity, detrimentally impacting downstream analyses. The present study primarily focuses on the following objectives: 1) simplifying the use of tICA cleanup (e.g., minimizing the need for manual interventions), 2) generalizing the applicability of tICA cleanup to a wide range of HCP-Style dataset settings (e.g., various group sizes), 3) addressing potential methodological limitations associated with tICA cleanup (e.g., the single subject effect within group tICA), 4) providing guidelines to assist the community in effectively using the tICA pipeline (e.g., recommendations regarding the minimum number of timepoints required for different tICA modes and the appropriate number of Wishart Distributions to use for a given number of MIGP PCA components). The resultant automated tICA pipeline, as part of the HCP pipelines, enables investigators to remove physiology-related artifacts from HCP-style datasets with minimal manual intervention.

### 4.1 The number of Wishart Distributions

A general objective is to estimate the largest number of reproducible sICA components given a particular number of MIGP PCA components and to set the number of Wishart Distributions accordingly. The first few Wishart Distributions mainly fit unstructured noise, but using too many reduces the signals of interest and reduces sICA dimensionality (Fig 5). Note that when too many Wishart distributions are used, the reproducibility tends to artificially increase, due to the reduced degrees of freedom as fewer ICA components are found. In setting a reasonable range for sICA dimensionality, we relied on an earlier analysis that used values of d=84 for 3T resting state and d=70 for 3T task in the HCP-YA dataset (Glasser et al., 2018) as a reference point based on the number of reproducible component clusters having ICASSO Iq value greater than 0.5. Subsequently, we conducted manual inspections to assess oversplitting and undersplitting effects on known network patterns within the resulting group spatial ICA components. We determined the optimal number for the large-scale HCP-YA datasets across various fMRI tasks while considering a cluster index Iq threshold above 0.85 as indicative of better reproducibility. Our recommended number of six Wishart Distributions, identified from the 3T HCP-YA dataset, was directly applied to the Lifespan HCP-Aging and HCP-Development datasets without the need for re-estimation. This yielded dimensionalities of 86 and 122, respectively, following group sICA. Notably, HCP-Development (ages 5-21) included subjects with substantial head motion, resulting in a larger number of components. Children might also have a larger number of reproducible functional networks. For datasets having smaller numbers of MIGP PCA series timepoints, five Wishart Distributions may be more appropriate for estimating dimensionality (see Fig S2 for the relation between the number of MIGP PCA, the WD and resulting sICA dimensionality and reproducibility).

### 4.2 The tICA decomposition modes

Temporal ICA has a tendency to identify single subject components when the number of subjects and timepoints is small, allowing the super Gaussianity of individuals with distinct temporal patterns to split into separate components. These individualized components result in less accurate global artifact cleanup. This single-subject phenomenon decreases as the number of subjects increases because each subject contributes less to the overall variance of the data (see section 3.2 *Effect of the number of subjects and total timepoints on the single subject effect*). We did not see this trend for HCP 7T subjects as shown in Fig S3 because there were fewer HCP 7T subjects and the effects of differential T2* patterns across subjects are enhanced at 7T. These patterns are induced by the unique b0 field inhomogeneities of each subject and the unique folding patterns relative to the main magnetic field. More generally, it is commonly impractical to collect hundreds or thousands of subjects in conventional studies, and many use cases for tICA cleanup may involve only dozens of subjects. To make the tICA pipeline applicable to small datasets that would be strongly impacted by single subject effects if the decomposition were estimated from scratch, we introduced the INITIALIZE and USE decomposition modes that take advantage of precomputed group sICA and tICA mixing matrices from the HCP high-quality datasets having thousands of subjects. These modes provide a way to substantially reduce the single subject effect while enabling users to choose between initialization with a reasonable starting point and direct use of a prior decomposition. Table S1 shows that the incidence of single-subject-dominated components dropped from 66.1% (80 out of 121 total components) to 3.8% (3 out of 78 total components) for HCP-D with 60 subjects, from 46.8% (52 out of 111 total components) to 1.3% (1 out of 78 total components) for HCP-D with 100 subjects using the INITIALIZE mode. The INITIALIZE mode allows for adaptation to the study without the problematic single subject behavior. The hierarchical classifier, even if not trained with any initialized datasets, can successfully identify 92-93% of total variance in the initialized evaluation datasets, which is generalizable to real world applications. Note that the officially released classifier was trained on all 21 datasets in the train-valid-test splits (Table 1) to cover ESTIMATE and INITIALIZE modes. However, the evaluation conducted in this study was using the classifier trained on only the training and validation datasets (Table 1), specifically to demonstrate the effectiveness of the methodology.

To provide investigators with guidelines for selecting tICA modes, we analyzed the relationship between the number of concatenated timepoints and the percentage of single-subject dominated components within a fixed dimension of group components, as estimated through large-scale group spatial ICA. Utilizing a tree model with monotonically decreasing constraints, we derived a stepwise non-increasing curve that offers valuable insights into optimal timepoint recommendations (Figs 6 and 7). We determined a practical threshold of 7%, allowing for a range of 5 to 9 single-subject components within the group tICA datasets, where the total number of group components ranged from 82 to 122. This threshold guided the identification of a plateau, resulting in recommended numbers of concatenated timepoints of 620k, 480k, and 1 million for HCP-YA task fMRI, HCP-A, and HCP-D concatenated fMRI datasets, respectively, corresponding to 170, 176, and 312 subjects.

Importantly, the HCP-D dataset, characterized by a significant degree of subject movement and the potential for distinct temporal patterns, displayed a higher susceptibility to single-subject effects. Conversely, HCP-A exhibited a lower requirement for the number of concatenated timepoints to achieve an equivalent percentage of single-subject dominated components. In practical application, the suggested total concatenated timepoints serves as a reference for users, allowing them to make informed decisions regarding the choice of tICA mode, with consideration given to the specifics of their data acquisition parameters and subject inherent characteristics.

### 4.3 Deep learning versus traditional machine learning

It is a long-debated question: which one is better, handcrafted features with traditional machine learning or deep learning without explicit domain knowledge? In this study, we explored both approaches and provide our answer to the question: handcrafted features perform better in tICA classification, and combining the two methods is preferred. tICA component classification focuses on identifying the signal or artifact category of group-level tICA components. We chose self-supervised deep learning at the individual tICA component level for two reasons: 1) The number of group components is limited, usually 60 to 130 for one group dataset and directly training supervised deep learning models on datasets with small sample size may suffer from overfitting. 2) self-supervised deep learning doesn’t require labels and can be directly applied to individual tICA spatial maps. Given the rich information carried by spatial maps, we implemented volume and surface autoencoders that first learn from individual spatial maps and obtain group features by averaging across the individual learned features, where the number of available samples multiplies with the group size (e.g., HCP 3T resting state training dataset has 84 group components and 535 subjects, then the total number of individual spatial maps becomes 84*535=44,940). We conducted experiments which showed that combination between surface and volume latent features achieves better performance compared with single source (however, between the two, the surface-based features were clearly superior to the volume-based ones), then we found that the handcrafted feature setting outperforms the latent feature setting when evaluated on all the components without exclusion (Fig 11 lower), but they have comparable performance when evaluated on the components that are not borderline calls. This result suggests that 1) spatial information alone is unable to identify the borderline components; 2) the latent features are helpful to distinguish clear signal and artifacts; 3) the handcrafted features provide a more comprehensive representation of group tICA components. Domain knowledge for classifying the tICA components plays a very important role, which covers multiple data sources from volume, surface, timeseries, power spectrum and greyplot. Spatial autoencoders, on the other hand, only focus on volumes and surfaces, and they ignore temporal and spatial-temporal patterns. Such an observation was evident again in spatial ICA classification: after training on 47,985 HCP-YA 7T retinotopy sICA components, we found that FIX, designed as a hierarchical classifier with handcrafted features, outperforms the deep learning based classification models in balanced accuracy and the (3*TPR+TNR)/4 metrics (see Fig S10): 1) volume based ResNet (He et al., 2016); 2) surface based Vision Transformer (Dahan, Fawaz, et al., 2022); 3) a naïve fusion Vision Transformer (Dosovitskiy et al., 2020) where volume, surface, timeseries and power spectrum are divided into patches and are concatenated as a single sequence as the input. This bolstered our conclusion about the tICA classification that the domain knowledge from multiple data sources is not easily captured by deep learning models, even with sufficient training datapoints and a more advanced model architecture (e.g., Transformer). Multimodal deep learning is still an unsolved problem in the deep learning community. As the final setting, we combined latent features and handcrafted features and found it brought consistently better performance on the evaluation datasets, therefore in this case the combination between deep learning and traditional machine learning achieves better performance.

### 4.4 Limitations and future work

There are several limitations to this study. The tICA cleanup as initially introduced (Glasser et al., 2018, 2019) was only intended for use in large-scale HCP-Style neuroimaging studies because 1) tICA cleanup requires sICA+FIX cleaned dense timeseries as input and this method does not perform well on datasets with low spatial and temporal resolution and a modest number of timepoints (Griffanti et al., 2014) and 2) tICA decomposition itself has even more stringent requirements (Smith et al., 2012). The improved tICA pipeline introduced here follows the same rationale but has been extended to be useful for smaller datasets that meet the HCP-Style requirements, by using the different tICA modes. We illustrated the INITIALIZE mode in an HCP-Style clinical setting in (Lamichhane et al., 2021), demonstrating its effectiveness in HCP-Style dataset with a smaller group size and lower spatial resolution than the existing large high-quality HCP datasets used in this study.

Unlike the surface autoencoder that is trained on the full mesh resolution of the cortical map, the volume autoencoder learns from volume maps that have been downsampled to 4mm, due to the GPU memory limitations and full-FOV representation. This presumably loses detailed and valuable spatial information and may account in part for the dramatically lower performance of the volume-based features relative to the surface-based features. Given the current low-dimensional separation provided by volume latent features as shown in t-SNE plots from Fig 8 and Fig S6, the resampling doesn’t affect the goal of generating well-separated latent representations of tICA individual volume maps at the group level (volume latent features can reasonably separate group tICA components). However, if the goal is to reconstruct the full resolution volume maps, it should be recommended to include no downsampling of the data. Moreover, given the comparable validation performances of combined latent features and surface latent features, we recommend paying more attention to the surface-based analysis if users wish to train the classifier from scratch and save time and computation resources.

As with any other supervised learning model, the tICA classifier is trained on expert labeling before being used in practice. Similar to the sICA classification, labeling can be skewed amongst experts due to varying levels of confidence in specific components that are combined with neural signals and artifacts. A workaround strategy is to involve multiple expert labelers and then average the labels in order to reduce the labeling bias. However, since tICA classification is new to the community and there are currently only a few available experts, we chose another strategy, evaluating performance on 1) components with obvious labeling only; 2) all the components including borderline components that may be classified differently by human experts, which ensures the model’s practical utility and dependability. On the other hand, given the high AUC in evaluation performances, adjusting the threshold allows for reasonable control of the classifier’s confidence level in terms of signals or artifacts, depending on the aggressiveness of the cleaning process.

The autoencoder utilized spatial information only. With more advanced models such as transformers, it is feasible to train a model that takes all the different data sources for each individual tICA component as a sequence of patches, learns the relations across patches from different sources, and reconstructs each of them concurrently. The learned individual-level embedding can be similarly averaged across the group to obtain the group embedding. However, the canonical transformer model (Vaswani et al., 2017) has quadratic complexity in terms of the input sequence length which requires a GPU with large memory. Recent studies have shown promising results in reducing the complexity (Dao et al., 2022; Jaegle et al., 2021; Wang et al., 2020; Xiong et al., 2021), making this a topic for future exploration.

We chose to include all the designed and derived features to train the model by taking advantage of an ensemble architecture, but this requires redundant information that can be potentially excluded from the feature set to reduce the complexity. Besides traditional filter and wrapper methods, a self-attention mechanism that calculates the feature importance scores during training, makes sure that the model learns the relationships between features, and identifies the most important features would be helpful. We consider exploration of feature selection strategies another important topic for future work.

Additionally, future studies should focus on benchmarking more models using more tICA datasets with various settings, and performing further ablation studies on the features to help elucidate important features that result in performance improvements.

## 5. Conclusions

We have demonstrated a fully automated temporal ICA pipeline to clean the globally structured artifacts in HCP-style datasets, where the automation is achieved by 1) estimating the sICA dimensionality by first reducing random noise from MIGP data using multiple Wishart distributions; 2) a hierarchical classifier (of tICA components) equipped with handcrafted features from domain knowledge, and spatial latent features from autoencoders. Three tICA modes were introduced to enable tICA cleanup to datasets with a wide range of subject numbers while avoiding excessive single subject components. We demonstrated that our sICA dimensionality estimation gave a reasonable result on the MIGP data comparable to prior manual estimates using a different method in the same data, and that the tICA components can be correctly classified from a wide range of HCP-style datasets with different characteristics including number and type of fMRI runs, acquisition settings, and type of sICA cleaning. To our knowledge, this is the first method to enable automatic classification of temporal ICA components in fMRI data. The tICA pipeline has been integrated into the HCP pipelines to let users clean respiration and other temporal artifacts from their HCP-style datasets with minimal human intervention. We will release tICA cleaned data for HCP-Young Adult, HCP-Aging, HCP-Development simultaneously.

## Acknowledgments

Supported in part by HCP-YA: Mapping the Human Connectome: Structure, Function, and Heritability U54MH091657; HCP-D: Mapping the Human Connectome During Typical Development U01MH109589; HCP-A: Mapping the Human Connectome During Typical Aging U01AG052564; AABC: Adult Aging Brain Connectome Project U19AG073585; Connectome Coordination Facility (CCF) I: 5R24MH108315; Connectome Coordination Facility II: 1R24MH122820; NIH RO1MH060974 (DCVE, MFG); and by the 14 NIH Institutes and Centers that support the NIH Blueprint for Neuroscience Research and the McDonnell Center for Systems Neuroscience at Washington University (HCP-YA, HCP-D, HCP-A, CCF I, II). Computations were performed using the facilities of the Washington University Research Computing and Informatics Facility, which were partially funded by NIH grants S10OD025200, 1S10RR022984-01A1 and 1S10OD018091-01 and by The McDonnell Center for Systems Neuroscience.

