## Supplementary Material for "Automating the Human Connectome Project’s Temporal ICA Pipeline"

### Introduction to Supplementary Material

The Supplementary Material comprises two sections: 1) Supplementary Results and Figures, which includes the tables and figures referenced in the Main Text, and 2) Supplementary Methods, which covers the sICA reclean process used for the 3.0 official release of HCP-Young Adult and HCP-Lifespan (Aging and Development), as well as the automated sICA reclean pipeline.

### Supplementary Results and Figures

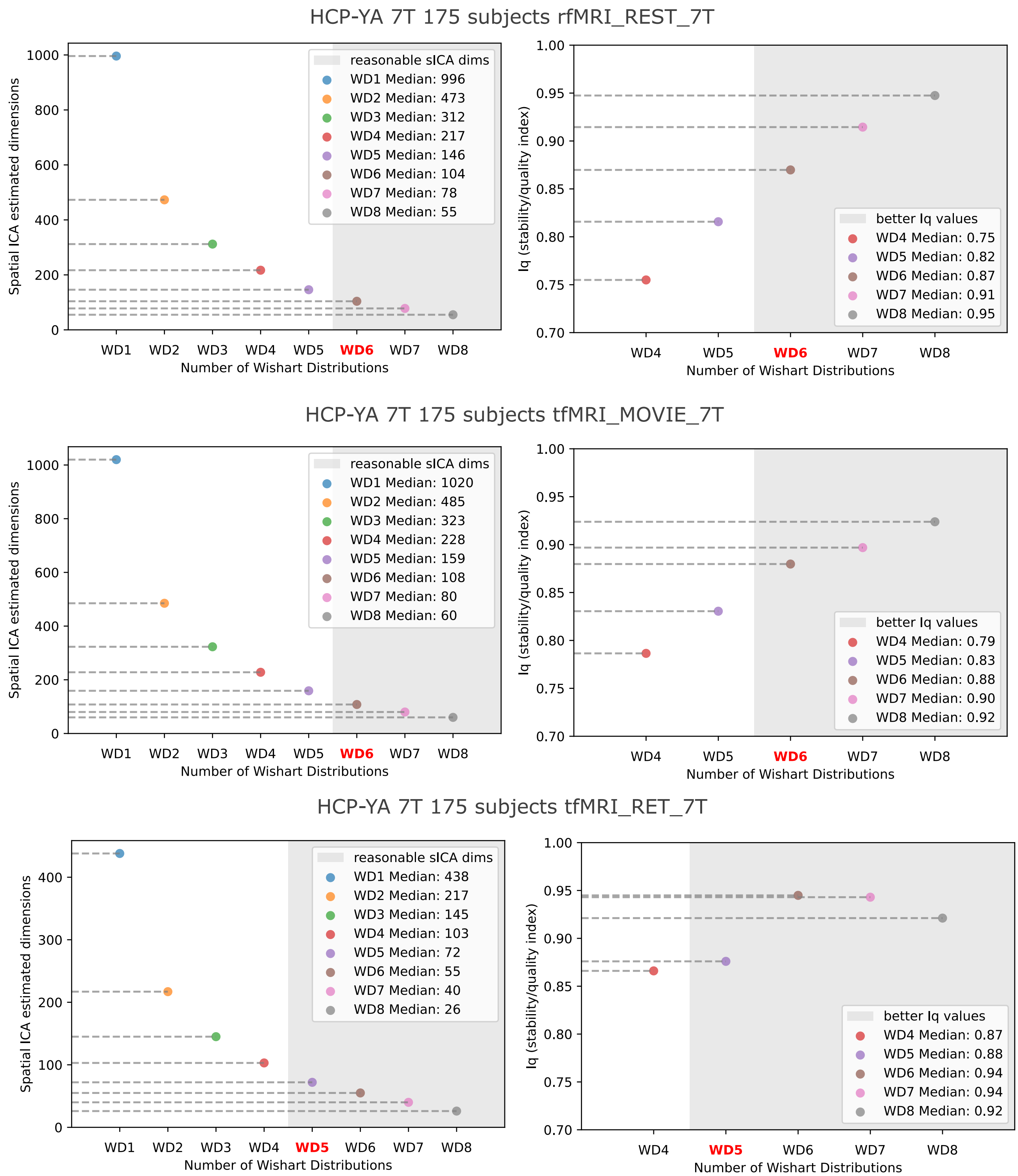


*Supplementary Figure 1* ***Multiple Wishart distributions on HCP-YA 7T datasets.*** *The trends are similar to the 3T HCP-YA in Fig 5. The only difference is that WD5 is chosen for the 7T retinotopy to cover sufficient components.*

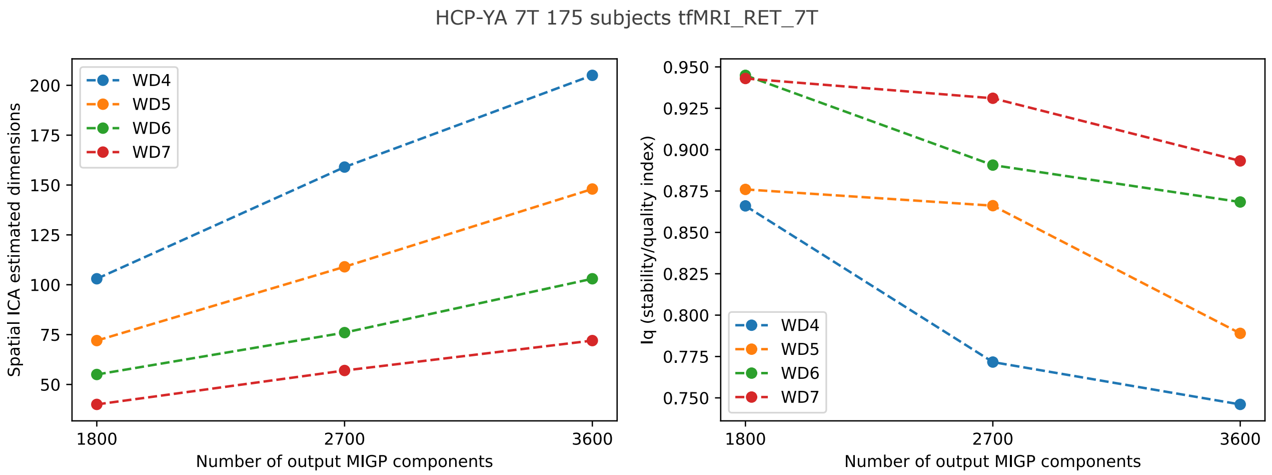


*Supplementary Figure 2* ***Effect of output MIGP components for HCP-YA 7T retinotopy.*** *The median measure from 3 experiments is illustrated as each point in the plots. A larger number of MIGP output components leads to a higher estimated sICA dimensionality. We ended up with the MIGP number of components matching with the expected timepoints, which is consistent across HCP datasets, and WD5 was chosen. When a user attempts to have a smaller number of expected timepoints, one may seek to either reduce the WD to sum up or increase the number of MIGP output components.*

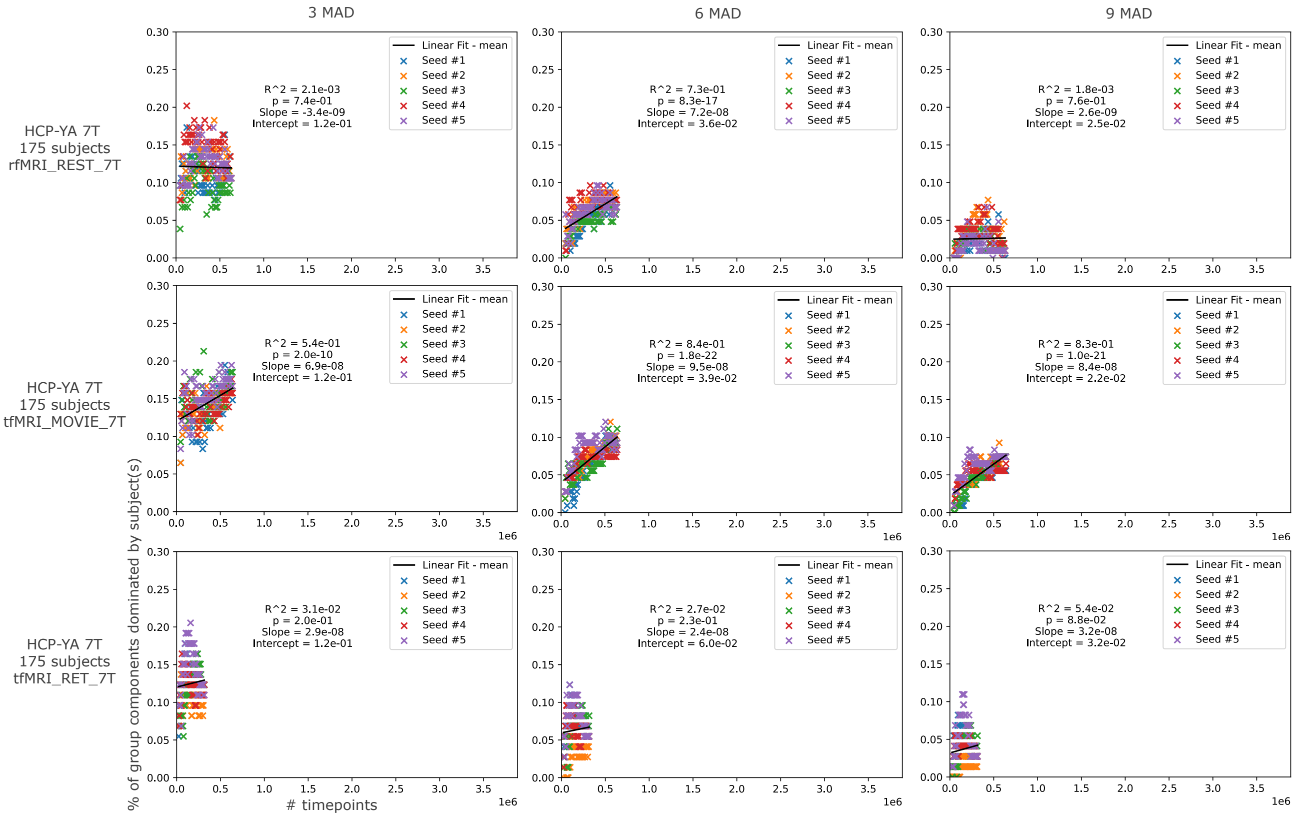


*Supplementary Figure 3* ***The single subject effect in HCP-YA 7T datasets.*** *The more subjects lead to an increasing trend of outlier components. The finding may not reveal the comprehensive pattern due to the limited total number of subjects in HCP-YA 7T datasets (175 compared with HCP-YA 3T 1071, HCP-A 1798, HCP-D 1695).*

| | percentage of single subject dominated components (%) | percentage of non single subject dominated components (%) | # single subject dominated components | tICA dimension |
|---|---|---|---|---|
| S1200_MSMAll3T535T_rfMRI_REST | 0 | 100 | 0 | 84 |
| S1200_MSMAll3T535T_tfMRI_Concat | 0 | 100 | 0 | 70 |
| S1200_MSMAll7T87T_rfMRI_REST_7T | 48.1 | 51.9 | 52 | 108 |
| S1200_MSMAll7T87T_tfMRI_MOVIE_7T | 44.2 | 55.8 | 50 | 113 |
| S1200_MSMAll7T87T_tfMRI_RET_7T | 32 | 68 | 24 | 75 |
| AABC_Version2_Prelim_Data_Visits_617T_fMRI_CONCAT_ALL_clean | 6.2 | 93.8 | 5 | 81 |
| HCD628_Winter2021_314T_fMRI_CONCAT_ALL_clean | 22.9 | 77.1 | 24 | 105 |
| HCD628_Winter2021_160independent_fMRI_CONCAT_ALL_clean | 16.3 | 83.7 | 20 | 123 |
| HCD628_Winter2021_100independent_fMRI_CONCAT_ALL_clean | 22.5 | 77.5 | 25 | 111 |
| HCD628_Winter2021_60independent_fMRI_CONCAT_ALL_clean | 38.6 | 61.4 | 44 | 114 |
| S1200_MSMAll3T535V_rfMRI_REST | 0 | 100 | 0 | 78 |
| S1200_MSMAll3T535V_tfMRI_Concat | 0 | 100 | 0 | 67 |
| S1200_MSMAll7T87V_rfMRI_REST_7T | 46.3 | 53.7 | 50 | 108 |
| S1200_MSMAll7T87V_tfMRI_MOVIE_7T | 38 | 62 | 38 | 100 |
| S1200_MSMAll7T87V_tfMRI_RET_7T | 21.9 | 78.1 | 16 | 73 |
| AABC_Version2_Prelim_Data_Visits_617V_fMRI_CONCAT_ALL_clean | 6.4 | 93.6 | 5 | 78 |
| HCD628_Winter2021_314V_fMRI_CONCAT_ALL_clean | 25.7 | 74.3 | 28 | 109 |
| HCD628_Winter2021_60V_fMRI_CONCAT_ALL_rclean | 66.1 | 33.9 | 80 | 121 |
| HCD628_Winter2021_100V_fMRI_CONCAT_ALL_rclean | 46.8 | 53.2 | 52 | 111 |
| HCD628_Winter2021_60V_fMRI_CONCAT_ALL_rclean_initialize78 | 3.8 | 96.2 | 3 | 78 |
| HCD628_Winter2021_100V_fMRI_CONCAT_ALL_rclean_initialize78 | 1.3 | 98.7 | 1 | 78 |

*Supplementary Table 1* ***The summary of single subject related components in each dataset.*** *The datasets with fewer subjects tend to have a relatively larger percentage of single subject dominated components, e.g., HCP-YA 7T datasets, HCP-D with 60, 100, 160 subjects. The INITIALIZE mode (the last two rows) helped to reduce the percentage for the evaluation datasets: HCP-D with 60 and 100 subjects.*

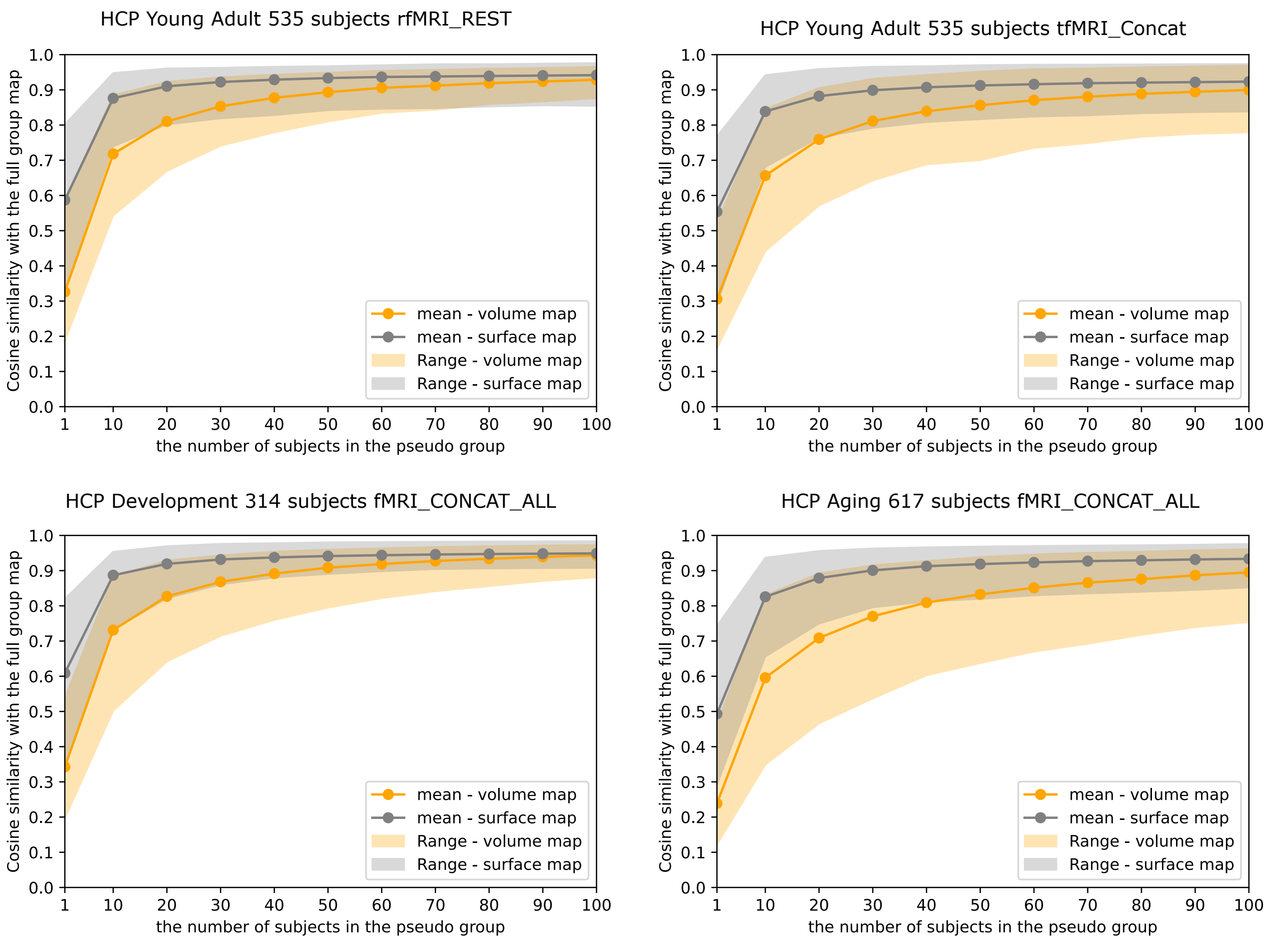


*Supplementary Figure 4* ***Effect of the number of subjects in the pseudo group.*** *The pseudo-group augmented maps have high cosine similarity with the original group maps in the training datasets for both volume (orange) and grayordinate (grey). The individual's grayordinate map has around 50% similarity with the group grayordinate map. 100 subjects make the pseudo map over 85% similar with the group grayordinate map.*

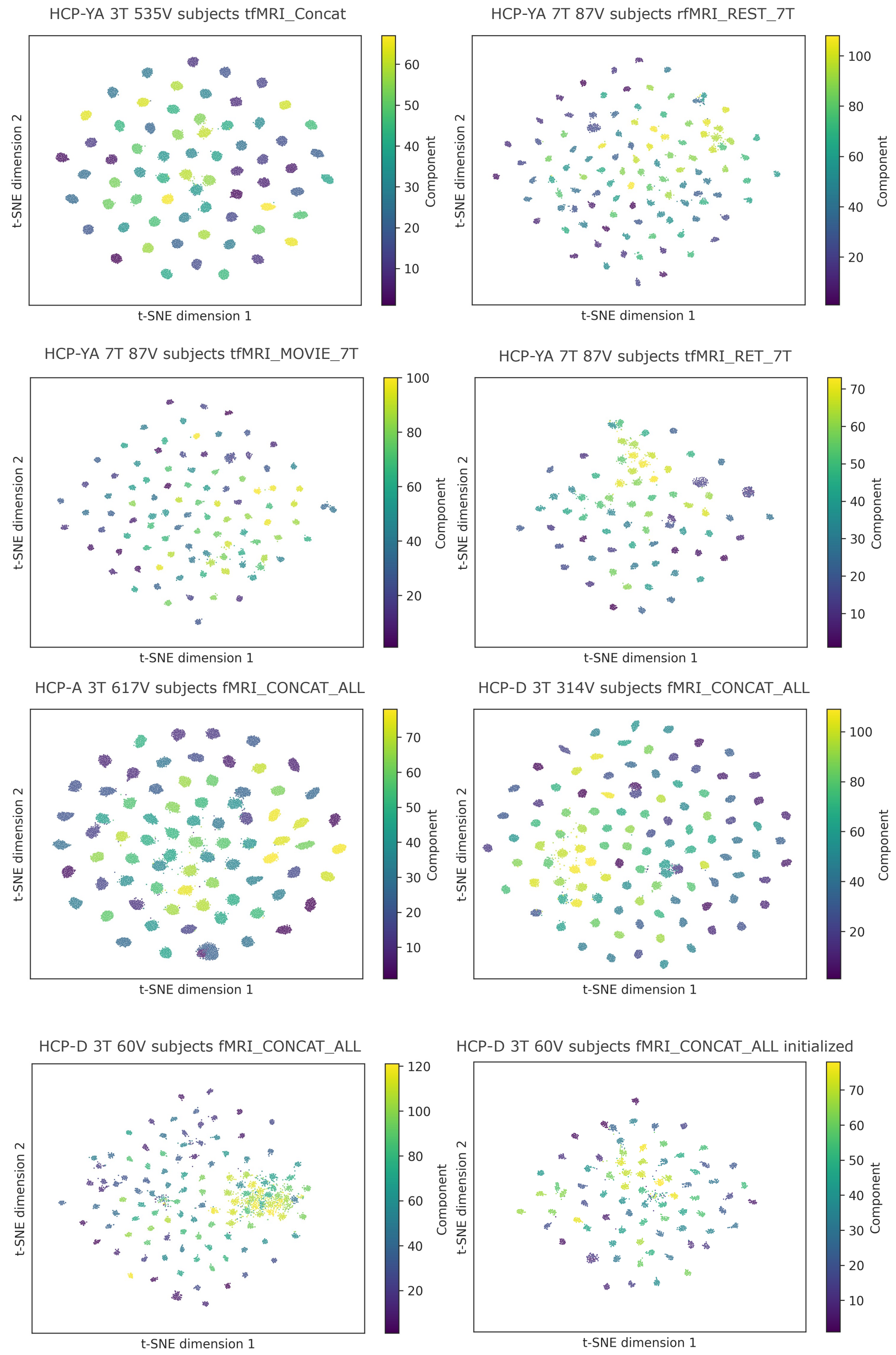


*Supplementary Figure 5* ***Two-dimensional projection of the learned surface representations using t-SNE****. Each point corresponds to one individual tICA component from one subject. Coloring is based on the group tICA component index.*

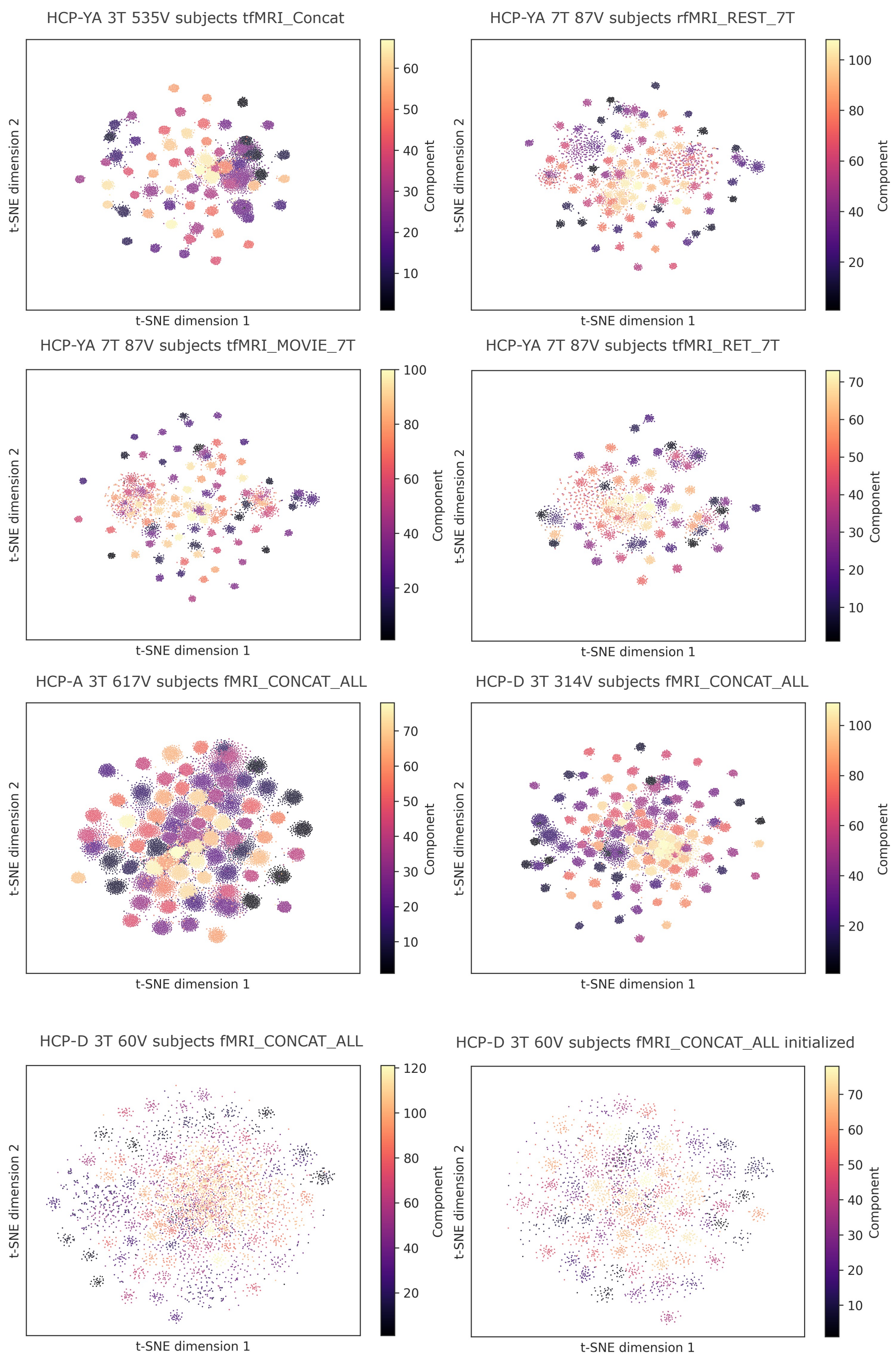


*Supplementary Figure 6* ***Two-dimensional projection of the learned volume representations using t-SNE****. Each point corresponds to one individual tICA component from one subject. Coloring is based on the group tICA component index.*

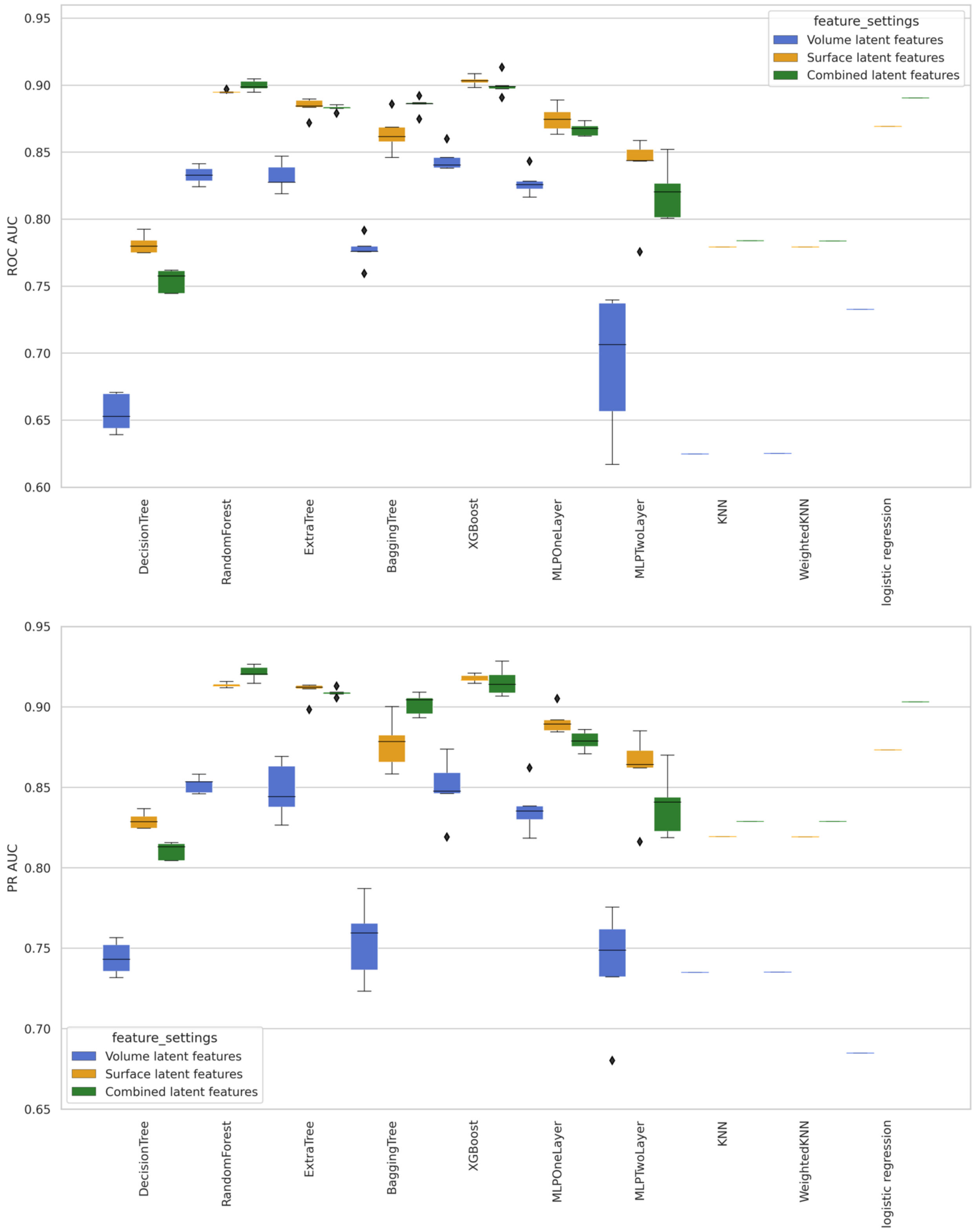
ROC AUC
0.95
0.90
0.85
0.80
0.75
0.70
0.65
0.60
feature_settings
Volume latent features
Surface latent features
Combined latent features
DecisionTree
RandomForest
ExtraTree
BaggingTree
XGBoost
MLPOneLayer
MLPTwoLayer
KNN
WeightedKNN
logistic regression
PR AUC
0.95
0.90
0.85
0.80
0.75
0.70
0.65
feature_settings
Volume latent features
Surface latent features
Combined latent features
DecisionTree
RandomForest
ExtraTree
BaggingTree
XGBoost
MLPOneLayer
MLPTwoLayer
KNN
WeightedKNN
logistic regression

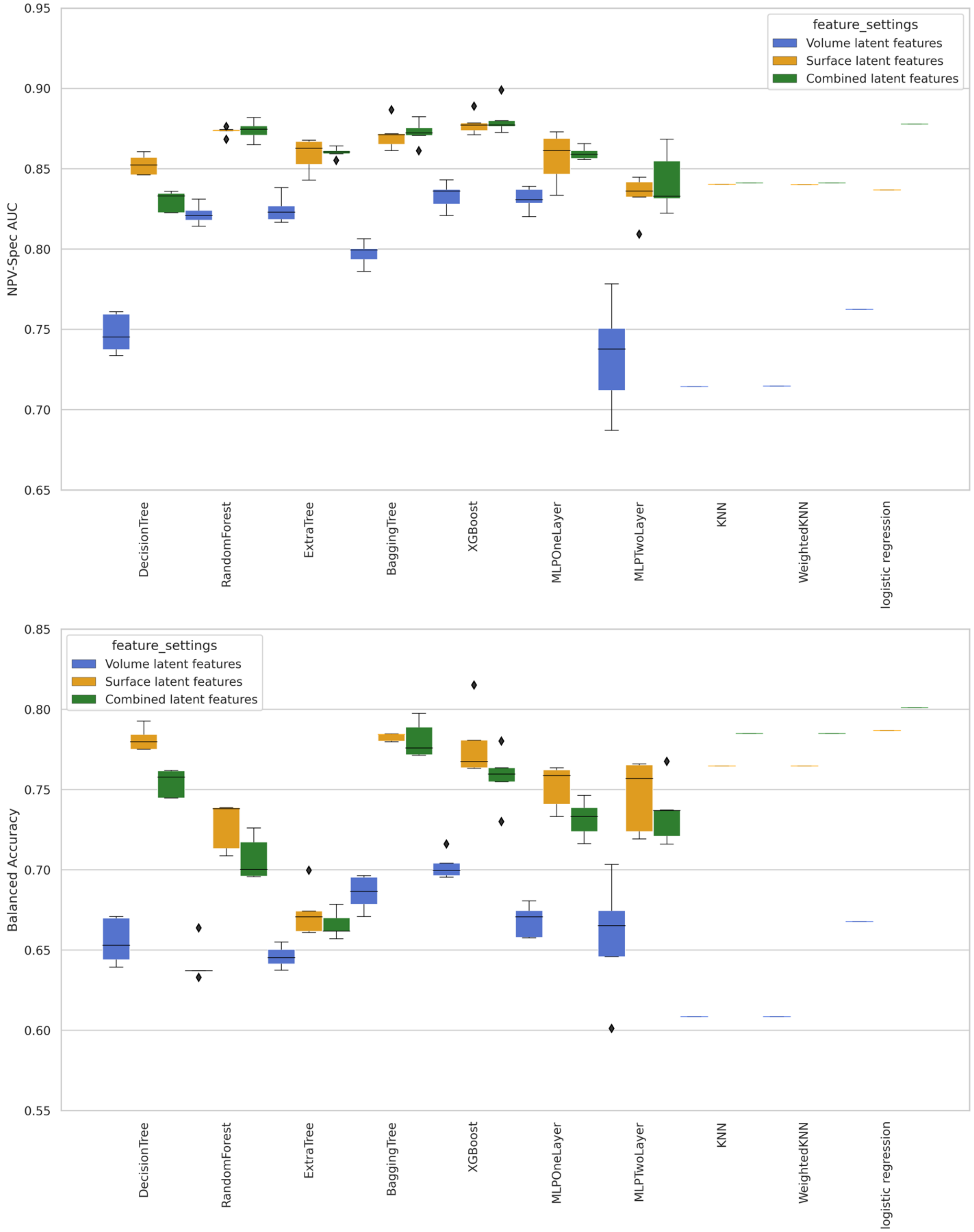


*Supplementary Figure 7* ***Effect of different latent feature settings****. The validation performance between base learners trained on only latent features under different settings, measured by AUC, PR-AUC, NPV-Specificity-AUC, and balanced accuracy. The three types of latent feature settings, volume latent features, surface latent features, and combined latent features are colored blue, orange, and green, respectively. The surface latent features outperform the volume latent features for all the base learners, indicating a more informative representation of spatial patterns.*

*Among the base learners with top five performances, three of them, logistic regression, Random Forest, BaggingTree trained on combined features have better overall performance than those trained on only the surface latent features, suggesting the effectiveness and necessity of combining the features from volume and cortical surfaces.*

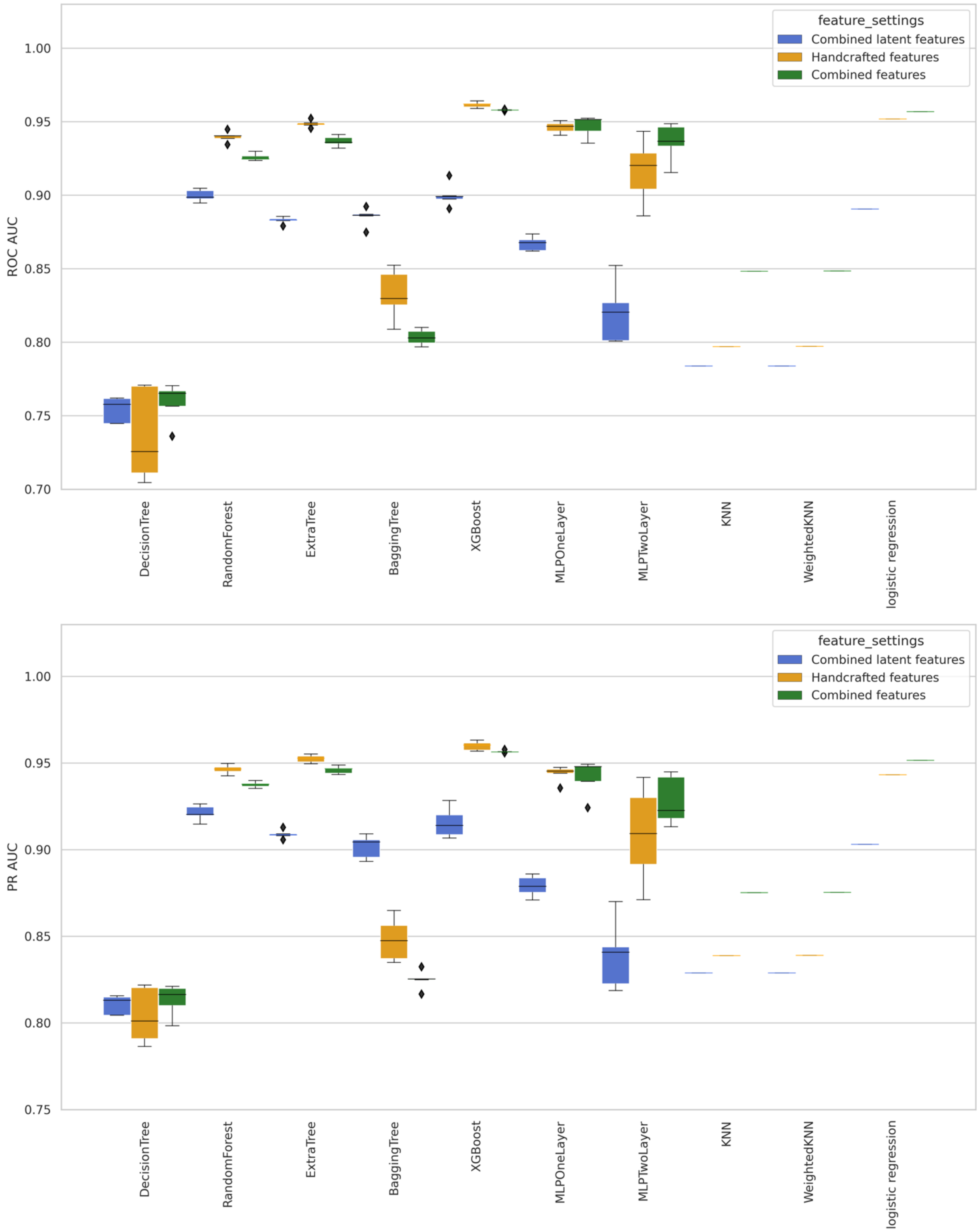

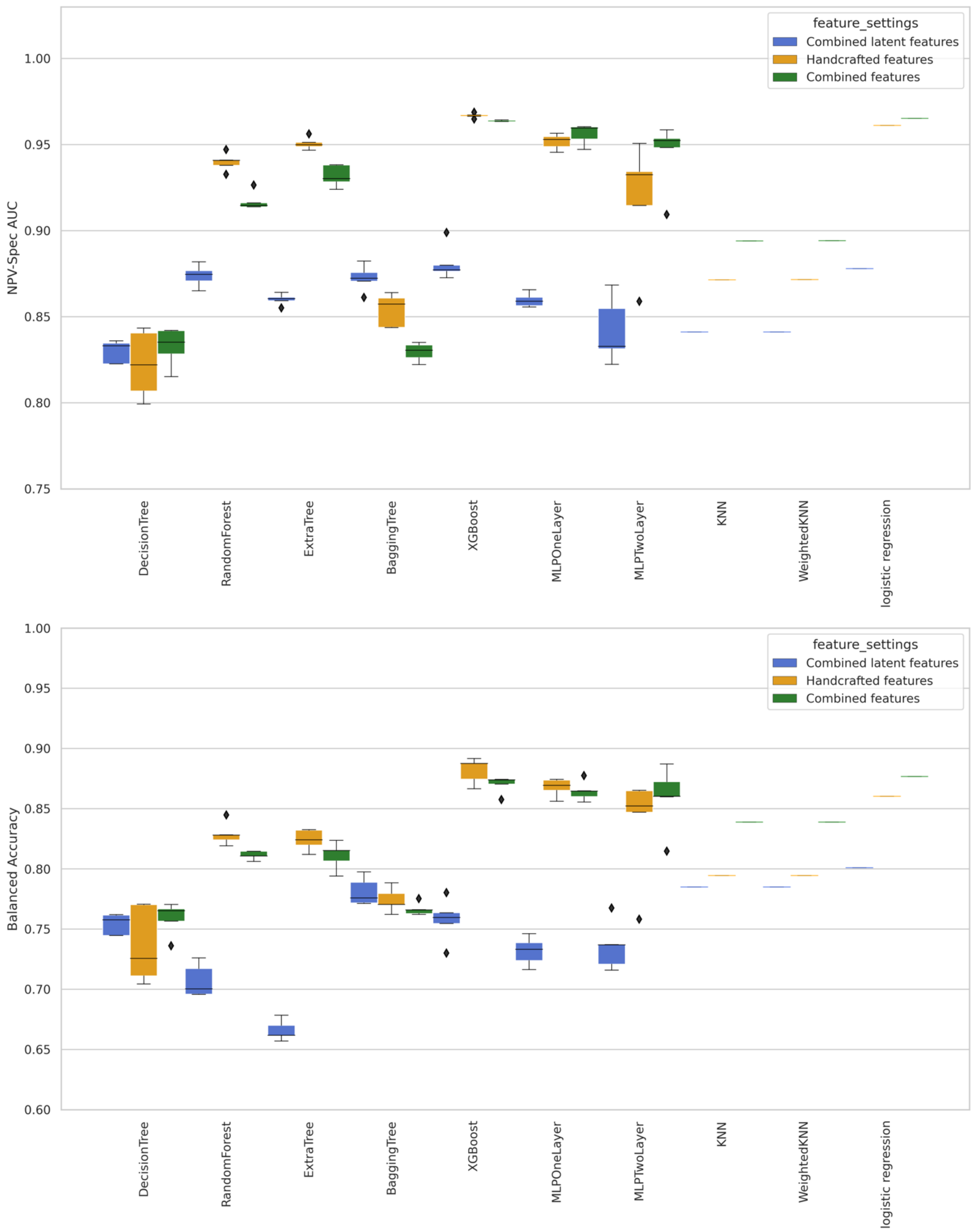


*Supplementary Figure 8* ***Effect of different feature settings****. The validation performance between base learners trained on different feature settings, measured by AUC, PR-AUC, NPV-Specificity-AUC, and balanced accuracy. The three types of features feature settings, combined latent features, handcrafted features and combined features are colored blue, orange, and green, respectively. The handcrafted features have a comparable performance with the*

*combined features, while the latent features were consistently beaten by the handcrafted features. To make sure no useful features are excluded, the combined feature setting is preferred as the final setting.*

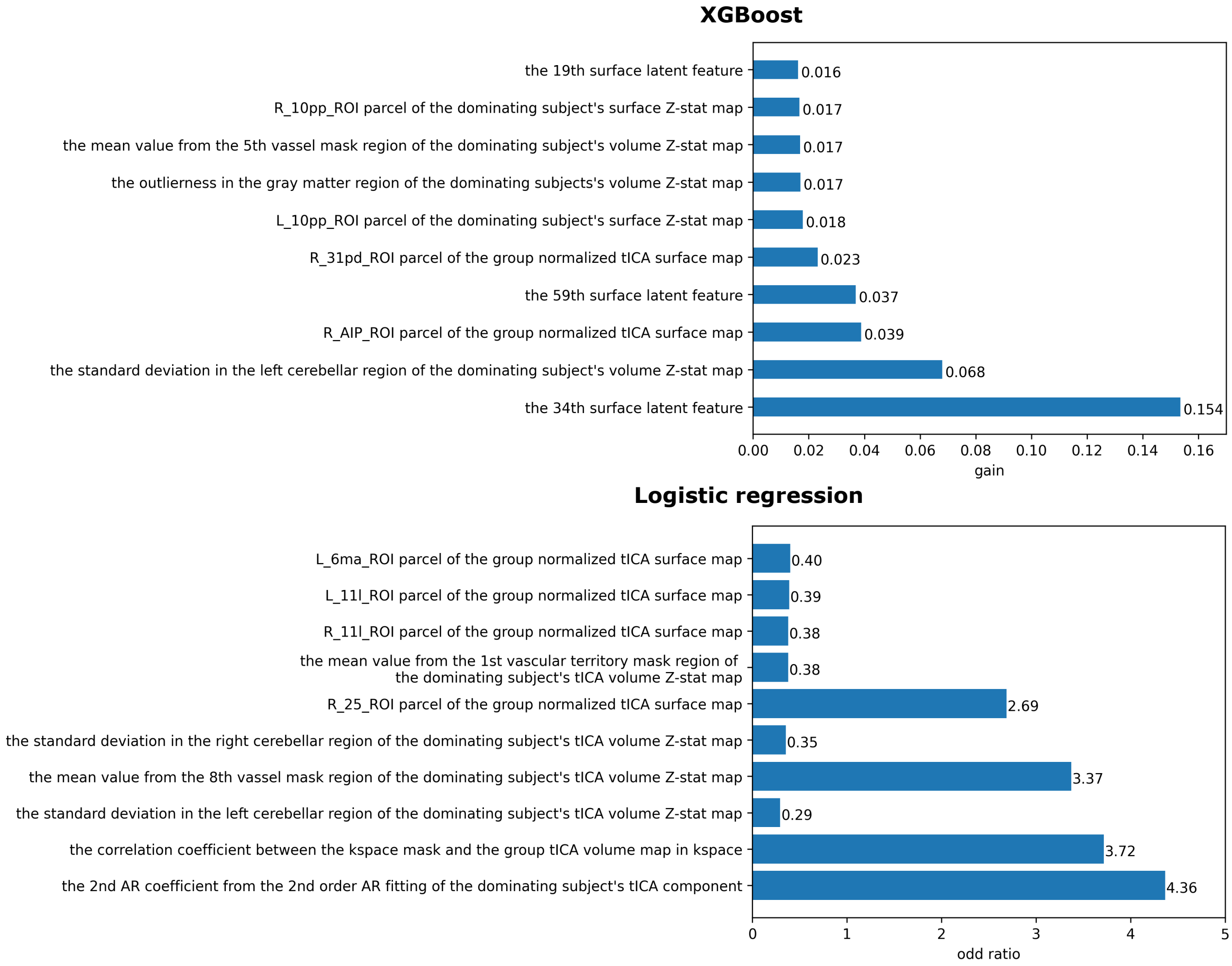


*Supplementary Figure 9* ***The feature importance and interpretability analysis****. The gain and odds ratio are used to measure the feature importance for XGBoost and logistic regression.*

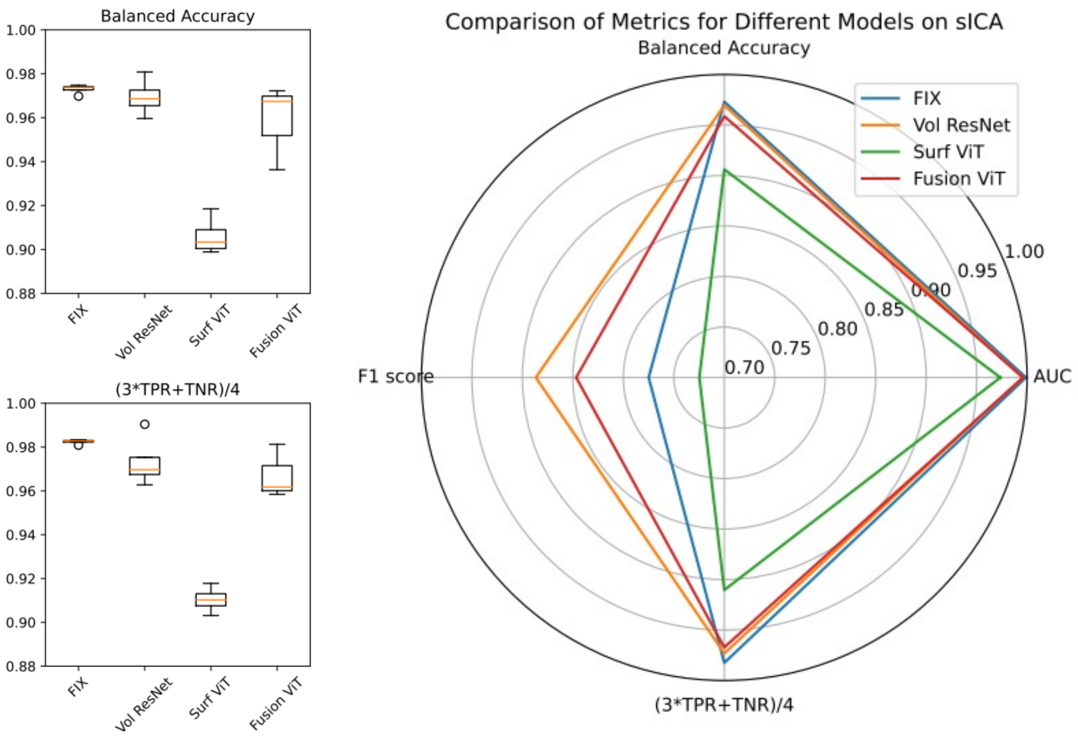


*Supplementary Figure 10* ***The deep learning vs. handcrafted features in spatial ICA classification****. The baseline model is FIX, and the other models are based on deep learning: 1) Vol ResNet: volume-based ResNet; 2) Surf ViT: surface-based vision transformer; 3) Fusion ViT: a fusion vision transformer where volume, surface, timeseries, and power spectrum are divided into patches then concatenated as a sequence of inputs. FIX has the best balanced accuracy and (3*TPR+TNR)/4.*

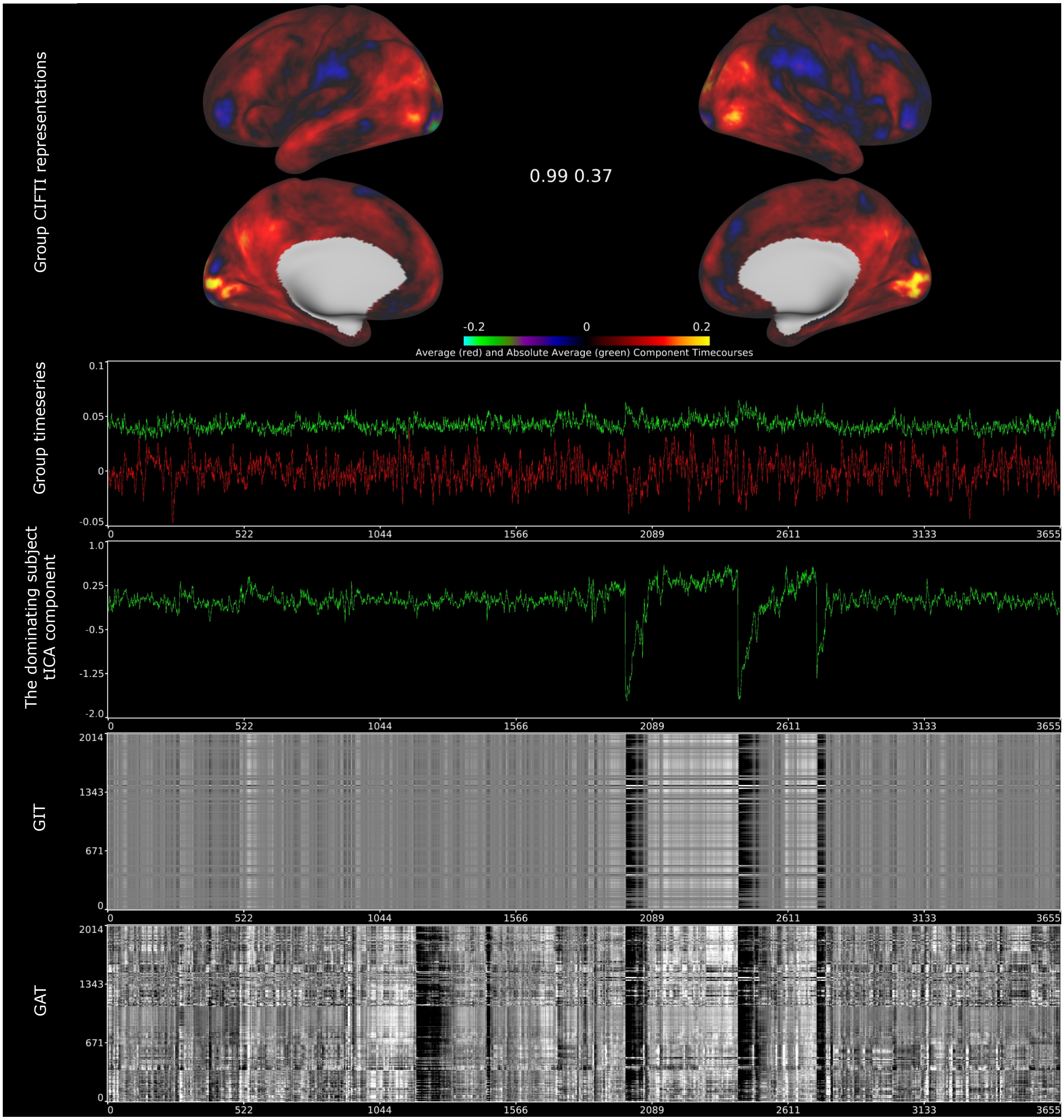


*Supplementary Figure 11* ***An example of a 7T volume distortion component****. The volume distortion component results from severe head motion in HCP 7T datasets leading to incomplete distortion correction and thereby misalignment of greymatter throughout an fMRI run. The extreme value changes can be found in the temporal ICA derived greyplots and component in the third run of subject 186949 from the HCP-YA 7T movie task in the tICA training dataset (87T). However, the group spatial maps exhibit visual signal-like pattern. This contradictory pattern cannot be adequately captured by spatial features alone. Data at https://balsa.wustl.edu/study/gmrqZ.*

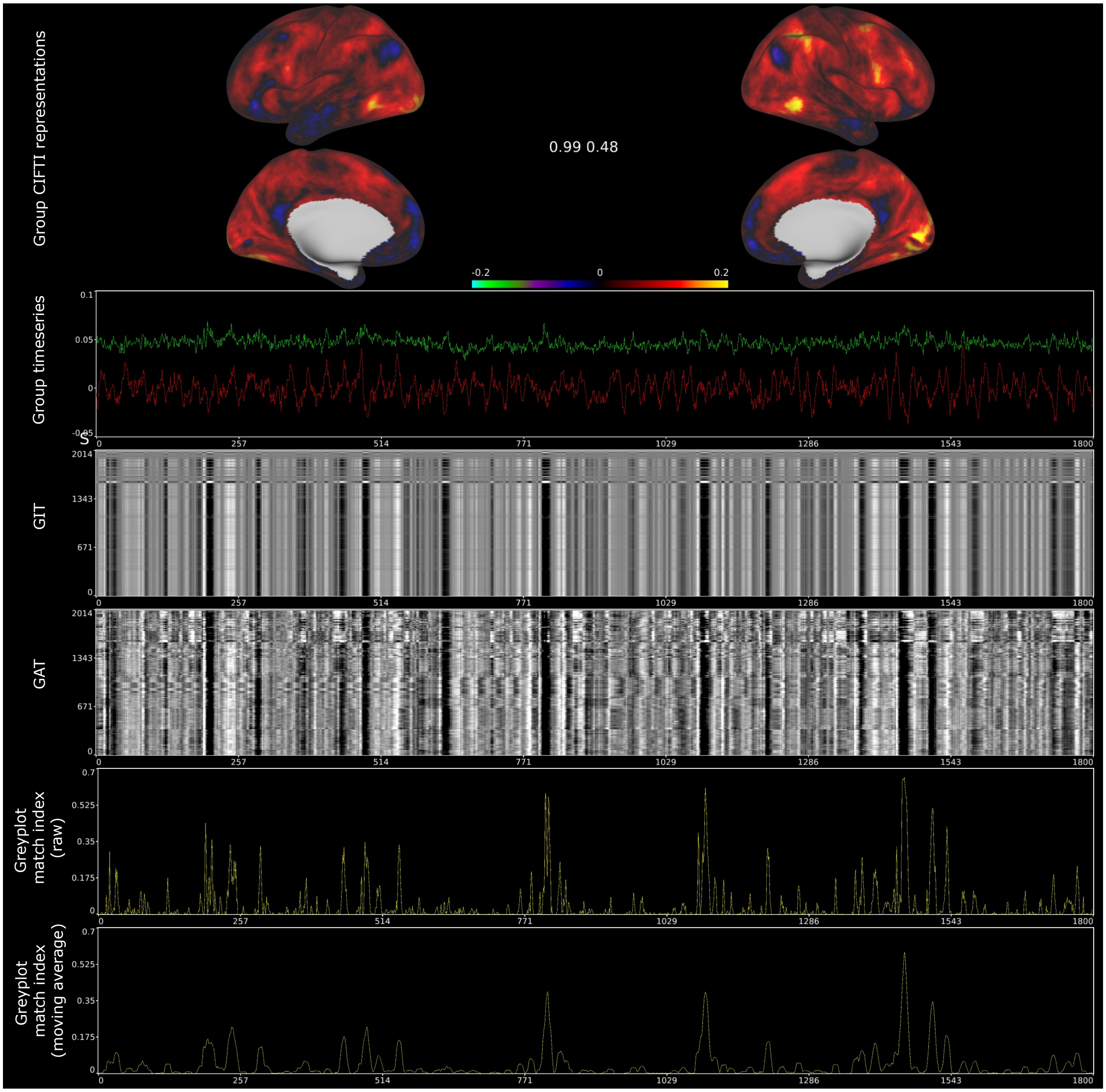


Supplementary Figure 12 ***An example of a single subject respiration component identified by matching the GIT and GAT****. The single subject respiration component is the #71 component from HCP-YA 7T retinotopy task in the tICA evaluation dataset (87V). The global effect can be quantified by matching GIT and GAT timepoint-wise with eta$^2$ multiplied by a rescaled sigmoid function, resulting in non-negative match indices shown in row #5 (raw values) and #6 (moving average with a window size of 11). The global stripes can be located by analyzing the peaks. Data at https://balsa.wustl.edu/study/gmrqZ.*

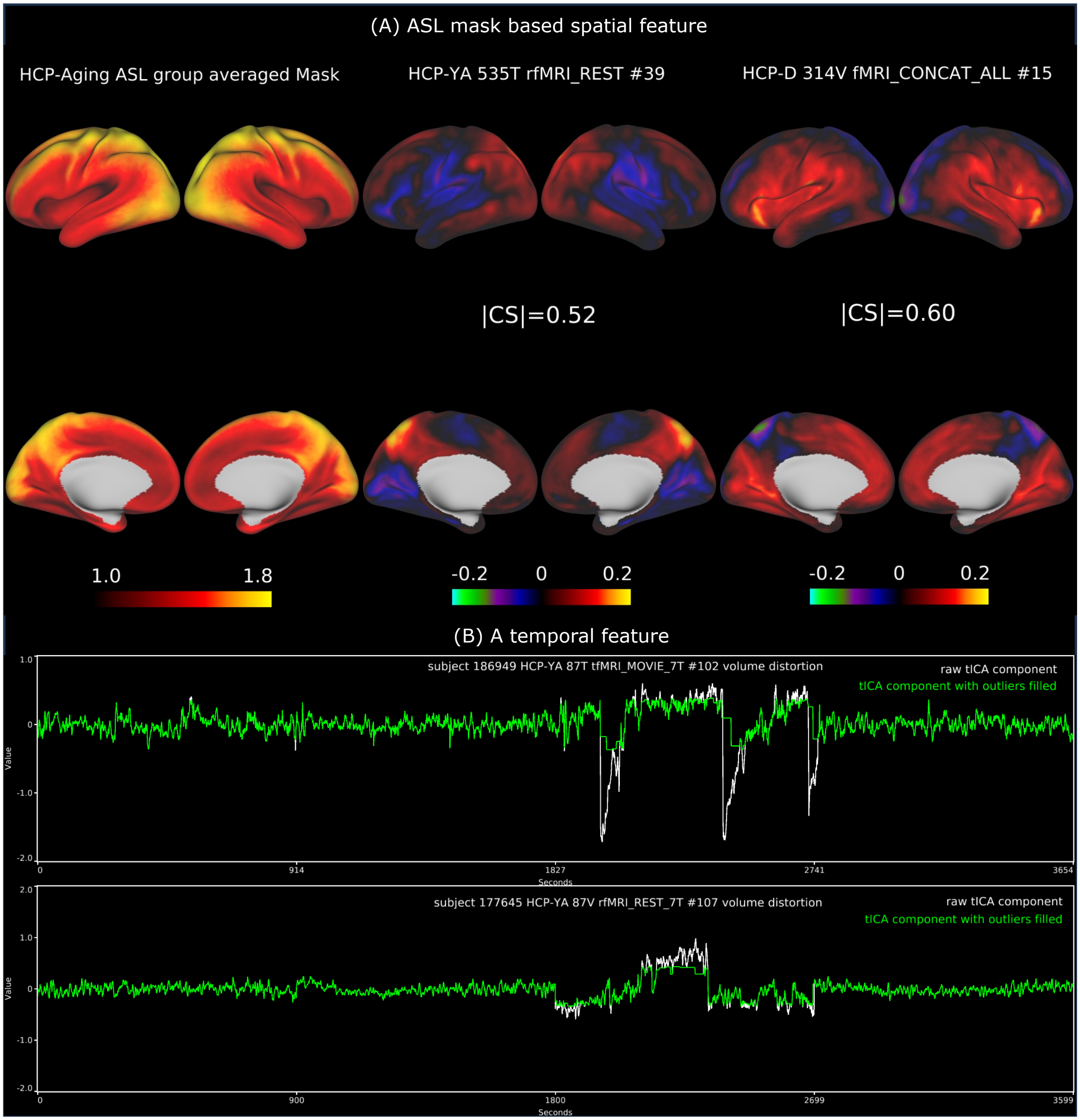


*Supplementary Figure 13* ***Two examples of artifactual components identified by spatial and temporal handcrafted features****. (A) Vascular delay components have high absolute cosine similarity with the ASL group averaged arterial transit time map. (B) Volume distortion components can be identified by computing the square root of the timepoint-wise sum of the differences between the dominating subject's raw tICA component and the component after replacing the outliers with neighbor non-outlier values. Data at https://balsa.wustl.edu/study/gmrqZ.*

**Supplementary Methods**

<u>The semi-automated sICA reclean process for the official release of HCP-YA and HCP-Lifespan</u>
The reclean process is designed to identify incorrectly classified sICA components using sICA+FIX (Salimi-Khorshidi et al., 2014). This is achieved by leveraging the existing large-scale partially hand-labeled (all predicted with FIX probabilities) sICA components from HCP-style datasets, along with additional handcrafted temporal and spatial features. The first two features are the percentage of explained variance and total variance of the sICA component.

The newly introduced temporal features consist of various metrics, including power greater than 0.25 Hz divided by the power between 0.005 Hz and 0.1 Hz, power smaller than 0.005 Hz divided by the power between 0.005 Hz and 0.1 Hz, and the amplitude of noisy timepoints divided by the amplitude of all timepoints, with or without weighting by explained variance.

The newly introduced spatial features include the sum of the absolute and mean absolute ratios between 1) CIFTI and non-greymatter region, 2) CIFTI and the non-greymatter plus the edge voxels, 3) cerebral cortex and non-greymatter, 4) cerebral cortex and non-greymatter plus the edge voxels, 5) the 4mm FWHM surface and tissue-constrained volume smoothed versions of 1 to 4. 6) CIFTI and volume, 7) cortex and volume, 8) subcortical and volume, 9) cerebellum and volume, 10) brainstem and volume, 11) diencephalon and volume, 12) left cortex and right cortex, 13) 2-voxel eroded white matter and volume, 14) 2-voxel eroded CSF and volume, 15) outer edge CSF and volume, 16) volume dropout region and volume, 17) CIFTI dropout region and CIFTI, 18) visual ROI CIFTI and non-visial ROI CIFTI, 19) language ROI CIFTI and non-language ROI CIFTI and 20) visual ROI CIFTI and cerebellum CIFTI. Additionally, we include the sum of absolute values of the 360 cortical areas from CIFTI using HCP-MMP1.0, and 10 sensory-motor subareas.

Furthermore, a set of features based on group sICA is included, with the number of features varying across different large HCP datasets. This set consists of 1) 4mm FWHM smoothed CIFTI absolute correlation coefficients with group sICA spatial maps of the same dataset, 2) power spectral correlations with group sICA power spectra and 3) absolute temporal correlations with group sICA average timeseries.

For HCP-YA 3T and 7T datasets, the framework for reclassifying sICA components involves four key steps:

1. Two MLP classifiers are trained on sICA components using different thresholds for the initial FIX probabilities, a) components predicted as signal with over 0.99 probabilities and components predicted as artifacts with less than 0.5 probabilities; b) components predicted as signal with over 0.5 probabilities and components predicted as artifacts with less than 0.01 probabilities. Once trained, both classifiers are used to infer probabilities for all components.

2. Another MLP classifier is trained on components with a probability difference of less than 0.1 between the results obtained from the two MLP classifiers in step 1. A decision threshold of 0.5 is applied to classify components as signal or artifacts.
3. sICA components with similar feature vectors are clustered assigned with the probabilities computed from the weighted average of similar components where the similarity is accessed by $eta^2$ (Cohen et al., 2008). The minimum cluster size is set at 6 components, and the maximum cluster size is limited to 2% of the total number of components. Cluster boundaries are determined by identifying local maxima in the derivatives of sorted $eta^2$ values. If the mean probability falls outside the range of 0.33 to 0.66, it is extended to the next most similar group of components. If no cluster meets the specified thresholds, the one that differs the most from 0.5 is chosen.
4. Finally, two additional rounds of MLP classifiers are trained on all components using a threshold of 0.5 for signal and nuisance classification. Components with reclassified categories compared to the FIX predictions are manually reviewed.

This framework was validated on the HCP 7T retinotopy sICA dataset (47,985 components), which was manually labelled by CY and the uncertain labels by CY are further confirmed by MFG. The accuracy of the sICA component classification increased from 96.1% to 99.25%. The same semi-automated recleaning process was subsequently applied to all other HCP-YA 3T and 7T datasets.

When it comes to the HCP-A and HCP-D datasets, the recleaning process is less complex, owing to the substantial number of semi-labeled sICA components available from the HCP-YA 3T and 7T datasets.

For the HCP-A recleaning process, a pair of classifiers is employed—a two-layer Multi-Layer Perceptron (MLP) and a Random Forest classifier. These classifiers are trained on all the HCP-YA 3T datasets, including both 3T resting state and 3T task fMRI data. Subsequently, they are validated on the HCP-YA 7T datasets, which encompass 7T resting state and 7T movie data. The final evaluation is conducted on the manually labeled 7T retinotopy dataset. To ensure robustness, this process is repeated five times with different random seeds, resulting in ten sICA classifiers trained on large-scale sICA component datasets. These ten classifiers are then used to make predictions on the HCA datasets, and any disagreements between the predicted sICA component categories from these classifiers and the FIX predictions are flagged for manual classification.

In the case of the HCP-D recleaning process, a similar approach is adopted. However, the models are trained on the recleaned HCP-A dataset due to their closely matched acquisition settings. Subsequently, these models are validated using the HCP-YA 3T datasets, and any disagreements compared with FIX predictions are manually inspected.

The MLP was implemented using the PyTorch framework with 9 hidden nodes, emulating the architecture of the areal classifier described in (Glasser et al., 2016). We employed a weighted cross-entropy loss function and guided the optimization process using the Adam optimizer. A learning rate of 1e-3 was initially chosen and gradually reduced by a factor of 0.3 when the validation loss did not increase for five consecutive epochs. We set a maximum of 100 epochs for training. The random forest classifier was implemented using the scikit-learn library (Pedregosa et al., 2011), and class-balanced weight terms were applied to considering the imbalanced datasets.

Automated sICA reclean pipeline

In the semi-automated approach for the official release of HCP-YA and HCP-Lifespan datasets, base learners like MLP or random forest have been explored (See section *Semi-automated sICA reclean process for the official release of HCP-YA and HCP-Lifespan*). These base learners have demonstrated the ability to effectively train on large-scale sICA components while achieving excellent performance. Hence, we have designed an automated sICA reclean pipeline, as illustrated in Fig S14, utilizing the extensive sICA components sourced from multiple HCP-style datasets with partially hand-labeled labels. This pipeline leverages lightweight base learners, such as XGBoost, Random Forest, and MLP.

We conducted an experiment to compare the performance of FIX (the version for the official release of HCP-YA and HCP-Lifespan), retrained FIX, and several base learners using the sICA components from HCP-Aging datasets. These datasets consist of 1798 sessions, which were divided into a training dataset (1438 sessions; 453,696 components) and an evaluation dataset (360 sessions; 112,632 components). The classification performance is presented in Table S2, and the training time and the size of trained model weight are detailed in Table S3. The results indicate that the lightweight base learners employed in the sICA reclean pipeline deliver comparable performance to FIX, yet require substantially less time for retraining and result in substantially smaller model weights.

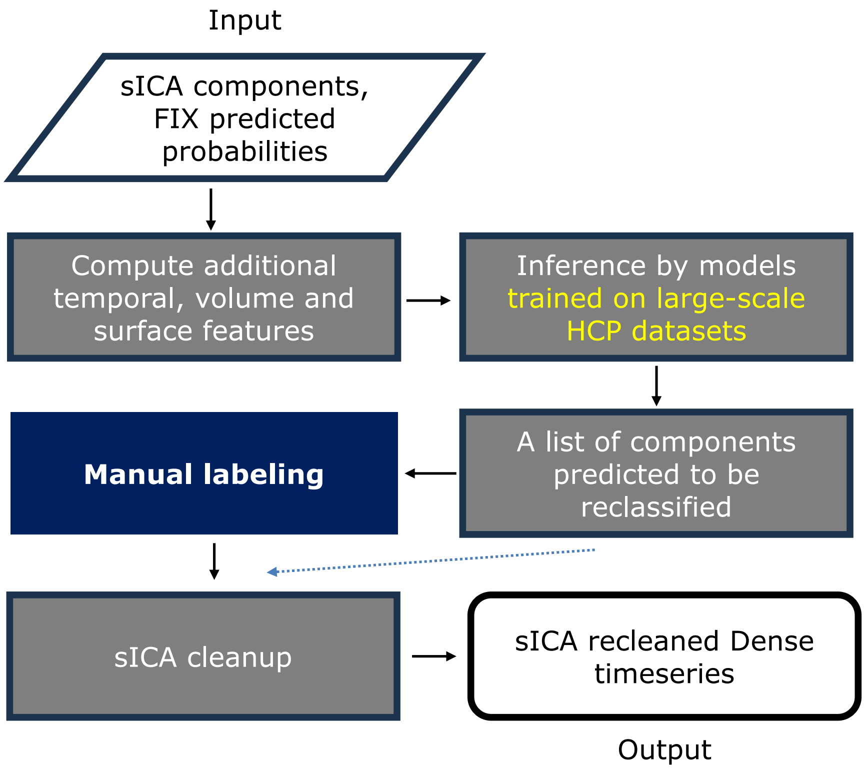


*Supplementary Figure 14* ***The automated sICA reclean pipeline***.

| model | F1 score | AUC | Balanced Accuracy | PRAUC |
|---|---|---|---|---|
| FIX | 0.974 | 0.99840 | 0.9754 | 0.9959 |
| FIX retrain | 0.984 | 0.99980 | 0.9905 | 0.9993 |
| MLP | 0.981 | 0.99932 | 0.9910 | 0.9972 |
| RandomForest | 0.981 | 0.99983 | 0.9846 | 0.9986 |
| WeightedKNN | 0.975 | 0.98913 | 0.9785 | 0.9883 |
| Xgboost | 0.995 | 0.99960 | 0.9909 | 0.9980 |

*Supplementary Table 2* ***The summary of classification performance on the HCP-Aging dataset.*** *The FIX used for the official release of HCP-YA 3.0 and HCP-Lifespan 3.0 has a worse performance than the retrained FIX. The base learners have comparable performance with the retrained FIX, with some metrics outperforming.*

| model | Training time (hr) | Trained weight size (GB) |
|---|---|---|
| FIX | 325.7 | 15 |
| MLP | 0.4 | 3E-4 |
| RandomForest | 0.1 | 0.02 |
| WeightedKNN | 0.0018 | 2.6 |
| XGBoost | 0.03 | 3E-4 |

*Supplementary Table 3* ***The summary of training time and size of trained models.*** *FIX requires a training time about 14 days while the lightweight base learners only need less than an hour.*
